# The length of haplotype blocks and signals of structural variation in reconstructed genealogies

**DOI:** 10.1101/2023.07.11.548567

**Authors:** Anastasia Ignatieva, Martina Favero, Jere Koskela, Jaromir Sant, Simon R. Myers

## Abstract

Recent breakthroughs have enabled the accurate inference of large-scale genealogies. Through modelling the impact of recombination on the correlation structure between genealogical local trees, we evaluate how this structure is reconstructed by leading approaches. Despite identifying pervasive biases, we show that applying a simple correction recovers the desired distributions for one algorithm, Relate. We develop a statistical test to identify clades spanning unexpectedly long genomic regions, likely reflecting regional suppression of recombination in some individuals. Our approach allows a systematic scan for inter-individual recombination rate variation at an intermediate scale, between genome-wide differences and individual hotspots. Using genealogies reconstructed with Relate for 2 504 human genomes, we identify 50 regions possessing clades with unexpectedly long genomic spans (*p <* 1 · 10^−12^). The strongest signal corresponds to a known inversion on chromosome 17. The second strongest uncovers a novel 760kb inversion on chromosome 10, common (21%) in S. Asians and correlated with GWAS hits for a range of phenotypes. Other regions indicate additional genomic rearrangements: inversions (8), copy number changes (2), or other variants (12). The remaining regions appear to reflect recombination suppression by previously unevidenced mechanisms. They are enriched for precisely spanning single genes (*p* = 5 · 10^−10^), specifically those expressed in male gametogenesis, and for eQTLs (*p* = 2 ·10^−3^). This suggests an extension of previously hypothesised crossover suppression within meiotic genes, towards a model of suppression varying across individuals with different expression levels. Our methods can be readily applied to other species, showing that genealogies offer previously untapped potential to study structural variation and other phenomena impacting evolution.

## 1 Introduction

In the presence of recombination, the genealogical history of a sample can be fully captured in the form of an ancestral recombination graph (ARG). This can be represented as a sequence of local trees describing the sample genealogy at each locus, connected by recombination events that reshape these trees along the genome. ARGs can, in principle, capture the effects of all the evolutionary forces that have shaped the observed genetic diversity of a sample of sequences, while providing a much more efficient representation of genomes than multiple sequence alignments (Kelleher et al., 2019). Thus, statistical inference methods which take ARGs as inputs have the potential to provide very powerful insights into evolutionary events and parameters.

However, the true underlying genealogy is usually not observable in practice, and the ARG must be reconstructed from the data, which typically comprises a set of genetic sequences sampled at the present time. This is a notoriously difficult problem, due to the computational cost of traversing the huge search space of plausible ARGs. This has been the main bottleneck for the widespread development of genealogy-based inference, but has seen impressive recent breakthroughs, with methods now capable of reconstructing and efficiently storing ARGs for tens or even hundreds of thousands of samples (Speidel et al., 2019; Wohns et al., 2022; Zhang et al., 2023; Zhan et al., 2023). Inference using ARGs reconstructed from large-scale human sequencing datasets is in its infancy, but has already produced novel scientific insights, for instance in untangling the history of human demography (Speidel et al., 2019; Wohns et al., 2022) and understanding the phenotypic effects of genetic variants (Zhang et al., 2023).

This is a rapidly developing field, however progress has been hampered by the fact that methods which work very well on simulated ARGs often lose power and accuracy when applied to reconstructed genealogies, for reasons that are, in general, poorly understood. Methods for quantifying the quality of ARG reconstruction have generally been limited to comparing simulated to reconstructed ARGs, to quantify how well local tree topology (Speidel et al., 2019; Kelleher et al., 2019) and pairwise coalescence times (Brandt et al., 2022) are recovered. Deng et al. (2021) and McKenzie and Eaton (2022) took the approach of deriving the distribution of the genomic distance between consecutive local trees (under a given model), and comparing this to the empirical distributions calculated from reconstructed ARGs. All of these studies have broadly demonstrated that different tools have somewhat different strengths, but since they commonly output strikingly different ARGs for the same dataset (Wong et al., 2024), more in-depth exploration is needed to understand (and correct) the underlying causes.

Moreover, all currently available methods of recording and inferring ARGs typically only account for mutations in the form of single base substitutions, and ignore the presence of genomic structural variants (SVs), such as duplications and inversions. SVs are a ubiquitous and evolutionarily important type of mutation, playing a key role in speciation and local adaptation (Kirkpatrick and Barton, 2006), and within human populations through altering the structure and expression of genes (Chiang et al., 2017; Abel et al., 2020). Thus, identifying and analysing the evolution of structural variants at all scales is an incredibly important goal, but so far no studies have attempted to leverage ARGs for this purpose, beyond simulation (e.g. Peischl et al., 2013).

Within an ARG, each edge has a well-defined genomic span during which it is present in the local trees (Figure 1). We analytically derive the theoretical distribution of this quantity using the well-known SMC’ model (Marjoram and Wall, 2006), which is an excellent approximation to the coalescent with recombination (Griffiths and Marjoram, 1997). Through calculating the empirical distribution of edge span in ARGs reconstructed by ARGweaver, Relate, tsinfer and tsdate, and ARG-Needle in simulation studies, we find that these tools recover the correct distribution to varying degrees of success.

**Figure 1:**
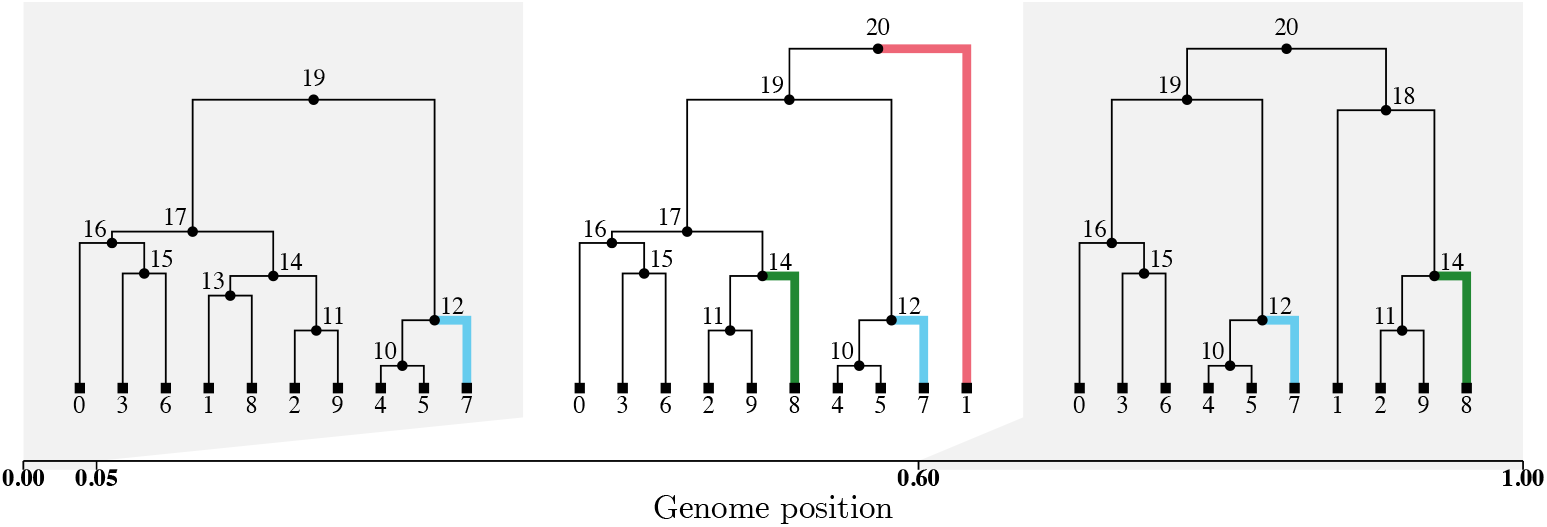
An ARG with *n* = 10 sequences, represented as a sequence of the corresponding local trees along the genome. The *time-length* of an edge is given by the time of its parent node less the time of its child node; *older* edges are those closer to the root of a local tree. Edge highlighted in red (1→ 20) spans one local tree and has span 0.55; edge highlighted in green (8 → 14) spans two local trees and has span 0.95; edge highlighted in blue (7 → 12) spans all three local trees and has span 1. ARG simulated using msprime and visualised using tskit (Baumdicker et al., 2022; Kelleher et al., 2016).

We then derive the distribution of the length of a haplotype block within an ARG, defined as the genomic span of a given clade of samples (as by Shipilina et al., 2023). Recombination is suppressed in individuals heterozygous for an inversion, which manifests in the ARG as a clade of samples that persists for a longer stretch of the genome than would otherwise be expected: we use this idea to construct a computational tool for detecting localised (between-clade) suppression of recombination (DoLoReS: Detection of Localised Recombination Suppression). This tool is tailored for use with Relate ARGs, implementing suitable adjustments to correct for possible sequencing and phasing errors in the data, and the particularities of the ARG reconstruction method. We demonstrate the power and accuracy of this tool for both simulated and reconstructed ARGs.

Finally, we apply DoLoReS to an ARG for the 1000 Genomes Project (1KGP; 1000 Genomes Project Consortium, 2015) reconstructed using Relate (Speidel et al., 2019). We detect several known inversions: for instance, one of the top significant hits is the 17q21.31 inversion polymorphism common in European populations (Stefansson et al., 2005). The second strongest signal corresponds to a previously unknown 760kb inversion on 10q22.3, which we validate using data from the Human Pangenome Reference Consortium (HPRC; Liao et al., 2023); this inversion is common in S. Asian populations (with a frequency of 21%), spans a number of genes associated with lung function, and correlates with a number of GWAS hits for hematological and immunological traits. We find several other new SVs, and show that our method also detects distinguishable signals of other structural variants, particularly copy number variants (CNVs) and complex rearrangements. This demonstrates that, while Relate only uses SNP data and does not explicitly infer or account for SVs, the reconstructed ARGs still capture the signal of SV presence.

Our work thus presents new results on the SMC’ model by characterising the distribution of genomic spans of edges and clades, and also demonstrates that these are very close to the equivalent distributions under the coalescent with recombination. This adds to earlier work demonstrating the quality of the SMC’ approximation (Wilton et al., 2015; Hobolth and Jensen, 2014) and using it to derive various quantities and distributions of interest (Eriksson et al., 2009; Harris and Nielsen, 2013; Carmi et al., 2014; Deng et al., 2021; McKenzie and Eaton, 2022). From the point of view of detecting inversions, (Bansal et al., 2007) looked for SNPs in long-range LD, and our work builds on this by constructing a genealogy-based statistical test for whether such LD is more extreme than expected. This also links to other work focused on detecting inversions through disruptions in LD patterns, such as Kemppainen et al. (2015) and Li and Ralph (2019). Finally, we note the connections to the work of (Peischl et al., 2013), who modelled inversions under the SMC by modifying the rates at which lineages designated as carriers and non-carriers can coalesce in local trees (and analysed this model through simulation).

Code implementing the methods and used to produce the figures is publicly available at github. com/a-ignatieva/dolores.

## 2 Results

### 2.1 The probability that an edge is broken up by the next recombination event along the genome is biased in reconstructed ARGs, particularly for old edges

For a given edge of the ARG, its span can be defined as the genomic positions where it is present in the corresponding local trees (as illustrated in Figure 1); this is determined by recombination events that change local tree topologies and coalescence times. Intuitively, the longer (resp. shorter) the *time-length* of an edge, the more (resp. less) likely it is to be broken up by recombination as we move along the genome, so its *span* along the genome should be shorter (resp. longer). This is a fundamental property of the ARG which has several implications. For instance, in the absence of recombination, the genealogy is a single tree with all edge spans equal to the length of the genome, and the probability that a given edge carries at least one mutation is proportional to its time-length (assuming mutations occur as a Poisson process along the edges). On the other hand, in the presence of recombination, the number of mutations per edge is less variable, because of the interplay between edge time-length and span.

The SMC’ is an approximation to the coalescent with recombination (CwR) model characterising the distribution over sample genealogies. Under this model, an ARG can be simulated by starting with an initial binary tree𝒯_1_, representing the sample genealogy at the leftmost point of the chromosome. The distance until the next recombination breakpoint along the genome is then drawn from an exponential distribution, with rate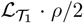, where *ρ* is the population-scaled recombination rate, and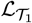 is the total branch length of𝒯_1_. Then a recombination point is chosen uniformly at random along the edges of 𝒯_1_, the subtree underneath this point is allowed to coalesce at a new location (with the usual coalescent dynamics), and the result is the next local tree 𝒯_2_. This process is repeated until the end of the chromosome is reached (see SI, Sections S1.1 and S1.2, for full details).

We first calculate analytically, under the SMC’, the probability ℙ_*𝒯*_ (*b* disrupted) that a given edge *b* in a local tree 𝒯 is disrupted by the next recombination event along the genome (Methods, Section 4.1). This probability does not have a simple form, but can be calculated exactly for a given edge and local tree. The left panel of Figure 2 shows ℙ_*𝒯*_ (*b* disrupted) calculated for each edge in an ARG simulated under the SMC’, against the normalised age of the edge. Middle and right panels show the same quantities, but calculated for each edge of an ARG reconstructed from the simulated data: using Relate (middle), and tsinfer/tsdate (right).

**Figure 2:**
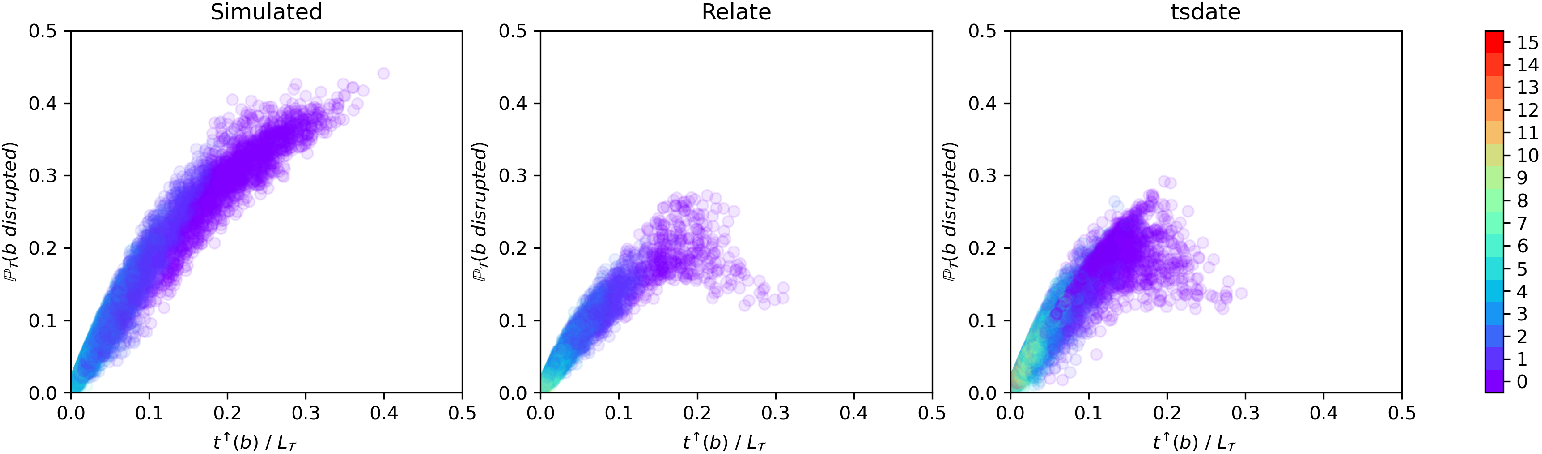
Value of ℙ_𝒯_ (*b* disrupted) calculated for each edge *b*, against the time at its upper endpoint *t*^↑^(*b*) divided by the total branch length of the tree, ℒ_*T*_. Left panel: for a uniform random sample of 10 000 edges in one simulated ARG with *n* = 100 samples (dataset 1 parameters given in Methods, Section 4.6.1). Middle and right: same quantity calculated for each edge of the ARG reconstructed from the simulated data using Relate (middle) and tsinfer/tsdate (right). Colour shows number of edges between the top end of the edge and the root of 𝒯 (purple dots correspond to edges extending from the MRCA node).

For the simulated ARG, the probability that a given edge is disrupted by a recombination event is higher for older edges. This is as expected, as edges at the top of a local tree tend to have greater time-length and co-exist with fewer other lineages. This makes such edges more likely to be quickly broken up by recombination, since (1) the rate of recombination on the edge is relatively higher, due to its greater time-length, and (2) if a recombination event happens below the edge, and the recombinant lineage has not yet coalesced by the time at the lower end of the edge, it is likely to disrupt the edge since it is one of the few remaining choices for coalescence.

However, old edges are generally difficult to accurately reconstruct using the sequences at the leaves due to lack of signal, unless they are strongly supported by mutations (which implies longer edge span along the genome). As a result, reconstructed ARGs have fewer old edges (with fewer points lying towards the right of the plots), and the old edges that *are* present tend to have lower probability of disruption than those of a similar age in the simulated ARG. This bias is seen for both Relate and tsinfer/tsdate. This demonstrates explicitly the difficulty with faithfully reconstructing the ARG topology and event times in the deep past.

### 2.2 The distribution of edge span is recovered with varying accuracy by different ARG reconstruction methods

We next derive an approximation to the distribution of edge span along the genome. If an edge *b* first appears at a position of the genome where the corresponding local tree is 𝒯, we show that the waiting distance along the genome until is it broken up by a recombination event is approximately exponentially distributed as

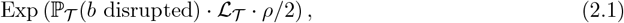

we also derive an equivalent approximation for the case where the recombination rate varies along the genome (Methods, Section 4.2). Given a simulated or reconstructed ARG, we can thus calculate a corresponding *p*-value for each edge under this model (assuming edges have independent exponentially distributed spans), and check if the *p*-values follow the expected (uniform) distribution using a Q-Q plot (SI, Section S1.8).

We apply this to ARGs reconstructed from simulated data using Relate, tsinfer/tsdate, ARG-Needle, and ARGweaver. The resulting Q-Q plots are shown in Figure 3 (the corresponding histograms are also shown in SI, Figure S13). For ARGs reconstructed using Relate and tsinfer/tsdate, we adjust for the fact that these tools do not detect recombination events that affect only edge time-lengths (SI, Section S1.8.1). Moreover, we adjust for the fact that Relate does not attempt to infer edge span when the edge is not supported by at least one mutation (SI, Section S1.8.2), in which case the local topology is resampled from one local tree to the next (unlike tsinfer, which more directly captures the span of edges in the reconstructed ARG based on shared ancestry).

**Figure 3:**
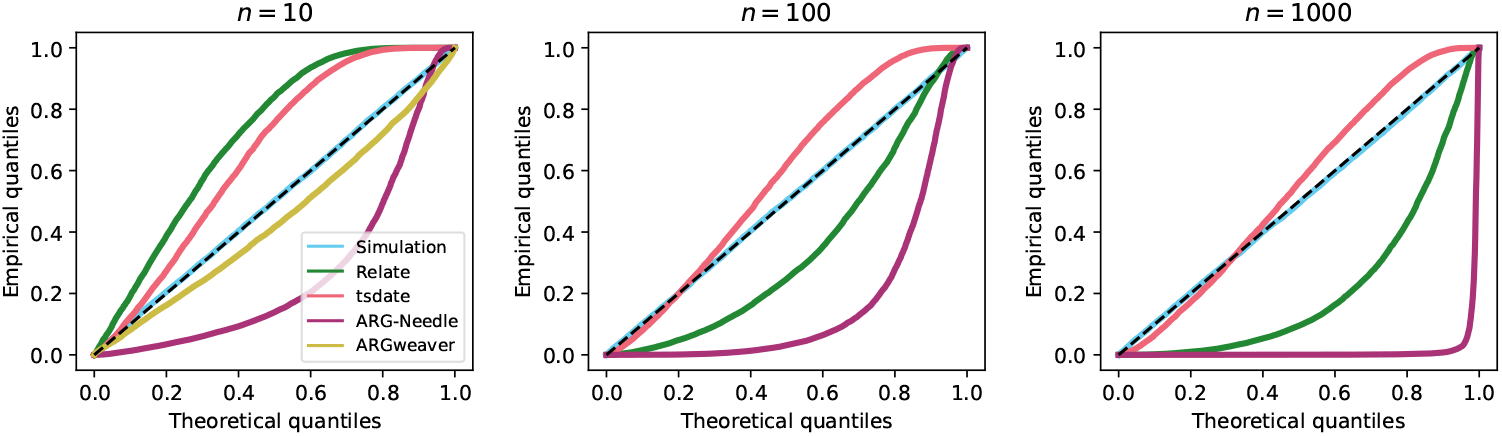
Q-Q plots for edge spans in ARGs simulated with parameters of dataset 1 (dataset 2 for ARGweaver) given in Methods, Section 4.6.1, with *n* = 10 (left panel), *n* = 100 (middle panel), *n* = 1 000 (right panel). Dashed line: diagonal from (0, 0) to (1, 1). ARGweaver results only shown for *n* = 10 due to excessive runtimes for larger sample sizes. Calculated for a random sample of 10 000 edges for each ARG. An overabundance of edges with short (resp. long) spans would drag the points below (resp. above) the diagonal.

ARGweaver accurately captures the edge span distribution, with the small deviation from the diagonal as the simulation used the SMC’ while ARGweaver uses the SMC (SI, Section S1.12), although these results were only calculated for *n* = 10 due to excessive runtimes for larger sample sizes. Note that while it is possible to use the SMC’ model with the latest version of ARGweaver, we found that the resulting ARGs contained cycles, which prevented us from calculating the required probabilities.

ARGs reconstructed by tsinfer/tsdate consistently have an excess of edges with long spans, while for Relate this depends on the sample size; both tools however produce skewed distributions of edge span. This is likely to be, in part, due to the waiting distances between trees being generally skewed, as shown by Deng et al. (2021). Moreover, the ARGs produced by tsinfer contain polytomies (nodes with more than two children), which is also not accounted for under the SMC’ model. This results in the total branch length of a reconstructed local tree to often be greater than it would be if the polytomies were resolved to make the tree binary. In addition, some recombination events that would disrupt the edge under the SMC’ (such as recombination points located on child edges), may not do so when the tree contains polytomies.

Relate first reconstructs the sequence of (correlated) local trees along the entire genome and then calculates edge spans, using an argument based on the similarity of clades subtended by the edge in successive trees. Thus, edge spans are calculated approximately, rather than being explicitly inferred, which we do not account for.

The threading procedure used by ARG-Needle to reconstruct the ARG uses the ASMC model (Palamara et al., 2018) to estimate coalescence times between the added sequence and the closely-related samples already in the ARG, with the lowest possible resulting coalescence time dictating which edge the sequence is threaded to and over what genomic length. This procedure (which relies on a combination of maximum a posteriori and posterior mean estimates for the coalescence times) is optimised for metrics having a direct effect on downstream analyses, and appears to lead to ARG-Needle consistently underestimating edge spans. We note that this is somewhat affected by the choice of time discretisation used within the ASMC (particularly for small sample sizes), but our overall findings do not change significantly when toggling this parameter.

Considering the unconditional distributions of the edge spans in the simulated and reconstructed ARGs (i.e. calculating the observed span of each edge and plotting the overall histogram) results in similar conclusions (SI, Figure S15). Further, histograms of the expected number of mutations per edge (being the product of the observed edge span, observed time-length, and the mutation rate) demonstrate large deviations between the distributions for simulated and reconstructed ARGs (SI, Figure S16).

### 2.3 The distribution of the length of a haplotype block is recovered well by Relate after a suitable correction

A clade *G* of an ARG can be defined through the set of samples it contains, by writing *G* = (*g*_1_, *g*_2_, …, *g*_*n*_), where *g*_*i*_ = 1 if sample *i* is in *G*, and 0 otherwise. The genomic span of *G* is defined as the interval [*a, b*], where *a* is the leftmost position at which the corresponding local tree has a branch subtending exactly the samples in *G*, and *b* is the rightmost such position. For instance, in Figure 1, the clade *G* = (1, 0, 1, 1, 0, 0, 1, 0, 1, 1) (containing samples 0, 2, 3, 6, 8, 9) has a genomic span of 0.55, since *G* first appears at position 0.05 and is broken up by the following recombination event at position 0.60.

Using the SMC’ model, we derive the distribution of the genomic span of a given clade *G*, conditional on the local tree 𝒯 in which it first appears (Methods, Section 4.3). This is approximately exponentially distributed as

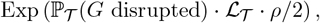

where ℙ_𝒯_ (*G* disrupted) is the probability that *G* either gains or loses at least one sample following the next recombination event along the genome. We also derive an equivalent result when the recombination rate varies along the genome. Since our definition of a clade is equivalent to that of a haplotype block (Shipilina et al., 2023), this distribution is that of the length of a haplotype block. It is important to emphasise that since we condition on *G* and the local tree in which it first appears, this distribution is conditional on the size and age of *G* (as well as the local tree topology and event times, and the recombination map), rather than averaged over all clades.

For a given (simulated or reconstructed) ARG, we can thus calculate a corresponding *p*-value for each observed clade under this model, and again check if these follow the expected uniform distribution using a Q-Q plot. Figure 4 (left panel) shows that the approximation provides an excellent fit for an ARG simulated under the SMC’ (blue points). Clade spans in ARGs reconstructed using tsinfer/tsdate (red points) tend to be over-estimated in general (possibly due to the presence of polytomies), while ARG-Needle (purple points) both under- and over-estimates this quantity. Relate (dark blue points) also tends to over-estimate clade spans; however, through analysing and correcting the causes of this, we propose a correction which effectively removes this bias (green points). Firstly, a clade might only be supported by mutations intermittently along its span, and between these regions Relate does not attempt to keep the clade intact. This causes some clade spans to be too short, and we correct this by extending the calculated span of a clade, if the clade disappears and subsequently reappears within a given distance (which is the cM limit input parameter to DoLoReS, which defaults to 1cM and should be chosen to be relatively large as explained in SI, Section S1.13.1). Secondly, since it is difficult to reconstruct the endpoints of clades exactly, which may results in genomic span being overestimated, we use the leftmost and rightmost mutations that support a given clade to calculate its genomic span. See SI, Section S1.13.1 for full details.

**Figure 4:**
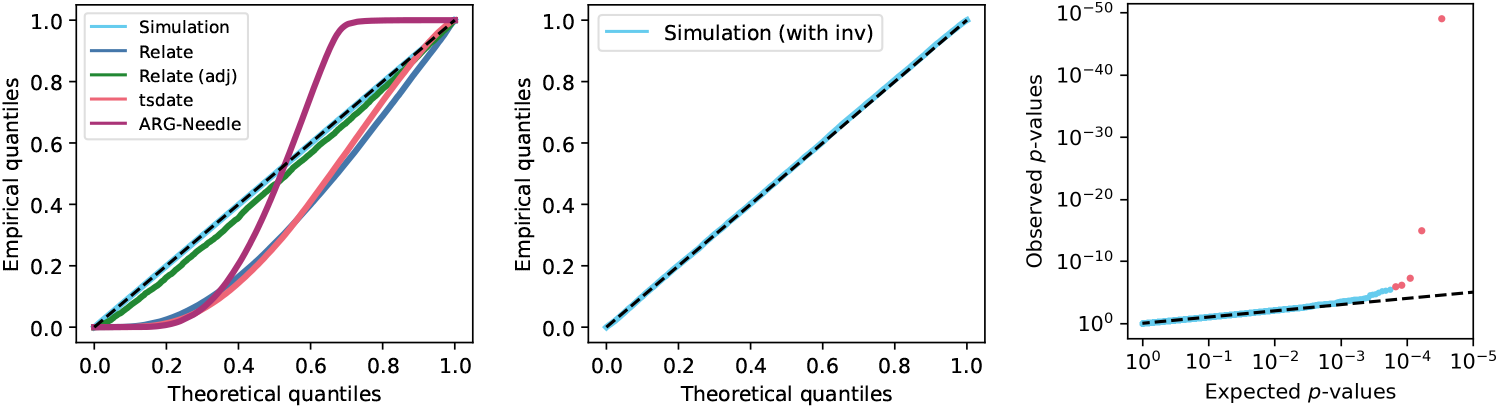
Left: Q-Q plot for clade *p*-values in simulated and reconstructed ARGs with *n* = 100 and parameters as for dataset 1 given in Section 4.6.1. An overabundance of clades with long spans would drag the points below the diagonal. Middle and right: Q-Q plot and *p*-value plot for ARG simulated using SLiM with one inversion under balancing selection (red points correspond to clades with *p*-value below the Bonferroni-corrected significance threshold; blue points correspond to clades with non-significant *p*-values).

### 2.4 Simulated and reconstructed ARGs capture signals of structural variation

Chromosomal inversions are a type of structural variant, whereby as a result of recombination, the genome breaks at two points and the segment between the breakpoints is reinserted in the opposite orientation. While recombination can proceed normally in individuals homozygous for the inversion, in heterozygotes recombination is substantially suppressed in the region containing the inversion, since a crossover recombination within the region is likely to result in the production of unbalanced gametes (Kirkpatrick, 2010). Detecting inversions computationally from sequencing data typically relies on paired-end mapping (detecting reads mapping in the opposite orientation to the reference), split-read methods (detecting reads that map onto the reference with gaps), and *de novo* assembly (directly reconstructing the sequenced genome to look for differences with the reference) (Tattini et al., 2015). For instance, DELLY (Rausch et al., 2012) implements a combination of these approaches and was used to identify hundreds of inversions in 1KGP data (Sudmant et al., 2015). In general, however, such methods suffer from high false positive rates and poor sensitivity, with their performance depending on the size of the inverted region, particularly for short-read sequencing data (Lucas Lledó and Cáceres, 2013); the detection of structural variants from long-read data is challenging due to high sequencing error rates (Sedlazeck et al., 2018). Inversions can also be detected by looking for disrupted patterns of linkage disequilibrium (LD) using population data (Kemppainen et al., 2015; Li and Ralph, 2019; Bansal et al., 2007), however this is sensitive to noise in the LD patterns and cannot be used to reliably detect inversions that are not large and high-frequency.

Suppose that in a given ARG, an inversion happens on an edge *g* which subtends a clade *G*. Suppression of recombination in heterozygotes implies that if a lineage within *G* undergoes a recombination, it will coalesce with lineages in *G* with high probability; likewise, if a recombination event happens on an edge not carrying the inversion, with high probability it will coalesce with edges outside *G* (SI, Figure S3). This implies that inversions can be detected by looking for clades that last for “too long” along the genome due to this local suppression of between-clade recombination. Note that the effect of an inversion differs from simple suppression or general regions of low recombination, since recombination is suppressed in a clade-specific way.

Note that we are imposing the simplifying assumption that recombination in heterozygotes is suppressed completely in the inverted region (so the clade *G* remains completely intact). In reality, recombination can occur in heterozygotes: multiple crossovers occurring in the inverted region would enable this, but such events have relatively low probability unless the inversion region is very large or the recombination rate is very high. Localised reshuffling of clades can also arise through gene conversion, which can indeed be at least as frequent within inversions as outside (Korunes and Noor, 2019; Crown et al., 2018), and for reconstructed ARGs we implement a correction to account for this when calculating the genomic span of a clade (described in SI, Section S1.13.1).

**Test 1:** Phrasing the above as a hypothesis test, for each clade we calculate its genomic span [*a, b*], and calculate a *p*-value as the probability of this clade having a span greater than *b* − *a* (Methods, Section 4.3.1). Simulation studies confirm that for ARGs simulated under the SMC’ model without inversions, these *p*-values are approximately uniformly distributed (Figure 4, left panel, blue points), as expected.

**Test 2:** Alternatively, we can estimate the number of recombination events *R* that occurred within the genomic interval [*a, b*], and calculate a *p*-value as the probability that *G* stays intact after at least *R* recombination events (Methods, Section 4.3.2). This is exactly equivalent to Test 1 when the ground truth ARG and recombination map are known. For reconstructed ARGs, we apply both of these tests since they are susceptible to false positives in different practical scenarios: Test 1 is sensitive to the choice of recombination map and presence of sequencing gaps, while Test 2 can result in false positives if there is a high level of recurrent mutation (which can cause reconstruction errors).

Both of these tests are implemented in DoLoReS, which outputs calculated genomic spans and other characteristics of each clade within an input ARG (in tskit format) and the corresponding *p*-values.

#### 2.4.1 Simulated ARGs

To demonstrate the power of these tests in detecting inversions, we used SLiM (Haller and Messer, 2019; Haller et al., 2019) to simulate an ARG with one 200kb inversion (Methods, Section 4.6.2), under balancing selection, since this is a common mechanism under which polymorphic inversions are maintained in different species (Wellenreuther and Bernatchez, 2018). Figure 4 shows the corresponding Q-Q plot (middle panel) and *p*-values for each clade (right panel) using Test 1. The spans of most clades are unaffected by the inversion, so most of the points on the Q-Q plot adhere tightly to the diagonal. However, there are outliers in the tail (right panel, shown in red) with significant *p*-values (after Bonferroni correction for multiple testing). Figure 5 shows the *p*-values for each clade using Test 1 (above the 0 line) and Test 2 (below the 0 line). The clades with significant *p*-values (using a Bonferroni-corrected significance threshold of 2 *·* 10^−6^) overlap the location of the inverted region, and the clade subtended by the edge carrying the inversion is a significant outlier with the lowest *p*-value. Repeating the simulation using SLiM with no inversion (and otherwise the same parameters), the *p*-values are approximately uniformly distributed and there are no clades with significant *p*-values (SI, Figure S17).

**Figure 5:**
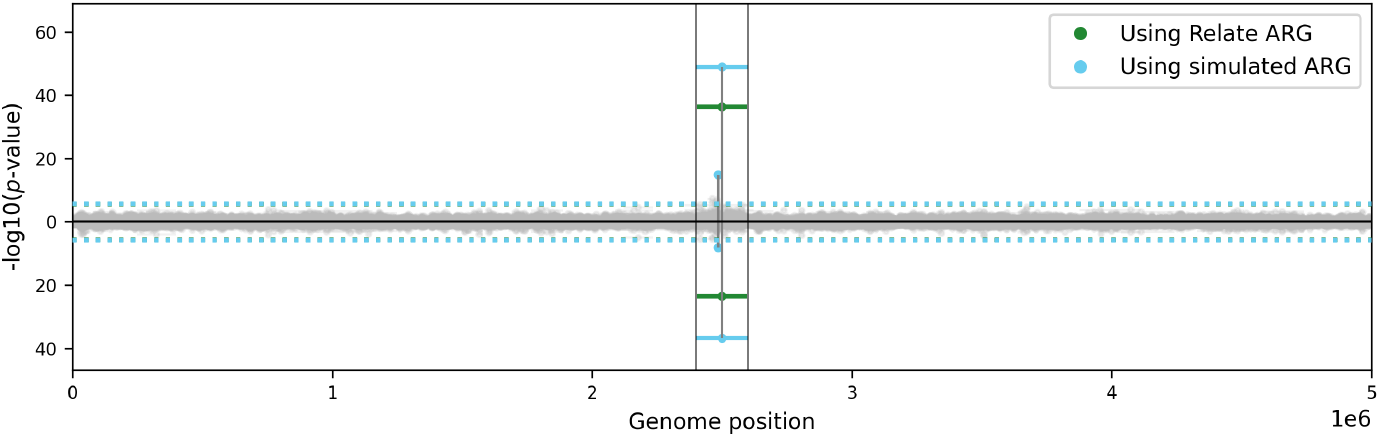
*p*-values for clades in simulated ARG and ARG reconstructed using Relate. Horizontal lines show the genomic span of each clade (with midpoint marked by circle); those with non-significant *p*-values are shown in grey, otherwise in blue and green for the simulated and Relate ARG, respectively. Corresponding *p*-values using Test 1 and Test 2 are shown on the *y*-axis above and below 0, respectively. Vertical black lines delineate the region of the inversion; dotted blue (resp. green) horizontal line shows Bonferroni-corrected significance threshold for calculations using the simulated (resp. Relate) ARG (note these overlap very closely).

#### 2.4.2 Reconstructed ARGs

We next applied DoLoReS to an ARG reconstructed using Relate for the data simulated using SLiM as described above (applying the correction described in Section 2.3). We detect one clade which has significant *p*-values using both tests, shown in green in Figure 5, which is exactly the clade carrying the simulated inversion. The corresponding Q-Q and *p*-value plots for Test 1 are shown in the top row of SI, Figure S18, showing that the Q-Q plot very close to the diagonal. The equivalent plots for a simulation with no inversion are shown in the bottom row, demonstrating that the *p*-values are approximately uniformly distributed and there are no false positives.

We performed further simulation studies to evaluate performance, simulating 100 replicates each for inversions with size varying between 0 and 200kb (and otherwise the same parameters as above), as described in SI, S1.13.2. The resulting ROC curves (SI, Figures S8 and S9) show excellent performance when using simulated ARGs, and that high sensitivity is maintained for ARGs reconstructed using Relate for inversions of 100kb or longer. While our theoretical results hold for clades of any size greater than one, we note that power to detect inversions drops as inversion frequency decreases, since smaller clades have shorter expected spans (so it becomes more difficult to detect outliers).

We also evaluated the performance of our method in predicting inversion genotypes, by considering the samples within the detected significant clades, and compared this against invClust (Cáceres and González, 2015), an inversion detection method based on clustering haplotypes using multidimensional scaling. As described in SI, Section S1.13.3, we found that our method outperforms invClust (SI, Figure S10), while, unlike invClust, not requiring candidate inverted regions as input. We also found that DoLoReS is accurate in predicting the position of the inverted region (SI, Figure S10), with the predicted region overlapping over half of the true region 81% of the time.

### 2.5 Structural variants can be detected using ARGs reconstructed from 1KGP data

We applied DoLoReS to an ARG reconstructed using Relate for the 1KGP (Phase 3) data by Speidel et al. (2019), splitting the ARG into the five super-populations to avoid confounding by population structure, accounting for varying population size through time, and applying several filters to correct for possible sequencing errors and artefacts, which also corrects for phasing errors and the presence of gene conversion (Methods, Section 4.4). The resulting *p*-values are shown in Figure 6 (and SI, Figures S19 and S20). There are a total of 125 significant clades, clustering into 50 localised regions.

**Figure 6:**
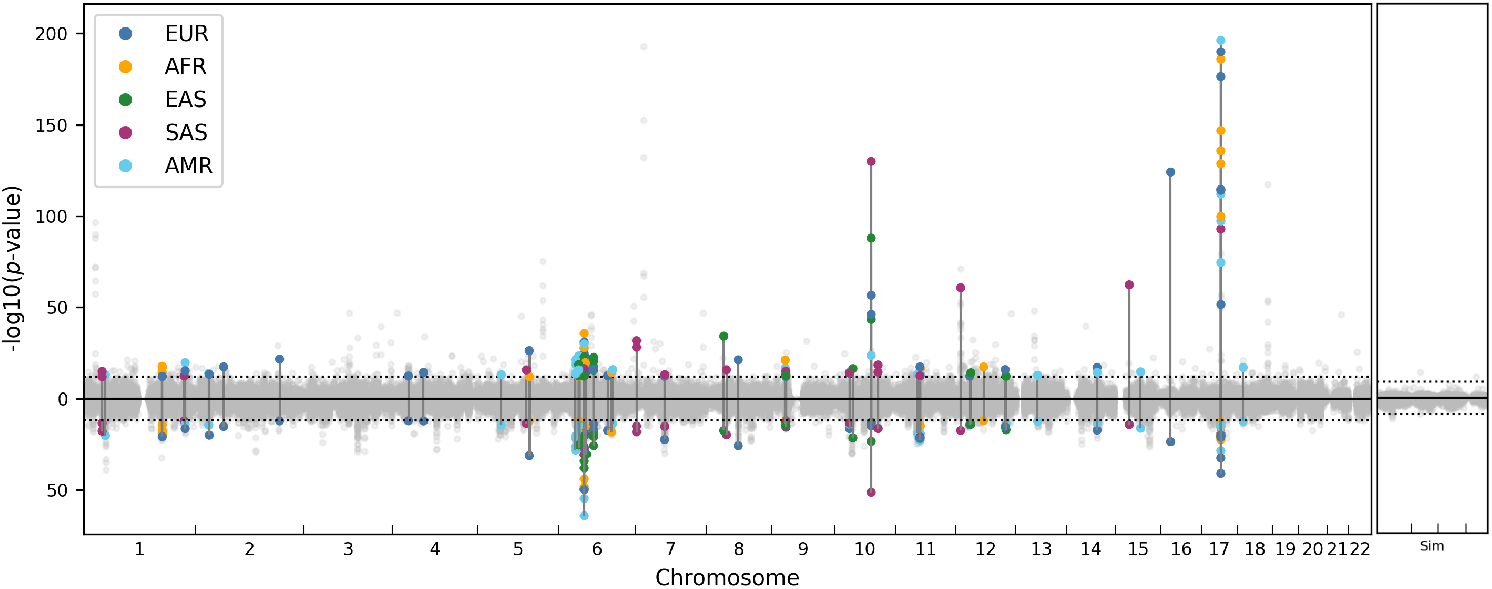
Left panel: *p*-values for clades in 1KGP ARG reconstructed using Relate. Each *p*-value shown as a point, with position on *x*-axis being the midpoint of the corresponding clade span; *p*-values using Test 1 and Test 2 are shown on the *y*-axis above and below 0, respectively. Points for clades with significant *p*-values shown in colour corresponding to the super-population (legend); corresponding *p*-values for Test 1 and Test 2 for each significant clade are connected by solid vertical lines. Dotted black horizontal lines show Bonferroni-corrected significance threshold of 1 *·* 10^−12^. Right panel: results for reconstructed ARG using simulated data, dotted line shows Bonferroni-corrected significance threshold of 1 *·* 10^−9^ (Methods, Section 4.6.3).

#### 2.5.1 17q21.31 inversion

One of the detected regions with the lowest *p*-values, on chromosome 17, corresponds to the known 900kb inversion common in European populations, with two distinct haplotypes H1 and H2, corresponding to inversion non-carriers and carriers, respectively (Stefansson et al., 2005). Since we separately test each clade within each of the population-specific ARGs, multiple clades from the same population can have significant *p*-values (creating the vertical stack of points seen in Figure 6). This is because if the samples are perfectly split into two clades *A* and *B* (of carriers and non-carriers of the inversion, respectively), both will show detectable signal of between-clade recombination suppression. Moreover, sub-clades of *A* or *B* can also have longer genomic spans due to the effects of locally suppressed recombination.

We detect strong signal of this inversion in all populations apart from E. Asian, estimating its average frequency at approximately 24% in European, 15% in American, 6% in S. Asian, and 2% in African populations, which aligns well with prior estimates (Donnelly et al., 2010). The ARG also allows for the estimation of the time of the inversion, through identifying the predicted clade carrying the inversion in each local tree, and extracting the time at the lower and upper end of the branch subtending this clade (Figure 7); this gives an average age of between 8 000 and 123 000 generations. This aligns with the estimates of 3m years of Stefansson et al. (2005) and 2.3m years of Steinberg et al. (2012), but highlights a large amount of uncertainty; the inversion has also been estimated to be much younger at around 100k years by Donnelly et al. (2010). We detect a region where the inversion appears relatively much more recent (highlighted in red on Figure 7), overlapping the 5’UTR region of the *CRHR1* gene (SI, Figure S22). This is the same as the region of very low sequence divergence between the H1 and H2 haplotypes found by Steinberg et al. (2012).

**Figure 7:**
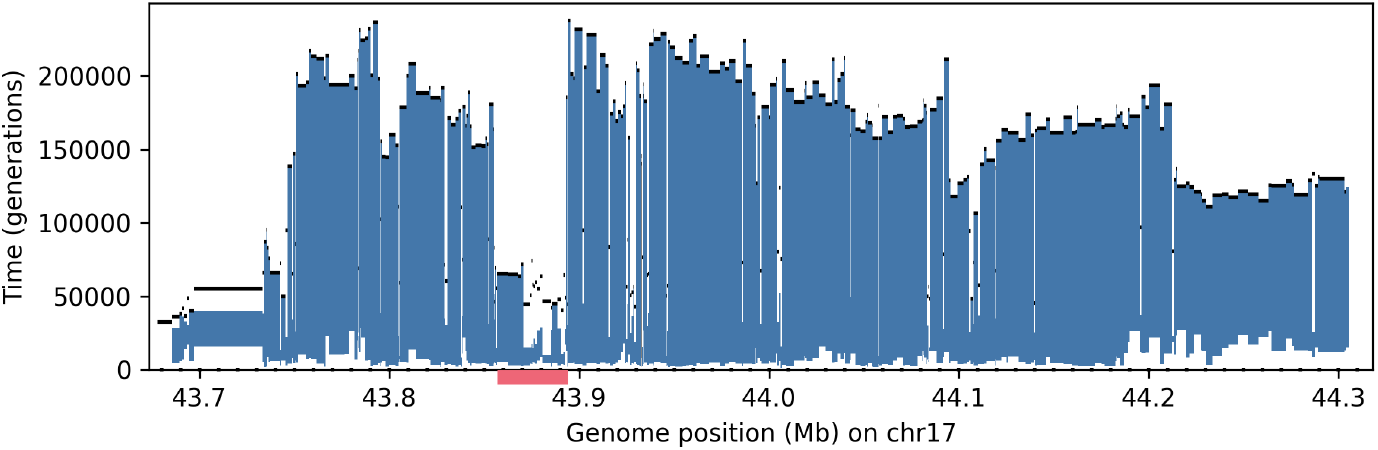
Age estimate for 17q21.31 inversion (the H2 haplotype) using the 1KGP ARG subsetted to European populations. Vertical lines at each genomic position are drawn between the time at the lower and upper end of the branch in the corresponding local tree which subtends the predicted carriers of the inversion. Black lines show MRCA time in full (all-population) ARG. Time is measured in generations, genome positions given in GRCh37 coordinates. Red bar highlights region where inversion appears much more recent.

The topology of the ARG, with H1 and H2 forming two disjoint clades with a very ancient MRCA time (SI, Figure S22K), is consistent with the inversion being very old. From the ARG we infer a change of local tree topology and H1/H2 MRCA time within the highlighted region in Figure 7, which is suggestive of a historic double crossover event between the two haplotypes (with the two breakpoints separated by around 40kb), as posited by Steinberg et al. (2012) (Methods, Section 4.5.3).

We also identify signal of a CNV at around 44.3Mb (SI, Figure S22G), through identifying instances where for a large number of individuals (between 6 and 19 depending on genomic position), their chromosomes perfectly segregate into the same two clades (Methods, Section 4.5.2). The clades we identify correspond to individuals who are homozygous for three copies of a known 25kb CNV at this position (which is in LD with the inversion). This demonstrates that CNVs are also detectable from reconstructed ARGs, through the signal they leave in the data that causes errors in ARG topology reconstruction.

Some significant regions correspond to other known inversions, including on 4q13.2 (Korbel et al., 2007), 11p11.12 (Porubsky et al., 2022), and a possible pericentromeric inversion on chromosome 6 (Martínez-Fundichely et al., 2014).

#### 2.5.2 16p12.2 complex structural polymorphism

The significant clade on chromosome 16 corresponds to a known 1.1Mb structural polymorphism, which was posited to be an inversion in a number of studies (Tuzun et al., 2005; Bansal et al., 2007). This was subsequently shown to be the result of mis-assembly of the reference genome due to complex structural variation in this region, with two common haplotypes that differ by a large (333kb) segmental duplication (Antonacci et al., 2010). Our method thus captures signal of local recombination suppression in individuals heterozygous for this polymorphism.

Other significant hits include known regions of complex structural variation on 11q11 (Korbel et al., 2007) and 15q13.3 (Antonacci et al., 2010).

### 2.5.3 Structural variation on chromosome 6

There is a large number of significant hits on chromosome 6, with a total of 43 significant clades, clustering into 10 distinct regions (SI, Figure S19).

We highlight the top significant hit on 6p11.2, shown in detail in SI, Figure S25. The detected clades span approximately 340kb overall. However, this region contains multiple CNVs (SI, Figure S25G-H), which are not in LD with the detected clades, but clearly cause general distortion of the reconstructed genealogies in this region (as shown by the high proportion of mutations which cannot be uniquely mapped to a branch of the local trees, SI, Figure S25C). As a result of this complex SV landscape, the significant clades are fragmented and absent in many of the local trees within this span (e.g. tree 3 in SI, Figure S25K). Thus, while the signal of recombination suppression is still detected by our tool (since it correctly handles the significant clades disappearing and reappearing within the region), this complexity makes it difficult to confidently assign precise SV boundaries and carrier status.

#### 2.5.4 10q22.3 inversion

We detect a 760kb region of locally suppressed recombination on chromosome 10; details of the genomic spans of the significant clades are shown in Figure 9. This indicates an inversion with an average frequency of approximately 9% (21% in S. Asian, 15% in American, 7% in European and 2% in E. Asian populations), with an age of between 2 625 and 13 392 generations (based on the lower and upper time of the edge subtending the inversion, averaged across its span). Analysis of the genealogies in this region indicates that the non-inverted orientation (with respect to the reference genome) is ancestral.

The presence of segmental duplications (blocks of DNA larger than 1kb and with *>* 90% sequence similarity, occurring multiple times along the genome) can enable non-allelic homologous recombination (NAHR), a potential mechanism through which large structural polymorphisms arise (Lupski and Stankiewicz, 2005). NAHR between two segmental duplications which appear in opposite orientations can, specifically, lead to an inversion of the sequence that they flank. The endpoints of the significant clades (Figure 9A) align exactly with the positions of inverted segmental duplications of length 50kb (Figure 9F), supporting the possibility of an inversion in this region. We confirmed that for predicted carriers of the inversion there is an enrichment of discordantly mapped paired-end reads between these breakpoints.

To further validate this finding, we used complete long-read sequencing data from 47 individuals generated by the HPRC (Liao et al., 2023) and the T2T-CHM13 reference (Nurk et al., 2022). The HPRC sequenced a sample of children in parent-child 1KGP trios, whereas the ARG includes the parents; we thus used tagging SNPs to determine the predicted status of each HPRC sequence. We identify five sequences carrying an inversion within the predicted region, corresponding to one homozygous (HG01258) and three heterozygous (HG01123, HG01978, HG02257) individuals (Figure 8 and SI, Figure S23).

**Figure 8:**
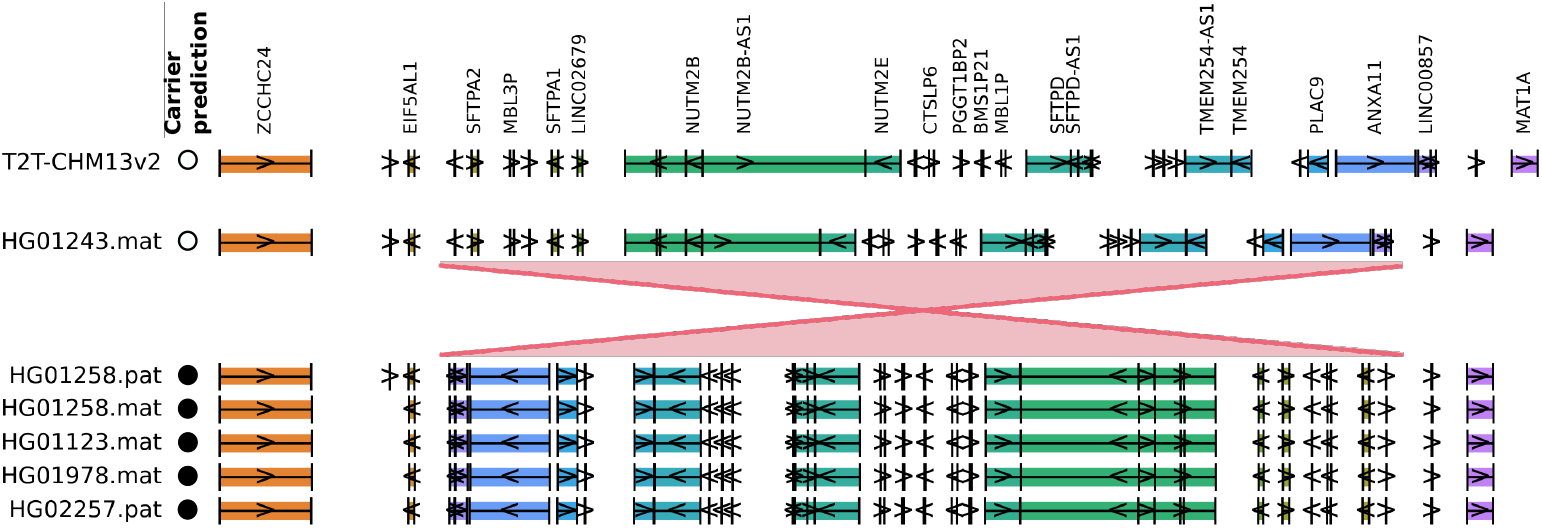
Validation of 10q22.3 inversion using HPRC data for 47 diploid individuals and the T2T-CHM13 reference. Five sequences (out of 95) display an inversion in the predicted region (shown in red), as indicated by a reversal of gene ordering compared to non-carriers (HG01243 chosen as a representative example; see SI, Figure S23 for a comparison against all sequences). Circles show predicted status of carrying the inversion (white = non-carrier, filled = carrier).

**Figure 9:**
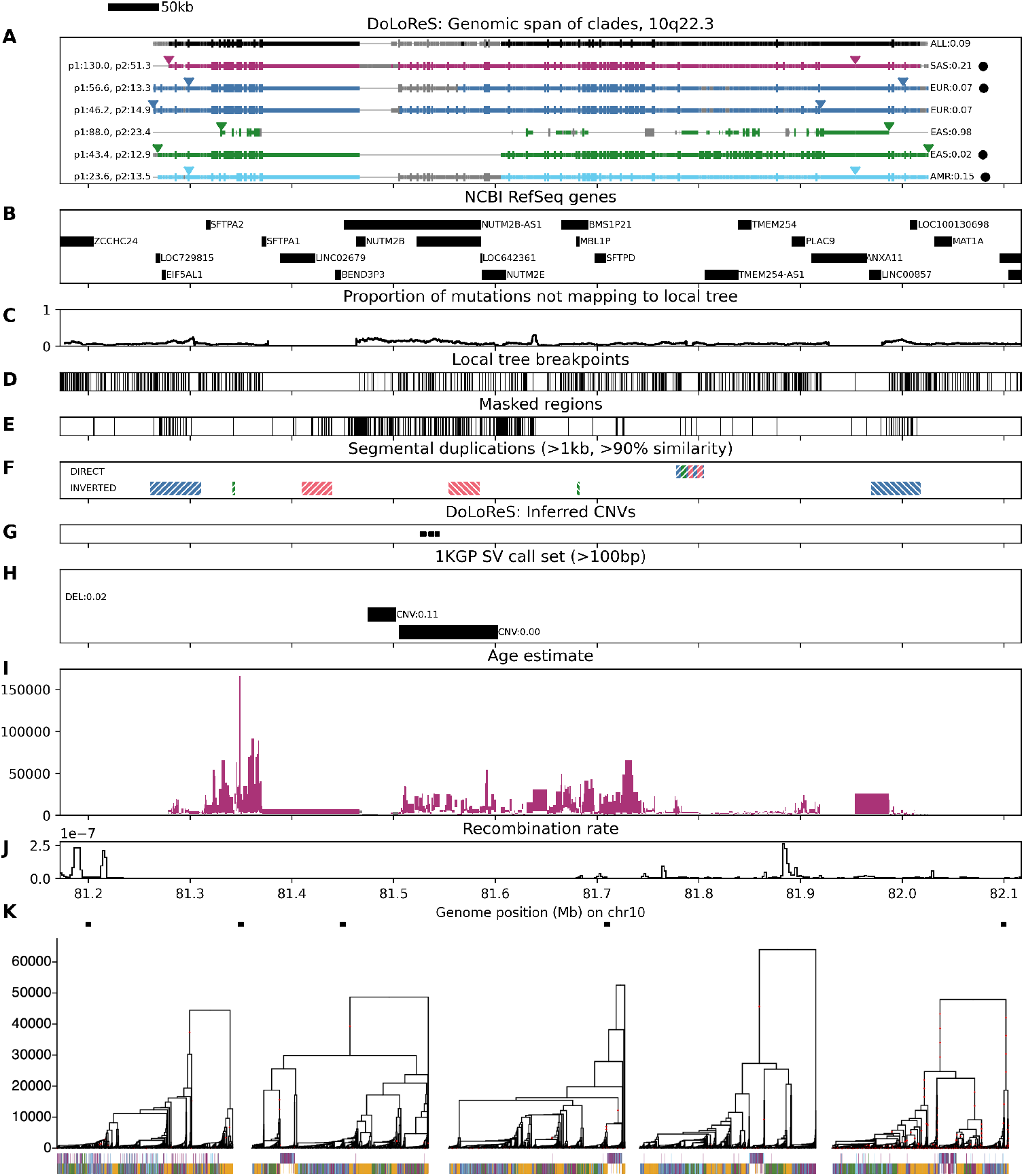
Details of 10q22.3 region. **A**: span of each significant clade shown as horizontal line (grey: clade is not present at that position exactly, but another highly correlated clade is). Vertical lines show positions of SNPs supporting the clade. Left label: *p*-values for Tests 1 and 2. Right label: population frequency. Black circles indicate predicted carriers. See Methods, Section 4.5 for details of this and other panels. **B**: positions of genes (UCSC Genome Browser NCBI RefSeq track). **C**: moving average of the proportion of SNPs not mapping onto the corresponding local tree. **D**: positions of breakpoints between local trees. **E**: regions masked during ARG reconstruction. **F**: segmental duplications. **G**: predicted positions of CNVs based on ARG clade analysis (Methods, Section 4.5.2). **H**: SVs in the 1KGP call set (labels show correlation with predicted inversion carriers). **I**: age of predicted inversion estimated using the ARG subsetted to S. Asian populations (Methods, Section 4.5.3). **J**: recombination rate (using HapMapII recombination map; averaged in bins of 2kbp). **K**: local trees at the positions indicated by squares (chosen at approximately equidistant points across the region, while avoiding regions with known CNVs where ARG reconstruction is unreliable); *y*-axis is time measured in generations; vertical lines drawn below each sample with colour corresponding to population (those belonging to a significant clade drawn in top row).

There are two relatively large CNVs within the span of the inversion: CNV1 at 81 474 561bp (30kb with 0 or 2 copies, overall allele frequency = 0.47) and CNV2 at 81 505 304bp (100kb with 0 or 2 copies, overall allele frequency = 0.02). CNV2 is not in LD with the inversion, but appears to distort the reconstructed genealogy around this region similarly to that on 6p11.2 (resulting in regions where the detected significant clades are broken up temporarily within the ARG, Figure 9A). However, the inverted haplotype has a deletion of CNV1, with all individuals homozygous for the inversion also homozygous for 0 copies of the CNV (using the 1KGP SV call set and analysing read depth as shown in SI, Figure S24; the deletion occurs within the *NUTM2B* gene and is also visible in Figure 8).

The region contains a number of genes associated with lung function (pulmonary-surfactant associated proteins *SFTPA1, SFTPA2, SFTPD*, and *DYDC2*) and immunity (*ANXA11*); SNPs supporting the significant clades in these regions are significantly associated with their expression (Supplementary Table S1). Searching for significant GWAS hits in LD with the predicted inversion carriers (Methods, Section 4.5.4) identified highly correlated variants associated with blood levels of SFTPD (rs2146192: *r*^2^ = 0.67) and Cystatin C (rs55855057: *r*^2^ = 0.64), decreased haemoglobin (rs61859980: *r*^2^ = 0.98, rs61863508: *r*^2^ = 1.0), decreased haematocrit (rs61859980: *r*^2^ = 0.98), and increased levels of blood urea (rs36073865: *r*^2^ = 1.0, rs55838345: *r*^2^ = 0.62, rs17678338: *r*^2^ = 0.6, rs17678338: *r*^2^ = 0.6).

#### 2.5.5 Other SVs

We scanned all of the identified significant regions for those with relatively high frequency, large genomic spans, (direct or inverted) segmental duplications near the identified breakpoints, and evidence from analysis of reads pointing to possible structural variation. This identified a total of 10 inversions (5 novel), 1 known deletion, 1 novel possible CNV or complex rearrangement, 7 complex rearrangements or other variants (3 novel), and 5 regions with strong indications of structural variation but no clear classification (2 novel). For 12 of these 24 variants, we were able to confirm the presence of structural variants in these regions in the HPRC data. The remaining 26 regions show no clear evidence of structural variation. Full details are presented in Supplementary Table S1. The identified variants include:

- a 550kb region on 11q11 (average carrier frequency of 23%), flanked by inverted segmental duplications of length 30kb, and overlapping a CNV from the 1KGP call set which is in LD with the predicted carriers (SI, Figure S26). This corresponds to a known complex rearrangement identified by Korbel et al. (2007); we verified the presence of structural variation in the region in the HPRC sample. The variant overlaps a number of genes and correlates with a large number of GWAS hits for cardiovascular traits.
- a 375kb region on 11q12.1 (200kb away from the region described above), with an average frequency of 4%, flanked by directly oriented segmental duplications (SI, Figure S27). Analysis of reads in this region indicates the presence of a small deletion and inversion within the predicted span correlated with the predicted carriers. This variant is in LD with a variant significantly associated with adolescent idiopathic scoliosis (rs17500359, *r*^2^ = 1.0).
- a 450kb region on 7q11.21, with an average frequency of 13%, flanked by a large number of long inverted segmental duplications (SI, Figure S28) with large number of discordant paired-end reads correlated with predicted carriers. Although we do not find any predicted carriers in the HPRC sample, there is evidence of a 130kb duplication within the region corresponding to a tandem duplication of *RABGEF1* (confirmed as present in 1KGP samples HG01356, HG01351 and HG01140 using analysis of reads), suggesting frequent copy number changes and rearrangements in this region.

We note that while we detect several known regions of structural variation, we do not find significant clades within the span of some other known large inversions, for instance on 8p23.1. We do not identify any clades in the reconstructed ARG that span this 4.5Mb region. We also find that clades that are highly correlated with the predicted carriers of a tag SNP for the inversion (from Wang et al., 2023) only span short regions (at most 15kb) and hence are not significant. Thus, our method fails to detect this inversion since the reconstructed ARG does not capture long-ranging LD within the region. We hypothesise that this might be because the probability of double crossover events within an inversion grows with its size. Our model aims to identify regions and clades with strong recombination suppression, and thus may not identify inversions that are large enough to recombine within their spans in this manner.

## 3 Discussion

We have found that the distribution of edge span is very accurately captured by our approximation based on the SMC’. The differences between the distribution of edge spans in ARGs simulated under the SMC’ model and that in ARGs produced using reconstruction tools are due to both model misspecification and the particularities of each algorithm. Our corrections for Relate result in almost complete recovery of the theoretical distributions. We suggest that the bias seen in ARGs reconstructed using tsinfer stems from the presence of polytomies, which lead to an excess of deep edges with long spans. It is nontrivial to adjust for their presence within our model, and it is not currently possible to break polytomies at random without re-sampling edges in each tree independently (which would prevent a proper calculation of their span). In general, deep edges can arise either through true demographic events or due to inaccuracies in ARG reconstruction; our method can be used to detect deep edges that are likely to be artefactual. We note that apart from ARGweaver, none of the ARG reconstruction methods we consider are explicitly optimised to recover the distribution of edge span, focusing instead on other aspects such as node times, local tree topologies, and patterns of LD (which *are* recovered well in our simulation studies). Our results can potentially be used to improve the estimates of edge span produced by tsinfer and Relate during the topology reconstruction step, and hence also improve the downstream inference of node times. The SMC’-based approximation we construct for the distribution of the length of a haplotype block (the span of a clade of samples) also provides a very close fit, based on simulations. The corresponding tool we develop for detecting regions of locally suppressed recombination has excellent performance on simulated ARGs, as well as (with appropriate adjustments) ARGs reconstructed using Relate. Since the method detects long clade span after adjusting for the age of the clade and the local tree topology, and hence specifically detects localised (between-clade) suppression of recombination, it has the power to discriminate inversions from other genealogy-distorting events, such as point mutations under balancing selection. Our method can be used with arbitrary models of varying population size, and (based on simulation studies) appears to be generally robust to misspecification of demographic history. An inherent limitation of the method is that deep, ancient, population structure can result in a similar signal of localised recombination suppression as SVs, so cannot be easily distinguished. Local adaptation in the face of gene flow at individual loci (“islands of divergence”) can also result in such signals: local adaptation causes stratification at one locus but not another, resulting in clades that span longer-than-expected regions of the genome (though for a clade to be significant would require the presence of linked adaptive loci). The method also cannot identify the specific types of genomic variants that might be causing suppression of recombination in heterozygotes, without utilising other sources of information (such as direct or inverted segmental duplications and other genomic features near the identified regions, or additional analysis of other types of data). However, it provides an alternative line of evidence to methods based on the analysis of paired-end reads, which can miss the presence of complex structural variants or those occurring in regions with poor read mapping.

Applying DoLoReS to the 1KGP ARG reconstructed using Relate identifies a number of regions with both known and novel SVs. Using the ARG allows for the genealogy-based analysis of the age and population frequencies of an SV, and potentially identification of recurrent inversion events. Our tool identifies a large and relatively common inversion on chromosome 10, which has remained unobserved using previous methods due to a lack of clear signal in this region from paired-end read mapping. This demonstrates the power of our method to pick up signals of localised recombination suppression. We limited our detailed examination to regions of size at least 50kb which are well-supported by SNPs, but note that there is a large number of other smaller regions with significant *p*-values (SI, Figure S20), which are more difficult to confidently validate. In general, while our theoretical results hold for clades of any size greater than one, in practice we limited our investigation to large and well-supported clades, since ARG reconstruction is noisy and error-prone, and we sought to focus on the strongest signals to investigate further.

It is difficult to provide guarantees on when our method will achieve a certain false positive rate outside of the scenarios we simulated, since this will depend on the ARG reconstruction method and the properties of the data, so will be application-dependent. We recommend performing simulation studies tailored to the specific species and dataset at hand, to check the performance of this and other ARG-based methods, and calibrate the input parameters. We analysed a large number of relevant metrics and orthogonal evidence to classify the likely reason for recombination suppression within each significant region, including those capturing ARG reconstruction quality (e.g. the proportion of SNPs not uniquely mapping to local trees), whether or not the hits span a centromere, genomic features (e.g. presence of segmental duplications), overlap with genes, an analysis of reads and sequencing depth, analysis of HPRC data using *k*-mer based approaches, and a search of the literature for previous evidence of SVs in these regions. The full details are presented in Supplementary Table S1. As shown in SI, Figure S21, these measures can help to delineate between likely SVs and other sources of suppression.

Roughly half of the detected regions show no clear signals of structural variation based on our analysis. We suggest that other, non-structural reasons may explain allele-specific suppressed recombination in these regions. Previous evidence suggests that recombination crossovers are suppressed within the boundaries of genes expressed in meiosis (McVicker and Green, 2010). This suggests that expression quantitative trait loci (eQTLs) altering meiotic gene expression might impact crossover rates by suppressing recombination crossovers on carriers of one particular allele and in particular in heterozygous individuals, similar to structural drivers. To test whether the identified regions and carriers supported this possibility, we first tested (as detailed in Section 4.5.6) whether our regions are enriched for closely matching gene boundaries. We observe strong enrichment (observed 14 single-gene regions, expected 0.67, OR = 31.2, *p* = 5 *·* 10^−10^), with most of the observed overlaps (10) among those not showing structural evidence. Moreover, the 14 corresponding genes are significantly enriched for being highly expressed during male gametogenesis (9 genes; OR=3.2; *p* = 0.047). Secondly, we tested whether SNPs defining carriers of recombination-suppressed alleles are enriched for known *cis*-eQTLs. Again, we see evidence of enrichment (*p* = 2 *·* 10^−3^), although whether these function in meiotic tisues is unknown. Specific example eQTL regions include *SCMH1* on 1p34.2, *SPATA6* on 1p33, and *ZFAND3* on 6p21.2, all essential for normal spermatogenesis (Takada et al., 2007; Yuan et al., 2015; de Luis et al., 1999). This enrichment of our regions for almost perfect overlap with genes, and eQTLs altering their expression, supports the hypothesis of meiotic allele-dependent suppression of recombination within genes. In contrast, although many (30) of our regions are in LD with GWAS hits, this overlap is not significantly higher than expected by chance (*p* = 0.14). Neither do we observe significant excess overlap with regions under selection identified by Akbari et al. (2024) (*p* = 0.60), although several regions contain one or more significant selection SNPs: 12q24.11 (110.7–111.1Mb), 9p21.1 (31.9–32.1Mb), 2p23.1 (31.8–32.4Mb), 17q21.31 (43.6–44.4Mb), with only the last possessing clear evidence of structural variation. Overall, the data suggest mainly mechanistic drivers of suppressed inter-allelic recombination crossover regions, rather than, for instance, local epistatic selection among trait-influencing variants.

It is clear that even though Relate only uses SNP data and does not explicitly model the presence of SVs, the reconstructed ARG faithfully captures some of these signals. While this allows for their detection, this also means that the genealogies can be distorted by the presence of SVs, most obviously through the phasing errors that they induce in the data. This highlights the value of considering structural variation as an important source of information when developing future ARG reconstruction and analysis methods.

## 4 Materials and methods

### 4.1 Probability that an edge is disrupted by a recombination event

Let 𝒯 be a fixed local tree, and consider a particular edge *b* within this tree. We would like to calculate, under the SMC’ and conditional on 𝒯, the probability ℙ_𝒯_ (*b* disrupted) that the next recombination event arriving along the genome disrupts *b*: that is, it changes the time-length of *b*, or the topology of the tree around *b* (we refer to the latter as *b* being *topologically disrupted*). The possible events that can cause *b* to be disrupted are when (1) the recombination point is on *b* and the coalescence point is not on *b*, or (2) the recombination point is not on *b* but the coalescence point is (SI, Figure S1). The full list of event types that do and do not disrupt *b* are illustrated in SI, Figure S2. By integrating over the possible positions of the recombination point and new coalescence event, we calculate the probability of each illustrated event as detailed in SI, Section S1.3. The resulting expression is in closed form and can be calculated for any given tree and branch.

We derive an equivalent probability ℙ_𝒯_ (*b* topologically disrupted), which only takes into account topology-changing recombination events, as detailed in SI, Section S1.4.

### 4.2 Distribution of edge span

Conditional on the local tree 𝒯, for a particular branch *b*, we are interested in the distribution of the genomic span before *b* is disrupted by a recombination event. We approximate this distribution under the SMC’ by making the assumption that as recombination events arrive along the genome, if they do not disrupt *b* then they also do not otherwise change the rest of the local tree (so 𝒯 stays fixed along the genome). Then since recombination events arrive along the genome as a Poisson process with rate ℒ_*T*_ *· ρ/*2 (where 𝒯_*𝒯*_ is the total branch length of 𝒯), by thinning, recombination events that disrupt *b* arrive at rate

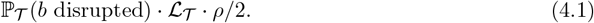

Thus the waiting time until *b* is disrupted by a recombination event is exponential with this rate. We derive an equivalent result when the recombination rate is not constant along the genome (SI, Section S1.8), when considering only topology-changing events (SI, Section S1.8.1), and when we condition on the branch having at least one mutation event (SI, Section S1.8.2).

The assumption that recombination events do not change 𝒯 within the span of *b* is very strong. However, through quantifying the effect of recombination on the height (SI, Section S1.7) and total branch length (SI, Section S1.6) of local trees, we show that the averaged effect of recombination on the rest of the tree does not appear to significantly affect the probability that the given edge is disrupted, making this an excellent approximation (SI, Section S1.11).

### 4.3 Distribution of clade span

For a given local tree 𝒯, we define each clade *G* through the samples it contains as in Section 2.3. We calculate the probability ℙ_*𝒯*_ (*G* disrupted) that, under the SMC’ and conditional on 𝒯, *G* is disrupted by the next recombination event along the genome: that is, that the membership of sample nodes in the clade changes in the next local tree (allowing events that disrupt edges within the clade without changing the group of subtended samples). This can happen when the recombination point is on an edge within the clade and the coalescence point is on an edge outside the clade, or if the recombination point is on an edge outside the clade and the coalescence point is on an edge within the clade, as illustrated in SI, Figure S3. We calculate the probability of these events as detailed in SI, Section S1.5.

The distribution of the genomic span of a clade *G* can then be similarly approximated by making the assumption that recombination events that do not disrupt *G* do not change 𝒯; then the waiting time until *G* is disrupted by a recombination event is exponentially distributed with rate

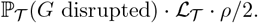

We derive an equivalent result when the recombination rate changes along the genome (SI, Section S1.9).

#### 4.3.1 Test 1

Given an ARG, for each clade *G*^(*i*)^ we calculate its genomic span [*a, b*], and use the approximation (4.1) to compute a corresponding one-sided *p*-value to test whether *G*^(*i*)^ has a significantly longer span than under the null hypothesis of no local (between-clade) recombination suppression:

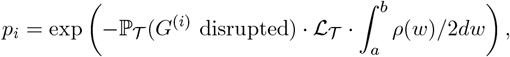

where *ρ*(*w*) is the recombination rate at position *w* (SI, Section S1.9). Simulation studies show that the test has excellent performance for simulated ARGs even for small inversions, and maintains good sensitivity for detecting inversions of over 50kb for Relate ARGs, while accurately pinpointing their position (SI, Section S1.13.2).

#### 4.3.2 Test 2

Under the model described above, an equivalent test can be constructed using the number of recombination events *R* occurring within the genomic span of *G*, which has a geometric distribution with rate ℙ_*𝒯*_ (*G* disrupted) (SI, Section S1.14). For each clade *G*^(*i*)^ within a given ARG, we thus calculate a corresponding (one-sided) *p*-value

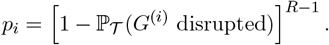

In practice, *R* is unknown, so we instead calculate the number of breakpoints between local trees within [*a, b*] (this is conservative as it strongly underestimates the number of recombination events in this interval).

### 4.4 1KGP ARG

We use tskit (Kelleher et al., 2016) to split the ARGs into the five super-populations (EUR: European, AFR: African, SAS: S. Asian, EAS: E. Asian and AMR: American), and analyse them separately. We adjusted for varying population size (as estimated by Relate) as detailed in SI, Section S1.9.1. We also applied the corrections detailed in SI, Section S1.13.1 (setting *L* = 1cM), which correct for reconstruction error and also the presence of gene conversion within inversions (by extending the calculated span of a clade if it disappears and then reappears within a short genomic span). We used the 1KGP genomic mask (which marks whether each nucleotide passes a set of quality filters, based on depth of coverage and reads mapping) to set the recombination rate in regions marked as ‘not passing’ to 0 (to avoid false positives for Test 1). Additionally, to be robust to the presence of phasing switch errors in the data, instead of defining a clade by the samples it contains as described in Section 2.3, we instead count how many samples from each individual it contains. That is, we assign each clade *G* a “genotype” ID (*G*_1_, …, *G*_*N*_), where *G*_*i*_ is the number of sequences (0, 1, or 2) within *G* from individual *i*, and consider the clade present in a given local tree if there is a clade with the same genotype ID. We filter out clades supported by fewer than 10 mutations, spanning less than 50kb or fewer than 10 local trees, and those having fewer than 10 or more than *N* − 10 samples (where *N* is the total number of samples in the ARG); this leaves 107k clades. Tests 1 and 2 are then applied for each remaining clade independently (using the HapMapII GRCh37 recombination map for Test 1, and counting the number of local tree breakpoints to estimate the number of recombination events *R* for Test 2). We require both *p*-values to be below the Bonferroni-corrected threshold of 1 *·* 10^−12^ (being 0.05 divided by the total number of clades in the ARGs).

### 4.5 Analysis of results

For each significant clade *G*, in each local tree within its genomic span, we identify the clade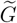with which it is most highly correlated (as measured by the correlation coefficient between their genotype IDs). In Figure 9A, the span of each significant clade *G* is plotted as a solid horizontal line (in colour corresponding to the super-population), if within the local tree at the given genomic position, there is a clade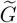 with a correlation coefficient of at least 0.95 (and in grey if the correlation coefficient is between 0.9 and 0.95). The leftmost and rightmost positions at which the clade appears in the ARG exactly are indicated by triangles.

We also compute the combined genotype ID for the corresponding predicted clade of samples in the full (all-population) ARG, and calculate its genomic span (shown in black in Figure 9A). Existence of this “superclade” provides additional evidence that the clades identified independently in each population ARG are not false positives.

#### 4.5.1 Genomic features and measures of ARG reconstruction quality

For each identified region, we extract the positions of nearby genes, using the UCSC Genome Browser NCBI RefSeq track (Pruitt et al., 2005). To check for known genomic features that tend to co-occur with SVs, we also extract the positions of segmental duplications of length greater than 1kb and *>* 90% sequence similarity falling within this region (Bailey et al., 2001, 2002), and the positions of SVs in the 1KGP call set (Sudmant et al., 2015). To detect issues caused by ARG reconstruction artefacts, we extract the positions of breakpoints between adjacent local trees, and the regions masked during ARG reconstruction (labelled as “not passing” in the 1KGP pilot mask); we also calculate a moving average (in 10kb windows) of the proportion of SNPs in the 1KGP data that cannot be uniquely mapped onto a branch in the local tree at the corresponding position (as a measure of ARG reconstruction error). To detect instances where poor estimation of event times may inflate the probability that the clade is disrupted by recombination, we also checked (for each significant clade) the proportion of mutations that fall within vs. outside the clade, and compared this to the average proportion of tree total branch length within vs. outside the clade (expecting these quantities to be similar if times are well estimated).

#### 4.5.2 Phasing errors

Using the individual-based definition of a clade means that it is possible for two clades in the same local tree to have identical IDs, if one clade contains one chromosome from each of *k* individuals, and the other clade contains the other chromosome for each of these *k* individuals. This has negligible probability to arise by random chance unless *k* is very small (4 or less, based on simulations with 1KGP-like parameters). This can, however, arise for larger *k* as the result of phasing errors due to structural variation, in particular due to mis-alignment of CNVs which results in ARG reconstructed artefacts. We thus record all instances where, in a local tree, two clades of size at least 6 have identical IDs, to look for this signal.

#### 4.5.3 Inversion age

A lower and upper bound on the age of an inversion can be obtained by identifying the most highly correlated clade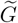 in each local tree within the genomic span of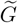, and (if the correlation coefficient is at least 0.9) obtaining the times at the bottom and top end of the branch that subtends *G* (call these times *s* and *t*, respectively).

While a neutral mutation can arise at any time uniformly distributed along this branch, an inversion prevents certain types of recombination events, so the changes in ARG topology and in *s* and *t* along the genome are informative of its age. Suppose that the inversion is old. Then Figures 10A-B imply *s* should be relatively recent (being approximately distributed as the coalescence time of *k* samples), while Figures 10C,D,H imply that both *t* and the MRCA time will be large, since they are constrained to be larger than the age of the inversion. Moreover, if the carriers and non-carriers form two (disjoint) clades in the ARG, the only type of event that can change this topology is that shown in Figure 10H, which has a very low probability under the SMC’ assumptions. However, a double crossover within the inverted region can allow any of the events shown in red, locally changing *s, t*, the MRCA time and/or the topology, in the region between the two recombination breakpoints.

**Figure 10:**
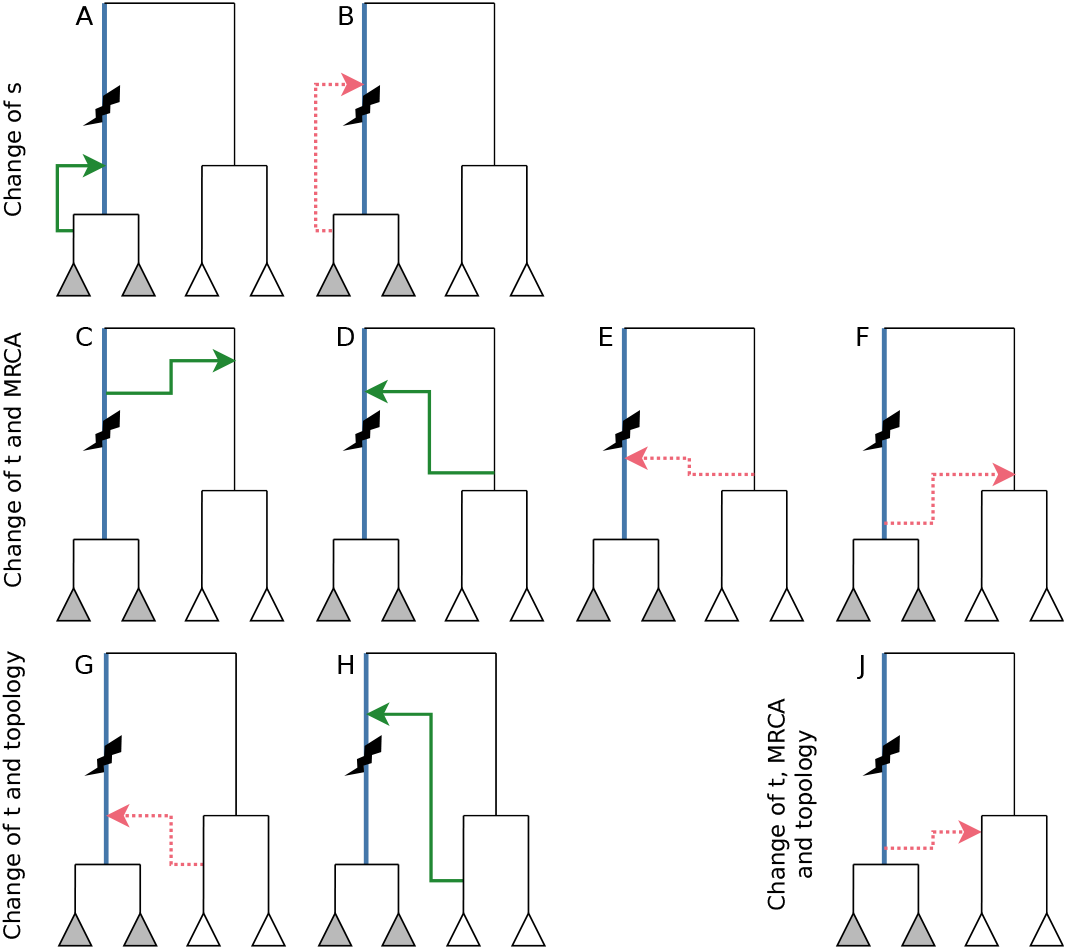
Possible recombination events that change *s* (time of the bottom end of the branch subtending the samples carrying the inversion, shown in blue), *t* (the of the top end of the branch), the MRCA of the carriers and non-carriers, and/or the order of coalescence of the clades. Inversion shown as lightning bolt. Recombination events shown as arrows, where the start of the arrow shows the time and location of the recombination event and the arrowhead shows that of the subsequent coalescence; green solid (resp. red dotted) arrows show feasible (resp. infeasible) events.

This description aligns with the observed ARG topology and branch time estimates for the 17q21.31 inversion (SI, Figure S22): the carriers and non-carriers form two disjoint clades with a very large MRCA time for most of the inverted region, apart from the region highlighted in red in Figure 7. In this region the corresponding local tree topologies change in a way consistent with an event of type F around position 43.86Mb (which changes *t* and the MRCA time), followed by a number of recombination events of type J between 43.87-43.90Mb (which change *t*, the MRCA time, and the tree topology as shown in SI, Figure S22K).

#### 4.5.4 Significant eQTLs and GWAS hits in LD with identified variants

For each identified significant clade, we searched the Open Targets Genetics catalog (Ghoussaini et al., 2021; Mountjoy et al., 2021) for genome-wide significant GWAS hits (*p <* 5*·* 10^−8^) in LD (*r*^2^ *>* 0.6) with SNPs supporting the clade. The full details of the identified SNPs are presented in Supplementary Table S2. We also checked SNPs supporting the clade for significant associations with gene expression using the QTL catalog.

#### 4.5.5 Analysis of HPRC data

For each identified significant region, we predicted the carrier status of each sequence in the HPRC dataset by checking whether it carries SNPs that occur on the branches that subtend the significant clades. For each sequence, we then calculated *k*-mer counts (setting *k* = 20) in (and around) the region, and counted the number of *k*-mers deleted, duplicated, or inverted when compared to the T2T-CHM13v2 reference. We then calculated the correlation between these *k*-mer counts and predicted carrier status. To look for changes in ordering indicating rearrangements or inversions, we also (1) plotted the ordering of genes in the region as annotated for each sequence, and (2) selected a subsample of equally spaced *k*-mers on the T2T-CHM13v2 reference, and plotted the relative positions of these *k*-mers on each sequence.

#### 4.5.6 Testing for enrichment

We tested for enrichment of our identified regions for overlapping with genes, genes involved in meiosis, SNPs under selection, GWAS hits and eQTLs. For each significant region we selected a tagging SNP in the middle of the region, constructed a list of 10 best-matched SNPs on the same chromosome (matching on the overall frequency, frequency in Europeans, and average recombination rate within the surrounding 1Mb region), and defined a region around the matched SNPs of the same physical (and approximately the same genetic) size. We then checked each matched region for overlapping SNPs under selection from Akbari et al. (2024), genes (using the Genome Browser NCBI RefSeq track), genes involved in meiosis (taking clusters 2-4 from Xia et al. (2020, Table S3) and filtering for genes within the top 25% by expression level in testis bulk sequencing data (obtained from the GTEx portal on 19/01/2025, file gene reads v10 testis.gct.gz), and GWAS hits and eQTLs as described in Section 4.5.4 (filtering the Open Targets eQTLs for those with an association score of at least 0.6). We then calculated bootstrapped *p*-values by resampling using these matched regions.

### 4.6 Simulation parameters

#### 4.6.1 Neutral simulations

Using stdpopsim (Adrion et al., 2020), a library of standardised population genetic simulation models integrated with msprime, we simulated two ARGs under the SMC’ with *n* = 100 (haploid samples) and the following two sets of parameters:

- dataset 1: Chr21 with HapMapII GRCh38 recombination map, mutation rate 1.29 *·* 10^−8^ per site per generation, *N*_*e*_ = 10 000 diploids, constant population size model;
- dataset 2: 5Mb of Chr21 with flat recombination map, recombination rate 1.2 *·* 10^−8^ per site per generation, mutation rate 1.29*·*10^−8^ per site per generation, *N*_*e*_ = 10 000 diploids, constant population size model.

The mutation rate and *N*_*e*_ estimates are the defaults in stdpopsim and in line with other commonly-used estimates for human data (Scally and Durbin, 2012; Takahata, 1993).

For each simulated dataset, we used ARGweaver, Relate (v1.1.9), tsinfer/tsdate (v0.3.0 and v0.1.5), and ARG-Needle to reconstruct an ARG, using the true simulation parameters as inputs for each tool (and for ARGweaver, a time discretisation grid with 100 points and selecting the MAP ARG from 1000 posterior samples, for ARG-Needle using 50 time discretisation points). We sense-checked the output of the ARG reconstruction methods using a number of metrics (comparing the MRCA times, local tree topologies, and LD decay, against the simulated ARGs).

#### 4.6.2 Simulations with inversion

We used SLiM (Haller and Messer, 2019; Haller et al., 2019) to simulate an ARG with one inversion (*n* = 100, 5Mb with recombination rate 1*·*10^−8^ per bp per generation, constant population size 10 000, inverted segment length 200kb, neutral mutations at rate 1 *·* 10^−8^ per bp per generation added using msprime). We used the recipe in Section 14.4 of the SLiM manual (version of 31 August 2024), which simulates balancing selection through a frequency-dependent fitness effect 1 − (*f* − 0.5) *·* 0.2 (where *f* is the current frequency of the inversion), to maintain the inversion at near intermediate frequency.

#### 4.6.3 1KGP simulation

To simulate data using parameters similar to the 1KGP data, we used stdpopsim with the AmericanAdmixture 4B11 demographic model, simulating the same number of African, European, Asian and Admixed samples as in the 1KGP, for chromosomes 18-22 (using the HapMapII GRCh37 recombination map). We then applied the 1KGP genomic mask and reconstructed an ARG using Relate, and applied the methods described in Section 4.4 to calculate *p*-values, using a Bonferroni-corrected significance threshold of 1 *·* 10^−9^ (using the individual-based definition of a clade). The results are shown in Figure 6 (right panel).

## 5 Data availability statement

Code implementing DoLoReS is publicly available at github.com/a-ignatieva/dolores. Scripts used to produce and analyse the simulated and 1KGP data are publicly available at github. com/a-ignatieva/dolores-paper. Simulated data and 1KGP results are publicly available at doi.org/10.6084/m9.figshare.29256770.v1.

## 6 Acknowledgements

This work was initiated at the *Stochastic modelling in the life sciences* Junior Trimester Programme held at the Hausdorff Research Institute for Mathematics, University of Bonn (funded by the Deutsche Forschungsgemeinschaft under Germany’s Excellence Strategy EXC-2047/1-390685813). SRM and AI are supported by the Wellcome Trust (Investigator Award 212284/Z/18/Z), MF by the Knut and Alice Wallenberg Foundation (Program for Mathematics, grant 2020.072, hosted by the Department of Statistics, University of Warwick), JK by the EPSRC (research grant EP/V049208/1), and JS by the ERC (Starting Grant ARGPHENO 850869). We thank Yan Wong and Pier Palamara for helpful discussions, and Sebastian Quintanilla Terminel and Kari Heine for useful comments.

## Supplementary Information

**S1 Supplementary Methods 1**

S1.1 The SMC’ model and ARG reconstruction 1

S1.2 Notation and background 2

S1.3 Probability that an edge is disrupted by a recombination event 3

S1.4 Probability that an edge is topologically disrupted by a recombination event 6

S1.5 Probability that a clade is disrupted by a recombination event 7

S1.6 Change in total branch length of tree following a recombination event 8

S1.7 Change in tree height following a recombination event 9

S1.8 Distribution of edge span 9

S1.9 Distribution of clade span 12

S1.10 Quality of approximation to the distribution of edge span 13

S1.11 Effects of recombination on a local tree 13

S1.12 Comparison of simulation models 15

S1.13 Detection of local recombination suppression: Test 1 16

S1.14 Detection of local recombination suppression: Test 2 19

**S2 Proofs 22**

**S3 Supplementary Figures 34**

## S1 Supplementary Methods

### S1.1 The SMC’ model and ARG reconstruction

ARGs were first described by Griffiths and Marjoram (1997) as realisations of the coalescent with recombination (CwR), a stochastic process operating backwards in time, generating a genealogy through a sequence of coalescence and recombination events (Hudson, 1983). Wiuf and Hein (1999) reframed the CwR as a stochastic process operating spatially along the genome: starting with the leftmost endpoint, local trees are generated sequentially moving to the right, reshaped by recombination events. While calculating the properties of the ARG under both frameworks is generally intractable, this seminal work spurred on a suite of simplifying approximations, enabling applications to large-scale genomic data.

The sequentially Markovian coalescent (SMC) model, proposed by McVean and Cardin (2005), imposed the assumption that the process along the genome is Markovian, which, in essence, prohibits recombination events in genetic material not ancestral to the sample. This was followed by the SMC’ extension (Marjoram and Wall, 2006), which was shown to be an excellent approximation to the CwR, based on the joint distribution of pairwise coalescent times and a quantification of bias in population size estimates (Wilton et al., 2015), and the distribution of the next local tree conditional on the current one in a two-locus model (Hobolth and Jensen, 2014). Thus, for a small trade-off in accuracy, the SMC’ model offered a substantially more tractable way of calculating analytic approximations to various quantities of interest, such as the correlation between coalescence time and linkage probability for a randomly sampled pair of sequences (Eriksson et al., 2009), and identity-by-descent tract length distributions and related quantities (Harris and Nielsen, 2013; Carmi et al., 2014). It also enabled the development of powerful new inference methods: for instance, by considering the genealogy of a single pair of sequences, Li and Durbin (2011) developed a HMM-based approach (the pairwise SMC, or PSMC) for inferring the history of human population sizes, which was subsequently extended by Schiffels and Durbin (2014) to multiple samples.

**Figure S1:**
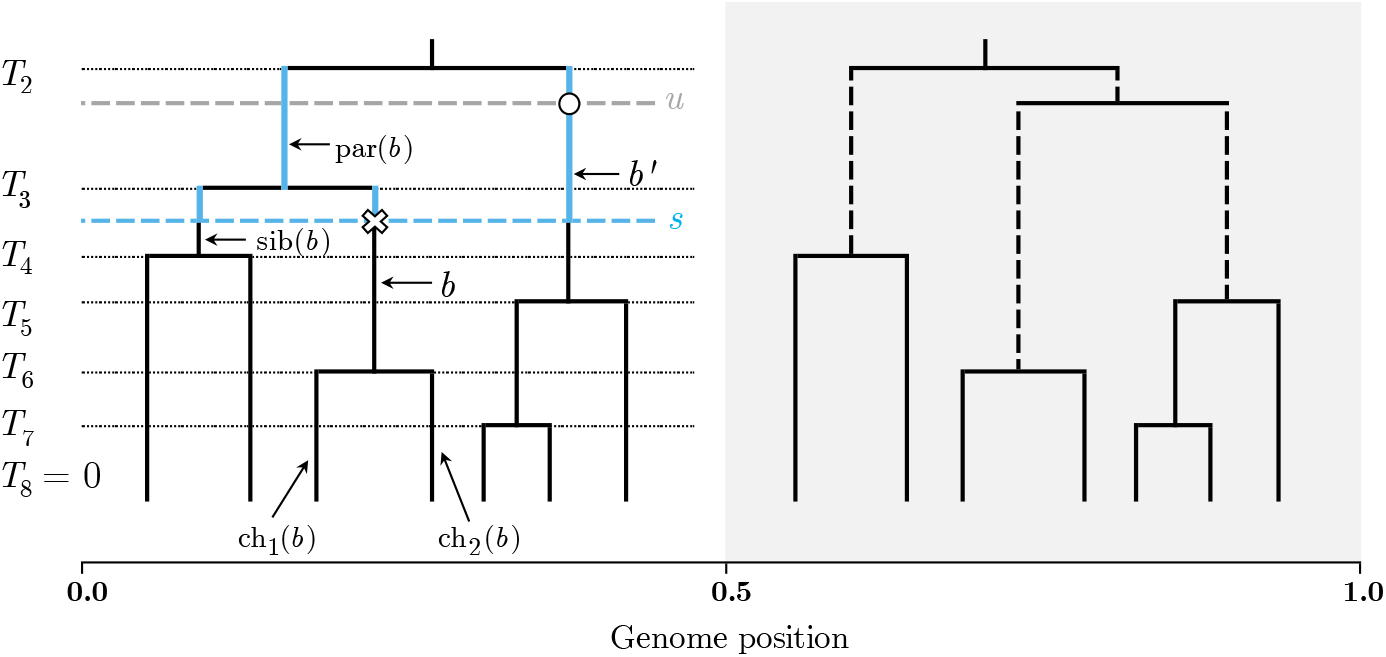
Illustration of the notation used throughout. The ARG has two marginal trees, where the tree on the left is𝒯, with *n* = 7. Coalescent event times are shown as black dotted lines. For the edge labelled *b, t*^↑^(*b*) = *T*_3_, *t*^↓^(*b*) = *T*_6_, *d*^←^(*b*) = 0, *d*^→^(*b*) = 0.5 (so the span of the edge is 0.5); *A*(*b*) = *{b*, sib(*b*), par(*b*)*}* and ℬ(*b*) = *{b*, sib(*b*), ch_1_(*b*), ch_2_(*b*)*}*. The recombination event occurs at genomic position 0.5; the recombination point ℛ = (*b, s*) is shown as a cross; the coalescence point 𝒞= (*b*′, *u*) is shown as a circle; *n*(*s*) = 3 and *L*_*𝒯*_ (*s*) gives the total length of the edges shown in blue. The tree on the right 𝒯′ is obtained by pruning the subtree below ℛ and reattaching at 𝒞; solid vertical lines show edges that have not been affected by the recombination event.

Meanwhile, the definition of the ARG has become decoupled from its initial description as the realisation of a stochastic process, to more broadly denote a genealogical network that captures genetic inheritance. Using this looser definition, the problem of explicitly reconstructing plausible ARGs from sequencing data has seen significant recent progress driven by the use of heuristic methods and principled approximations to the CwR. ARGweaver (Rasmussen et al., 2014) implements an MCMC scheme based on a time-discretised version of the SMC (or SMC’) to obtain posterior samples of ARGs compatible with a given dataset. Relate (Speidel et al., 2019) and tsinfer/tsdate (Kelleher et al., 2019; Wohns et al., 2022) reconstruct a single ARG from data, by using methods based on the Li and Stephens (2003) framework to first reconstruct the topologies and then estimating the edge lengths using Bayesian approaches with coalescent-based priors. ARG-Needle (Zhang et al., 2023) reconstructs a single ARG by sequentially threading in each sample, by first identifying the most closely related samples already in the ARG via genotype hashing, and subsequently estimating coalescence times under the Ascertained Sequentially Markovian Coalescent (ASMC) model (a coalescent-based HMM). These methods scale to thousands of human genome-length samples and have already been applied to many large-scale datasets, resulting in powerful inference of evolutionary events and parameters, such as the history of human demography (Wohns et al., 2022), past population sizes (Speidel et al., 2019), signals of selection (Hejase et al., 2022), and genetic associations for complex traits (Zhang et al., 2023).

### S1.2 Notation and background

The notation is illustrated in Figure S1. Let 𝒯 be a fixed local tree with *n* leaves. Denote by *T*_*i*_ (for *i* ∈ *{*2, *· · ·, n}*) the population-scaled time at which the number of lineages in 𝒯 jumps from *i* to *i* − 1, with *T*_2_ being the time of MRCA and setting *T*_*n*+1_ := 0. Let *n*(*t*) be the number of lineages at time *t*, so *n*(0) = *n*, with *n*(*T*_*j*_) = *j* − 1 and *n*(*t*) = 1 for *t* ≥ *T*_2_.

For an edge *b* ∈ 𝒯, denote the lower end time by *t*^↓^(*b*) ≥ 0 and the upper end time by *t*^↑^(*b*) ≤ *T*_2_, with the *time-length* of the edge given by *t*(*b*) = *t*^↑^(*b*) − *t*^↓^(*b*). Let *d*^←^(*b*) and *d*^→^(*b*) be the leftmost and rightmost endpoints of the genomic span of edge *b*, respectively, with its *span* given by *d*^→^(*b*) − *d*^←^(*b*).

Let par(*b*), sib(*b*), ch_1_(*b*) and ch_2_(*b*) denote the parent, sibling, left child and right child edge of *b* respectively, such that we have the following relations:

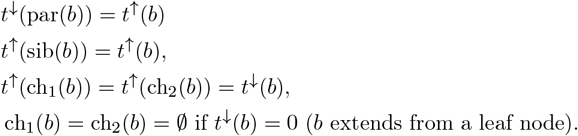

Define the sets of edges 𝒜(*b*) := *{b*, sib(*b*), par(*b*)*}* and ℬ(*b*) := *{b*, sib(*b*), ch_1_(*b*), ch_2_(*b*)*}*, and denote by *b*_*r*_ the root lineage extending past the MRCA node.

Let *L*_𝒯_ (*t*) be the sum of edge lengths in 𝒯 above time *t* and up to *T*_2_:

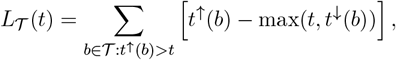

so that *L*_*T*_ (0) is the total branch length of 𝒯 (condensed asℒ_*T*_ := *L*_*T*_ (0) in the main text). Denote by 𝒯_*x*_ the local tree at position *x* along the genome.

Under the SMC’, moving along the genome, a recombination event happens after an exponentially distributed waiting time with rateℒ_*𝒯*_ (0) *· ρ/*2; when this event happens, a locationℛ is selected uniformly at random along the edges of 𝒯, say on edge *b* at time *s*, which we denote as

ℛ∈ *b* orℛ = (*b, s*). A new coalescence point𝒞 is selected by allowing the recombining lineage to coalesce at rate 1 with all the lineages present above time *s* (including *b*_*r*_). We denote a coalescence point on edge *b*′ at time *u* as 𝒞 ∈ *b*′ or𝒞 = (*b*′, *u*). The next tree along the genome is then formed by pruning the subtree below the recombination pointℛand reattaching it at the chosen coalescence point 𝒞. We writeℛ ∈ ℬ(*b*) to mean that the recombination point is on one of the edges inℬ(*b*).

The difference with the spatial formulation of the CwR is that the coalescence point is restricted to be on the local tree 𝒯, whereas under the CwR it could be placed on any edges of the ARG corresponding to the full sequence of trees to the left of the recombination position. In essence, the SMC’ approximation disallows any recombination events that occur in non-ancestral material, making the process Markovian along the genome.

The difference between the SMC and SMC’ models is that under the SMC’, the coalescence point can be chosen on the same edge *b* containing the recombination point, above time *s* (so that recombinations can occur that do not change the tree topology or edge lengths), whereas events of this type are disallowed under the SMC.

### S1.3 Probability that an edge is disrupted by a recombination event

Considering a fixed edge *b* ∈ 𝒯, when a recombination event occurs, we would like to know the probability that *b* is affected by this recombination event. This includes both changes in the time-length of *b* and events that change the topology of the clade around *b* (we will say in these cases that *b* is *topologically* disrupted by the recombination). This can happen because either (1) the recombination point is on *b*′ ∈ ℬ(*b*) and the coalescence point is not on *b*′, or (2) the recombination point is not on *B*(*b*) and the coalescence point is on *b*. The possible scenarios that do and do not disrupt *b* are illustrated in Figure S2.

We have, for a given tree𝒯,

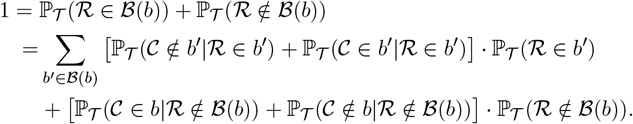

The probability that *b* is disrupted by the recombination event is thus

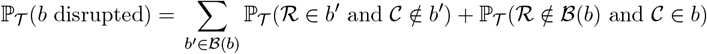

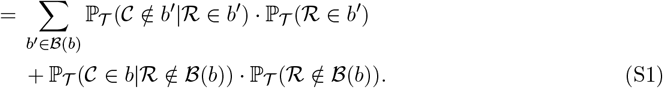

**Figure S2:**
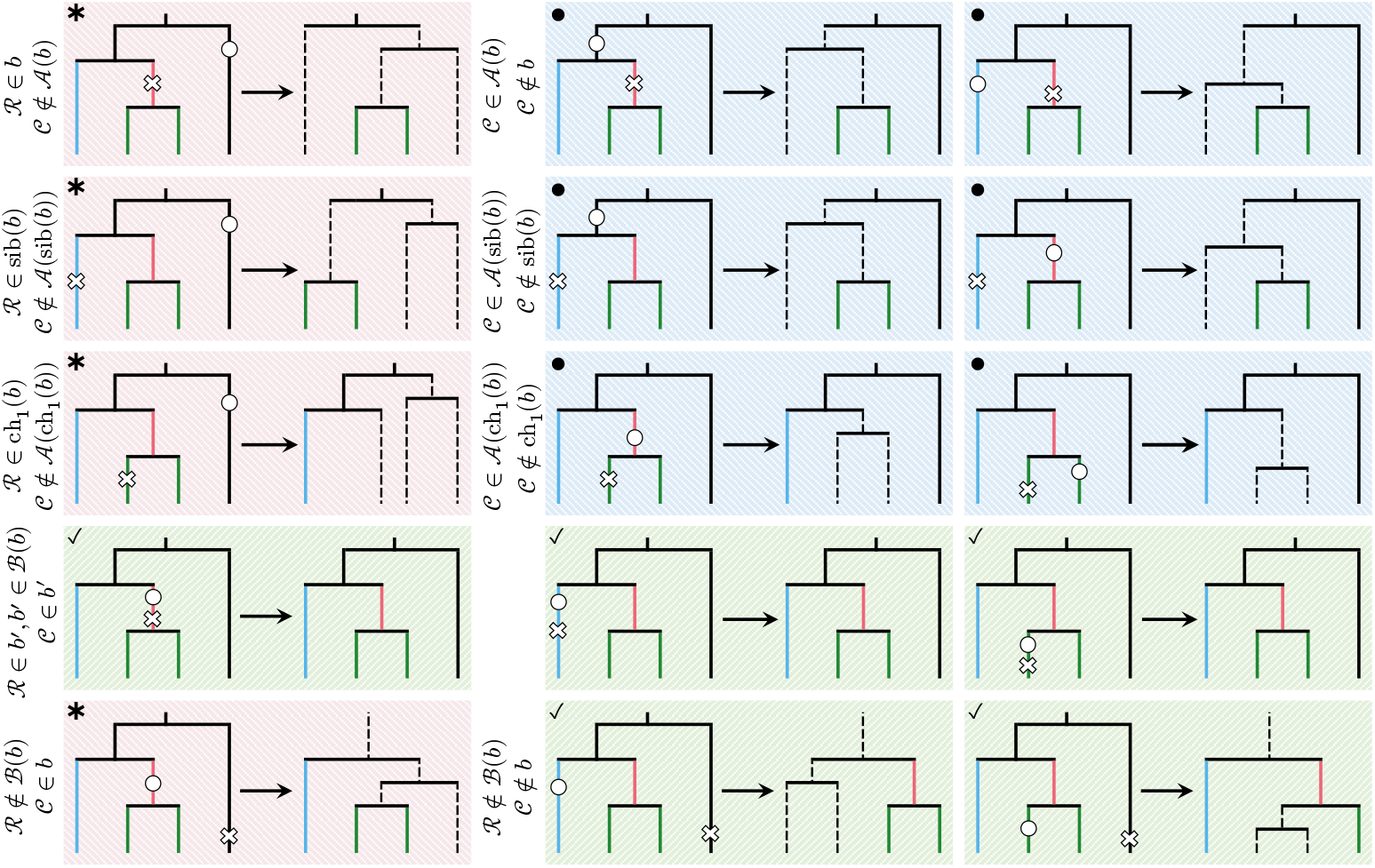
Possible events that do and do not disrupt edge *b* (shown in red); sib(*b*) is shown in blue, ch_1_(*b*) and ch_2_(*b*) in green. Recombination points are shown as crosses; coalescence points as circles. edges that are disrupted (or newly added) are shown as dashed lines. Events highlighted in green (marked with ticks) do not disrupt *b*. Events highlighted in red (marked with stars) disrupt the edge in terms of both edge length and topology (*b* is topologically disrupted); those highlighted in blue (marked with dots) disrupt *b* via changing only its time-length.

We now calculate each of these probabilities in turn for an arbitrary edge *β* ∈𝒯under the SMC’.

### S1.3.1 Probability recombination point is on edge *β*

Under the SMC’, the recombination point is chosen uniformly at random along the edges of the tree. Thus, the probability that the recombination event happens on edge *β* is the ratio of the edge length to the total branch length of𝒯, so

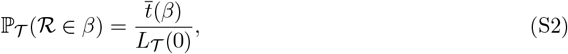

and

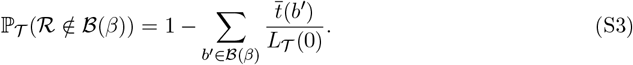

### S1.3.2 Probability coalescence point is not on *β* **given recombination point is on** *β*

The probability that, conditional on the recombination event happening on edge *β* at time *s*, the coalescence point is on *β* has been derived by Deng et al. (2021), which in our notation is as follows.

**Proposition S1.1** (Deng et al. (2021), Proposition 1). *Letting k* = *n*(*s*), *so that T*_*k*_ *is the first coalescence time just above time s*,

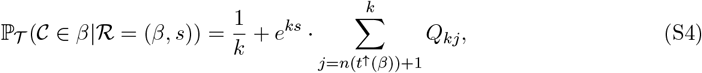

*where*

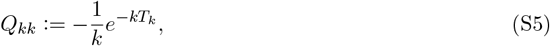

*and*

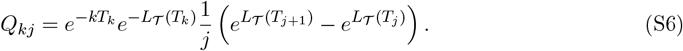

Marginalising out the recombination time *s*, hence summing over *k* = *n*(*s*) in (S4), gives the following.

**Proposition S1.2** (Deng et al. (2021), **Proposition 2)**.

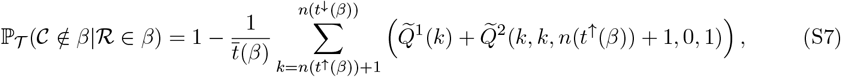

*where*

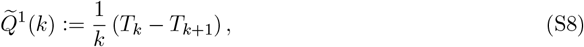

*and for x, y, A, B* ∈ Z, *x* ≥ *k*, 2 ≤ *y* ≤ *x*,

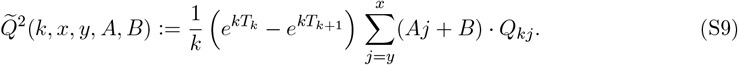

The proofs, translated into our notation, are given in Sections S2.1 and S2.2.

### S1.3.3 Probability coalescence point is on *β* **given recombination point is not on** *B*(*β*) We start by conditioning on the recombination time *s* to obtain the following

**Proposition S1.3**. *Conditional on the recombination point ℛ being at time s and on an edge outside the setℬ*(*β*), *with k* = *n*(*s*), *the probability that β is disrupted is*

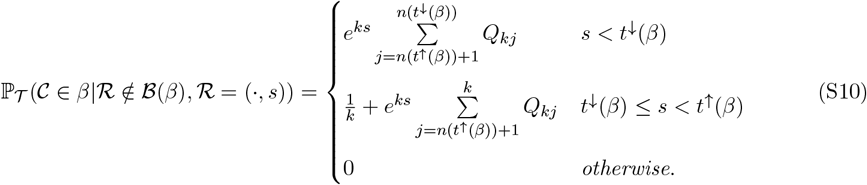

*with Q*_*kk*_ *and Q*_*kj*_ *as defined in* (S5) *and* (S6), *respectively*.

The proof is given in Section S2.3.

Let *t*_1_, *t*_2_, *t*_3_, *t*_4_ denote the event times *t*^↓^(ch_1_(*β*)), *t*^↓^(ch_2_(*β*)), *t*^↓^(sib(*β*)), *t*^↓^(*β*) sorted in increasing order, and define *t*_0_ := 0 and *t*_5_ := *t*^↑^(*β*). Integrating out the recombination time in (S10), we have the following.

**Proposition S1.4**. *Conditional on the recombination point being on an edge outside the set B*(*β*), *the probability that β is disrupted is*

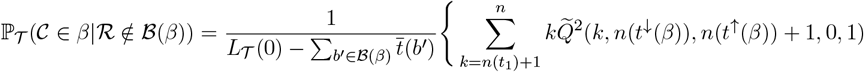

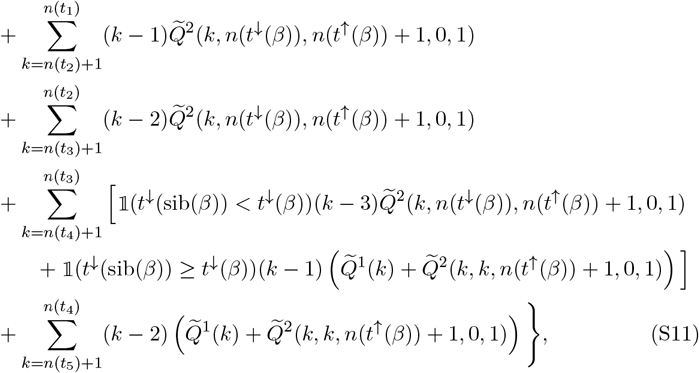

*with*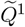*and* 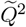*as defined in* (S8) *and* (S9), *respectively*.

The proof is given in Section S2.4.

Substituting the expressions (S2), (S3), (S7) and (S11) into (S1) gives the desired probability that edge *b* is disrupted by the next recombination event.

### S1.4 Probability that an edge is topologically disrupted by a recombination event

Most ARG reconstruction algorithms focus on identifying the presence of recombination events through finding patterns of mutations not consistent with tree-like evolution. This, in general, does not allow for the detection of recombination events that only change edge lengths (panels highlighted in blue in Figure S2). We therefore also calculate the probability that an edge *b* is topologically disrupted (corresponding to panels highlighted in red in Figure S2).

**Theorem S1.1**. *The probability that an edge b is topologically disrupted by a recombination event is given by*

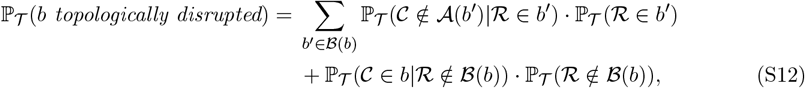

*with*

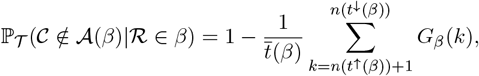

*where, for k* ≤ *n*(*t*^↓^(sib(*β*))),

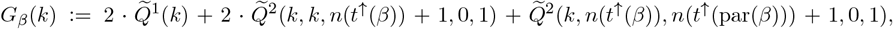

*and for k > n*(*t*^↓^(sib(*β*)))

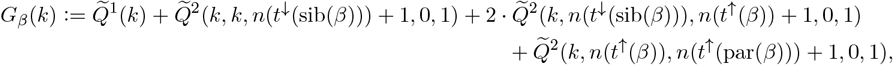

*and*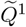*and*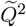*are as defined in* (S8) *and* (S9), *respectively*.

The proof is given in Section S2.5.

### S1.5 Probability that a clade is disrupted by a recombination event

We now calculate the probability that a particular *clade* of edges is disrupted by the next recombination event, i.e. that the membership of sample nodes in the clade changes from one local tree to the next (but allowing for events that disrupt edges within the clade without changing the group of subtended samples). This can happen when a lineage within the clade recombines and coalesces outside the clade or its root edge, or if a lineage from outside the clade recombines and coalesces into the clade, as illustrated in Figure S3.

**Figure S3:**
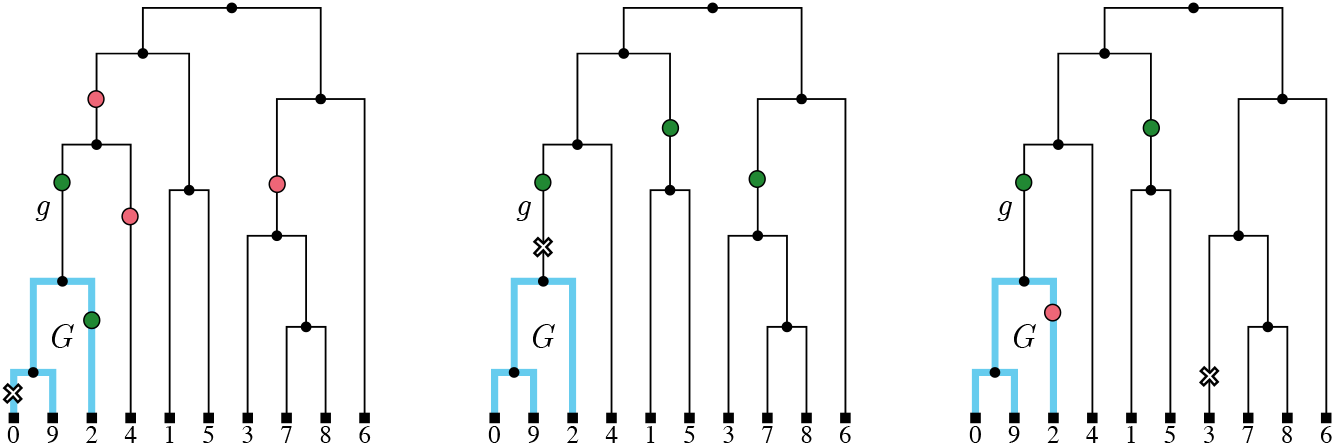
Recombination events that do and do not disrupt a clade. Clade *G*, subtended by edge *g*, contains samples {0, 2, 9}; edges belonging to *G* are shown in blue. In each tree, for the given recombination point (marked by a cross), red (resp. green) circles show examples of coalescence points that would (resp. would not) result in *G* being disrupted.

Let *G* be the set of edges subtended by an edge *g*, with clade MRCA time *t*^↓^(*g*) = *T*_*m*_, *n*_*G*_(*t*) the number of lineages in clade *G* at time *t*, abusing notation to write *G* ∪ *g* := *G* ∪ *{g}*. Let *n*_*G*∪*g*_(*t*) be the number of lineages in *G* ∪ *g* at time *t* and *L*_*G*_(*t*) the total branch length within the clade above time *t*. Then

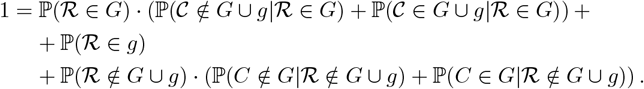

Considering only the events that disrupt the clade, we obtain

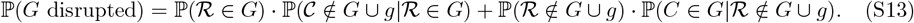

**Theorem S1.2**. *The probability that a clade G is disrupted by a recombination event is*

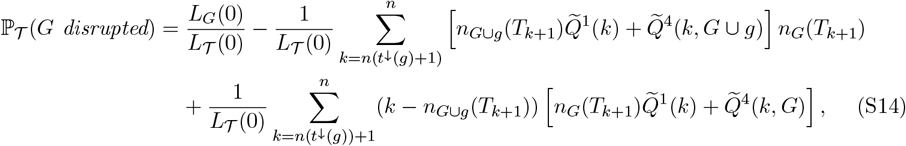

*where*

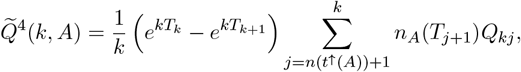

*taking t*^↑^(*G* ∪ *g*) = *t*^↑^(*g*) *and t*^↑^(*G*) = *t*^↓^(*g*).

The proof is given in Section S2.10.

### S1.6 Change in total branch length of tree following a recombination event

Conditioning on the tree𝒯, we now consider the distribution of

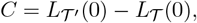

the change in total branch length following a single recombination event. First, considering the sign of the change, we have the following.

**Proposition S1.5**. *The probability of C being negative, zero, or positive is given by*

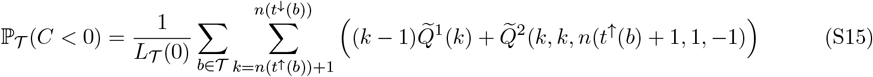

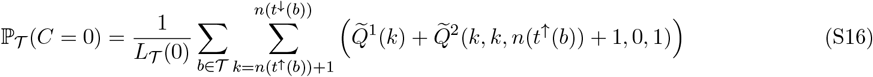

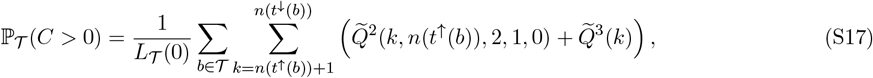

*respectively, where*

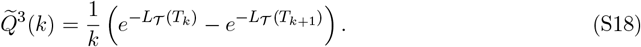

The proof is given in Section S2.6.

To explore the distribution of the magnitude of the change in edge length (when this is non-zero), we derive an approximation of its density.

**Proposition S1.6**. *Conditional on the change in total branch length being non-zero, the density of C is given approximately by*

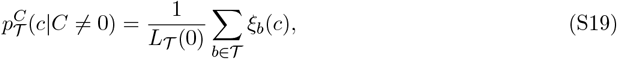

*where ξ*_*b*_(*c*) *is given by*

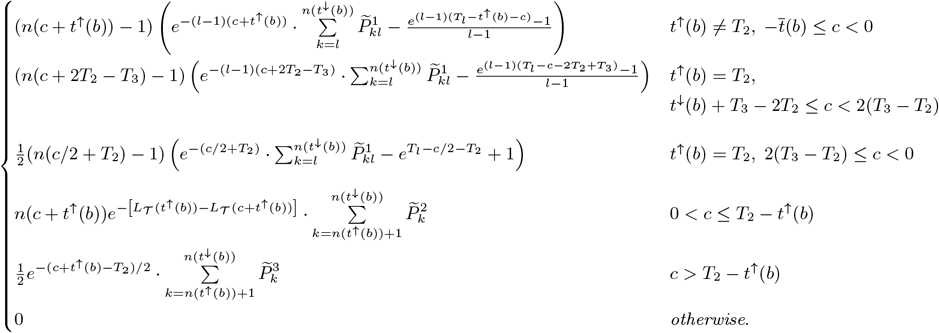

*and*

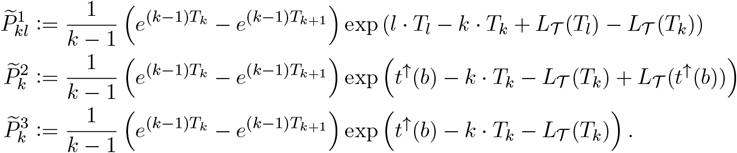

The proof is given in Section S2.7. This is an approximation rather than an exact result under the SMC’, since it assumes that after conditioning on the coalescence point not being on the same branch as the recombination point, the coalescent dynamics follow the SMC model (we find this to give a very close approximation, and simplifies our calculations).

### S1.7 Change in tree height following a recombination event

The height of the tree, *H*(𝒯) = *T*_2_, can change following a recombination event if (1) the coalescence point is above *T*_2_, in which case a new root is formed and the tree height increases, or (2) the recombination happens on one of the two lineages descending from the MRCA, then the tree height can either increase or decrease. Let the setℳ := *{*ch_1_(*b*_*r*_), ch_2_(*b*_*r*_)*}* contain the two edges descending from the MRCA, and let *H* = *H*(𝒯 ′) − *H*(𝒯) be the magnitude of the change in height.

Then we have the following.

**Proposition S1.7**. *Conditional on* 𝒯, *the probability of the change in tree height being negative, zero, or positive is given by*

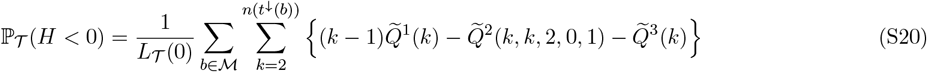

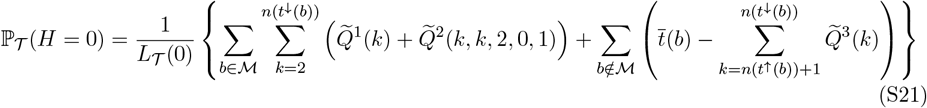

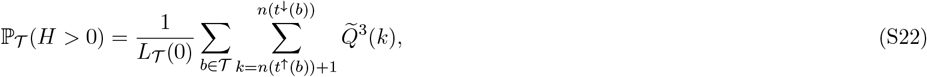

where 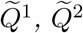 and 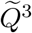*are as defined in* (S8), (S9) *and* (S18), *respectively*.

The proof is given in Section S2.9.

### S1.8 Distribution of edge span

Under the SMC’, for a given edge *b*, the distribution of its span can be characterised by considering the rate at which edge-disrupting recombination events arrive as we move left-to-right along the genome. However, the instantaneous rate at which *b* is disrupted at position *τ* may not be the same as that at position *τ* ′ *> τ*, due to the effect of other recombination events that might occur between *τ* and *τ* ′. Thus, the span of an edge is the waiting time to the next edge-disrupting recombination event, but the rate at which this happens is inhomogeneous along the genome (and is, in fact, itself random).

Similarly to Deng et al. (2021), however, we find that if a recombination event between adjacent trees𝒯 and𝒯 ′ does not disrupt *b*, then

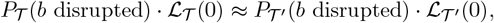

where *P*_𝒯_ (*b* disrupted) is given by (S1), based on simulation results (Section S1.11). Thus, an approximation to the distribution of edge span can be constructed by assuming that the rate at which edge-disrupting recombination events arrive *is* homogeneous along the genome, which is equivalent to assuming that recombination events that do not disrupt the edge *b* also do not change the local tree: so if is the local tree at position *d*^←^(*b*), after each recombination event the newly formed local tree is𝒯 ′ =𝒯. Recombination events occur as a Poisson process along the genome with rate ℒ_𝒯_ (0) *· ρ/*2, allowing us to thin the process by multiplying this rate by the probability that the event is edge-disrupting, thereby offering a tractable approximation for the arrival of edge-disrupting recombination events. Then conditional on *d*^←^(*b*), the edge span *d*^→^(*b*) − *d*^←^(*b*) is distributed as the waiting time to the first event in a Poisson process with rate

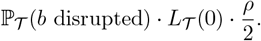

That is,

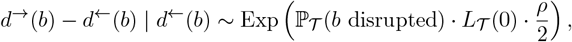

or, by rescaling,

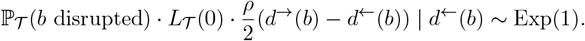

Analogously, if the recombination rate is not constant along the genome, with the population-scaled recombination rate at position *w* given by *ρ*(*w*)*/*2, then the intensity of the process at position *w* is instead given by

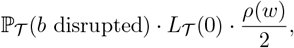

and we have

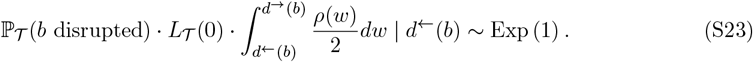

The quality of this approximation can be verified by simulation, using the probability integral transform as follows. For the *i*-th edge *b*_*i*_ ∈ *{b*_1_, …, *b*_*m*_*}* of a simulated ARG, take 𝒯 to be the local tree at position *d*_←_ (*b*_*i*_), compute *q*_*i*_ as

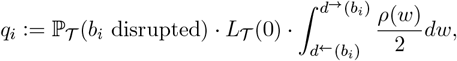

and let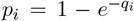. Then a Q-Q plot can be constructed by plotting the ordered quantities *p*_(1)_ ≤ … ≤ *p*_(*m*)_ against the corresponding quantiles of the uniform distribution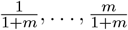. If the approximation fits well, the points should lie on the diagonal. A Kolmogorov–Smirnov (K–S) goodness of fit test can be used to test the null hypothesis that the computed *p*_*i*_ values are uniformly distributed.

We note that if any specific edge at a given position along the genome is selected, it may seem that its genomic span should be the sum of the waiting times to the left and to the right of the given position. This is the well-known “waiting time paradox” and we refer to Feller (1971, p. 12) for a thorough explanation.

### S1.8.1 Considering only topology-disrupting events

If we were to consider only events that topologically disrupt the edge, we instead have the approximation

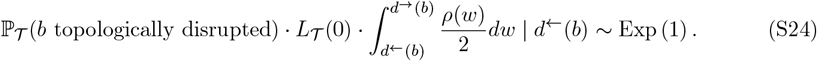

A similar procedure to that described above can be used to check goodness of fit.

### S1.8.2 Conditioning on edge having at least one mutation

ARG reconstruction algorithms utilise mutations to infer changes in local tree topologies due to recombination, so it may be of interest to consider only edges in reconstructed ARGs that are supported by at least one mutation. Suppose that mutations occur as a Poisson process along the edges with constant rate *θ*, and the recombination rate at position *x* is *ρ*(*x*)*/*2. Conditional on the left endpoint of the given edge *d*^←^, let *D* be its right endpoint, which using (S24) has the density

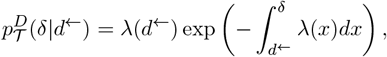

where

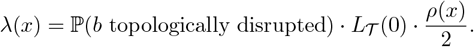

Let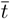 be the time-length and *M* the number of mutations on the edge. Then the conditional distribution of *D* is given by

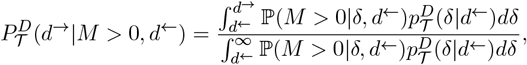

with

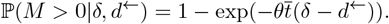

We have

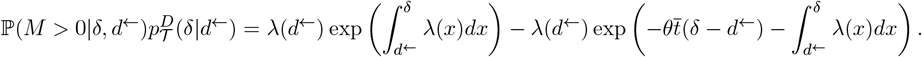

Assuming that the recombination map is piecewise constant, we split the part of the genome to the right of *d*^←^ into portions where the recombination rate is constant between the (ordered) breakpoints *d*^←^ =: *w*_0_ *< w*_1_ *< w*_2_ *<* … *< w*_*k*_ *<* …, adding an extra breakpoint *w*_*k*_ := *d*^→^. Then we can write

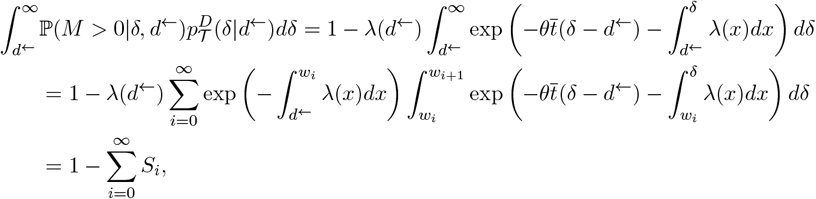

where, by integrating,

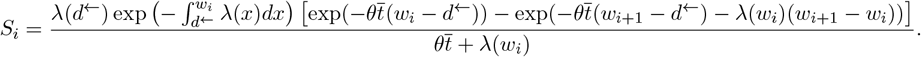

Similarly,

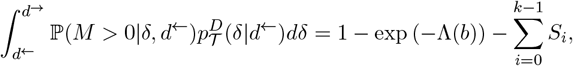

where

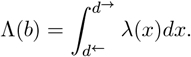

Combining, we have

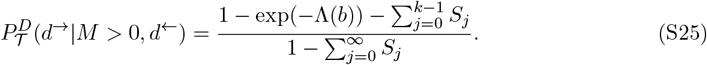

Note that in the limit *θ* → ∞, this reduces to 1 − exp (−Λ(*b*)), as expected (since this effectively removes the conditioning).

For the case where the recombination rate is constant along the genome, with *ρ*(*x*) = *ρ* and *λ*(*x*) = *λ*, we obtain

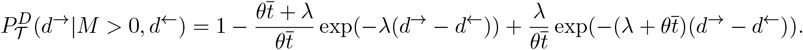

The *p*-values for each edge of a simulated ARG can now be computed by evaluating the cdf (S25) for the given values of *d*^←^ and *d*^→^, and the Q-Q plot can again be constructed by plotting these against the corresponding quantiles of the uniform distribution. However, this potentially requires summing over a large number of increments of the recombination map. Instead, we propose to approximate (S25) by

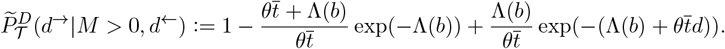

We find this to be a very close match to the exact distribution based on simulations with human-like parameters, while being very fast to compute.

### S1.9 Distribution of clade span

Similar approximations to those employed when investigating the distribution of edge span along the genome can be used for the distribution of the waiting time until a clade *G* is broken up by a recombination event (that is, when the clade either gains or loses one or more samples as a consequence of recombination). Consider an inhomogeneous Poisson process with intensity at position *w* given by

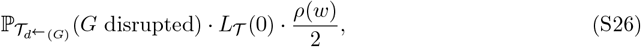

conditional on the local tree at *d*^←^(*G*) (defined as the leftmost position along the genome where the clade arises). Again through rescaling time, we have

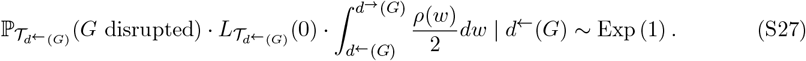

Thus, for each clade *G*^*i*^ in a simulated ARG, we can calculate its left and right endpoints *d*^←^(*G*^(*i*)^) and *d*^→^(*G*^(*i*)^), respectively. Letting

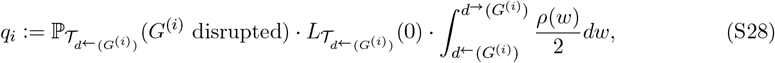

the quality of the approximation can again be checked using a Q-Q plot as described above.

### S1.9.1 Adjusting for varying population size

Given a population size function *N* (*t*), *t* ≥ 0, let

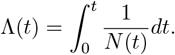

Let 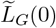 be the total length of branches in *G* measured in generations, and 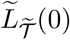 the total branch length of𝒯 measured in generations. Then we have

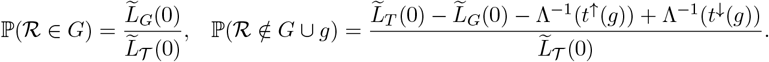

Conditional on the recombination point being on a branch within *G*, the density of the recombination time is

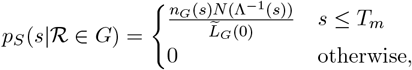

and similarly,

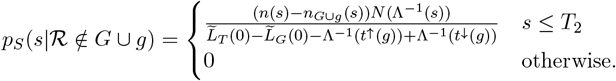

Suppose now that *N* (*t*) is piecewise constant, and for each 2 ≤ *k* ≤ *n*, write

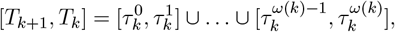

where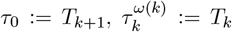, and *N* (*τ*) =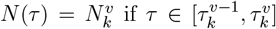. That is, *ω*(*k*) is the minimal number of (disjoint) intervals where the population size is piecewise constant, while there are *k* lineages in the tree. Then following similar calculations as in the proof of Theorem S1.2, we have

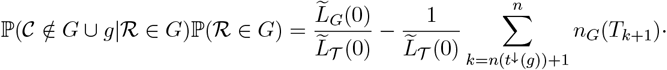

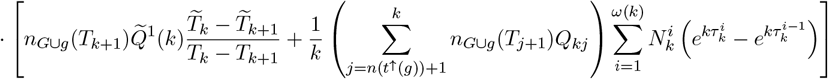

where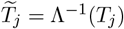, and

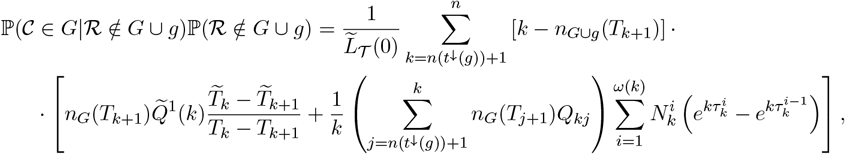

which can be substituted into (S13), and in turn into the expressions in Section S1.9, to give the corresponding approximation to the genomic span of *G* under an arbitrary piecewise constant population size model. This can also be applied to an arbitrary population size model, through averaging the population size over a suitable time grid and thus approximating it with a piecewise constant function.

### S1.10 Quality of approximation to the distribution of edge span

We first assess the quality of the approximation derived in Section S1.8 by simulating an ARG under the SMC’ and checking if the simulated edge spans follow (S23), by using the procedure described in Section S1.8. The simulation parameters are given in Section 4.6.1 (main text), and we sampled 10 000 edges from each ARG (uniformly at random) for testing, to speed up computation. The corresponding Q-Q plots are shown in Figure S4 (blue points). The points adhere very closely to the diagonal, demonstrating that the approximation provides an excellent fit. The K–S *p*-values of 0.31 (left panel) and 0.75 (right panel) also suggest good agreement. Grouping edges by their depth (the number of edges on the way to the MRCA) or clade size (number of samples subtended by the edge) in the tree and constructing Q-Q plots for each group also did not reveal significant deviation from the diagonal (Figure S14), suggesting that the approximation holds for all edges.

**Figure S4:**
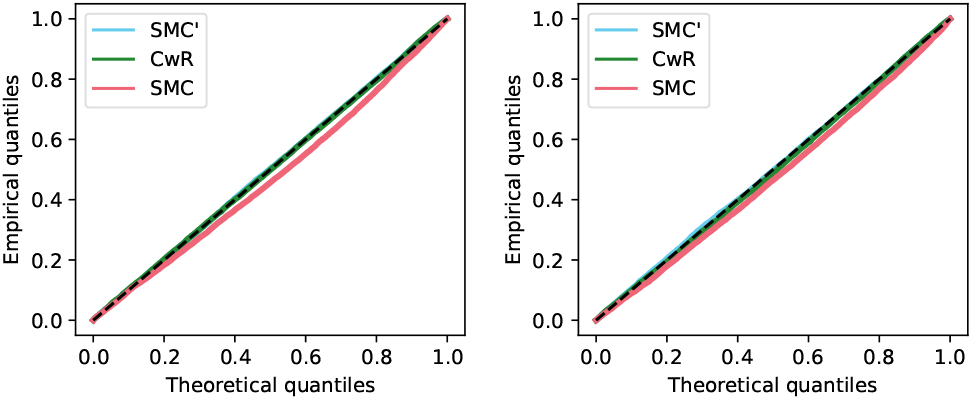
Q-Q plots using (S23) computed from ARGs simulated using the SMC’ (blue), CwR (green), and SMC (red) models with *n* = 100. Note the blue and green points closely overlap, and overlay the diagonal. Left panel: dataset 1 parameters; right panel: dataset 2 parameters. Dashed line: diagonal from (0, 0) to (1, 1).

### S1.11 Effects of recombination on a local tree

The very good quality of the approximation above can be understood by considering the effects of a recombination event on properties of the local tree.

For a given edge *b*_*i*_ that exists in local trees 𝒯_(1)_, …, 𝒯_(*k*)_, let

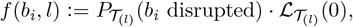

for 1 ≤ l ≤ k. Let

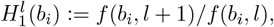

for 1 ≤ l ≤ k ™ 1, and

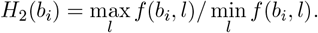

We calculate these quantities for a uniform random sample of 1 000 edges from an ARG simulated using dataset 1 parameters in Section 4.6.1 (Main Text); the corresponding histograms are shown in Figure S5. These suggest that the quantity *P*_𝒯_ (*b*_*i*_ disrupted) *·* ℒ_𝒯_ (0) stays relatively conserved following each recombination event, even if individually *P*_𝒯_ (*b*_*i*_ disrupted) and ℒ_*>*𝒯_ (0) may vary more significantly. This is similar to the findings of Deng et al. (2021, Figure 5).

**Figure S5:**
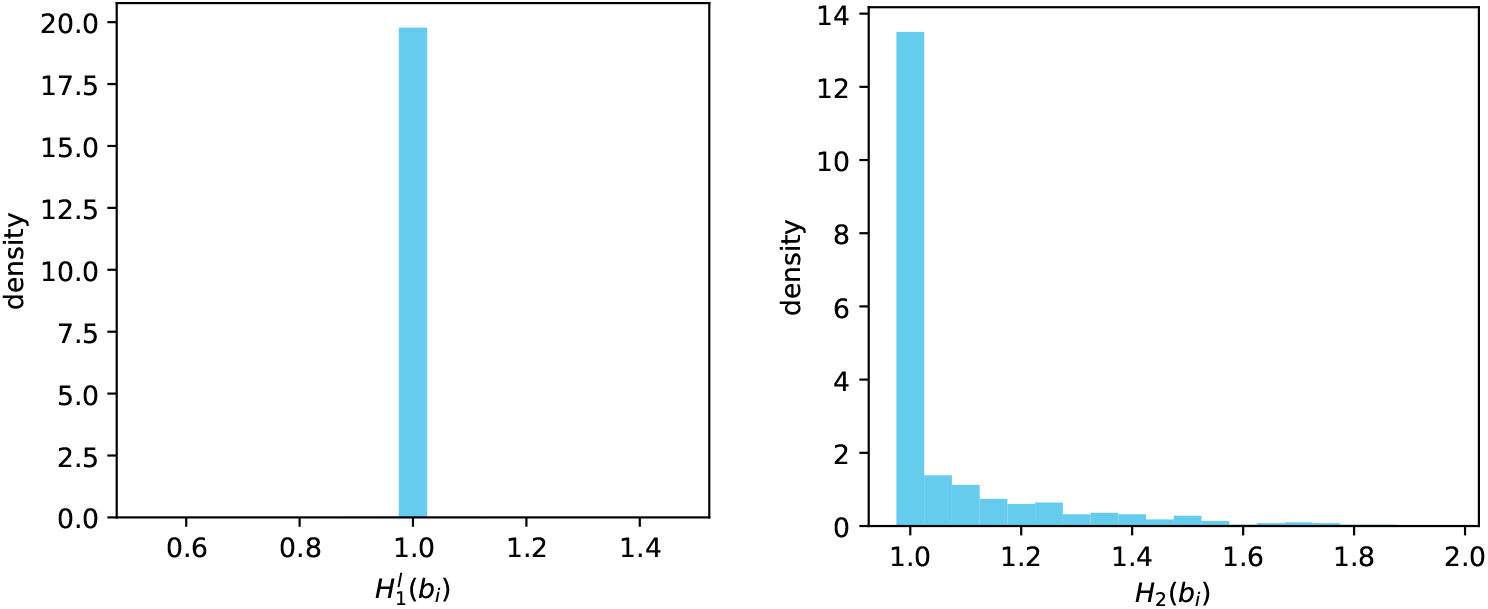
Histograms of *H*^*l*^ (*b*_*i*_) and *H*_2_(*b*_*i*_) for a simulated ARG.

To calculate a Monte Carlo estimate of the marginal probability that the change in total branch length is negative, zero, or positive, we average over local trees simulated using msprime under the SMC’ model. The results are shown in Figure S6. Recombination has a stabilising effect on the total branch length of local trees: when the total branch length is small (resp. large), the probability that the recombination event will increase the total branch length increases (resp. decreases).

**Figure S6:**
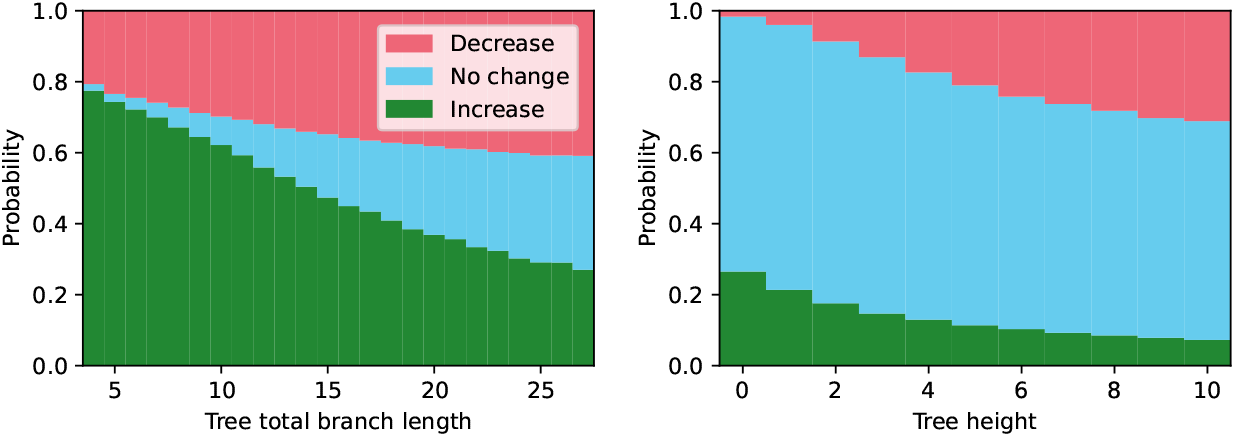
Mean probability of change in total branch length (left) and tree height (right) being negative (red, top stack), zero (blue, middle stack) or positive (green, bottom stack). Trees were simulated and binned according to total branch length (left) or height (right), with 100 trees simulated per bin, and sample size *n* = 100. For each tree, probabilities were calculated using (S15)-(S17), the stacked bar plot shows the mean probabilities for each bin.

Similarly, we can average over trees simulated under the SMC’ model to estimate the marginal probability that the change in tree height following a recombination event is negative, zero, or positive. The results are shown in Figure S6 (right panel). As with total branch length, recombination tends to increase (resp. decrease) the tree height with higher probability when the tree height is small (resp. large).

Figure S7 shows the density (S19) for three simulated trees with varying total branch lengths *L*_𝒯_ (0), for *n* = 10 and *n* = 100. In both cases, the density is concentrated around zero and skewed to the left (resp. right) when the total branch length is small (resp. large); it is roughly symmetric about zero for middling values of *L*_𝒯_ (0).

**Figure S7:**
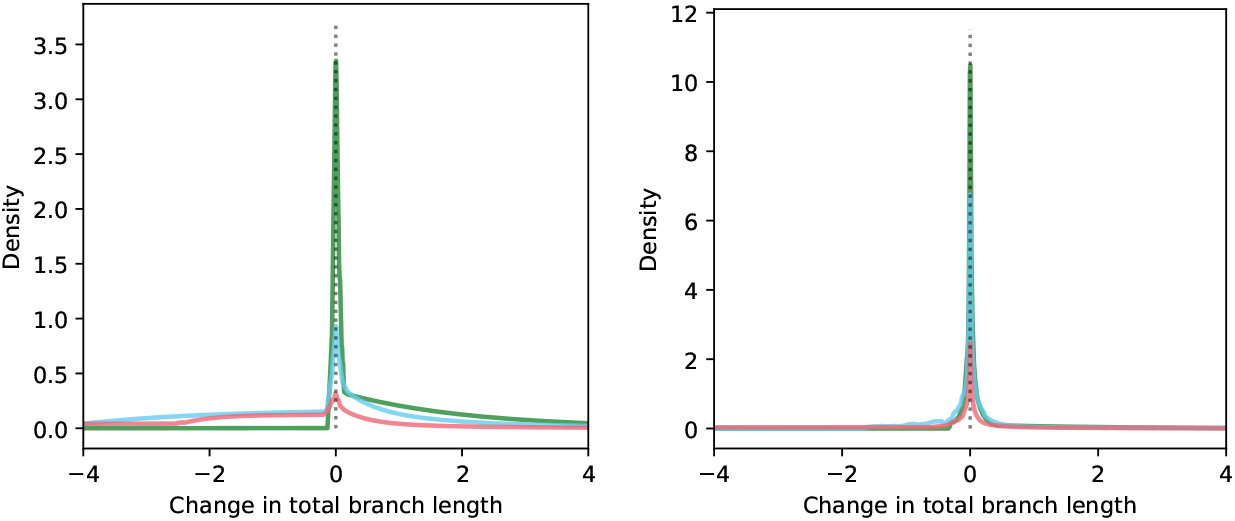
Density of change in total branch length (S19) for three simulated trees, with *n* = 10 (left panel) *n* = 100 (right panel). Left: trees have total branch length 1 (green), 6 (blue) and 24 (red). Right: trees have total branch length 5 (green), 10 (blue) and 28 (red).

These results shed light on why, despite the strong assumption that recombination events that do not disrupt the given edge also do not change the rest of the local tree, our approximation to the distribution of edge span gives an extraordinarily close fit for data simulated under the SMC’ model. For an edge that is close to the leaves, the probability that the edge is disrupted by the next recombination event is very small (Figure 2). Thus, many recombination events will occur before this edge is disrupted. As can be seen in Figure S6, recombination has the effect of stabilising the total branch length, with events causing an increase (decrease) in total branch length being more likely if the current total branch length is relatively small (large). Thus, it seems the fluctuations in total branch length average out and do not significantly affect the overall rate of edge-disrupting recombination events. On the other hand, for an edge that is close to the root of the tree, per Figure 2 the probability of the edge being disrupted by the next recombination is relatively high. Thus, a relatively small number of recombination events are likely to occur before they affect the given edge. As illustrated in Figure S7, when a recombination changes the total branch length of the tree, the magnitude of this change is concentrated around 0. Thus, the effect of recombination on the rest of the tree does not appear to significantly affect the probability that the edge is disrupted.

This applies at the level of each individual edge, so after rescaling each observed edge span by its specific event rate as per (S23), these rescaled edge spans follow an Exp(1) distribution. Accounting for mutiple testing using a Bonferroni correction, we can thus use the resulting *p*-values to detect outlier edges with longer-than-expected spans. The same reasoning applies for the genomic spans of clades.

### S1.12 Comparison of simulation models

We next simulate ARGs under the CwR and under the SMC, with the same two parameter settings given in Section 4.6.1 (main text) and again compare the resulting edge spans to (S23). For the CwR, the span of an edge is taken to be the sum of all the genomic intervals where that edge appears in the local tree (to account for the presence of recombination events that occur in non-ancestral material). Figure S4 shows that the approximation is an excellent fit to the CwR (green points), with K–S *p*-values of 0.06 (left panel) and 0.94 (right panel). This suggests that the distribution of edge spans under the SMC’ and that under the CwR are remarkably close.

Under the SMC (red points), edge spans are shorter in general, with points falling below the diagonal (with both K–S *p*-values *<* 0.001). This is due to the model disallowing recombination events that do not change the local tree, so edges are more frequently disrupted by recombination.

### S1.13 Detection of local recombination suppression: Test 1

Given an ARG, for each clade *G*^(*i*)^, we calculate its left and right endpoints *d*^←^(*G*^(*i*)^) and *d*^→^(*G*^(*i*)^), and would like to estimate the probability of observing a clade span greater than *d*^→^(*G*^(*i*)^) − *d*^←^(*G*^(*i*)^) via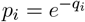, with *q*_*i*_ as defined in (S28). This is a one-sided *p*-value, and we test whether *G*^(*i*)^ has a significantly longer span than otherwise expected by comparing this against a Bonferroni-corrected significance threshold (0.05 divided by the total number of tested clades). Simulation studies confirm that for ARGs simulated under the SMC’ model without inversions, these *p*-values are approximately uniformly distributed (Figure 4, left panel, blue points), as expected.

### S1.13.1 Reconstructed ARGs

As can be seen from Figure 4 (left panel), similarly to edge span, the distribution of clade span in ARGs reconstructed using Relate, tsinfer/tsdate and ARG-Needle is skewed, with clade span generally overestimated by these methods. Thus, directly applying the test as described above will lead to a high false positive rate. We now describe a correction which can be applied to counteract two main problematic features of reconstructed ARGs, focusing particularly on Relate (due to the presence of polytomies for tsinfer/tsdate being difficult to correct for, and the large bias seen with ARG-Needle which both under- and overestimates clade span).

Let *G* = *{G*_1_, *G*_2_, …, *G*_*N*_ *}* be a list of all clades in the reconstructed ARG. The first issue is that due to a lack of mutations around the leftmost and rightmost endpoints of a clade, Relate may overestimate its span, causing false positives. To correct for this, we proceed as follows. Suppose that the root edges of a clade *G*^(*i*)^ in trees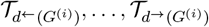have mutations at positions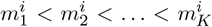. We (1) remove from *G* all clades with fewer than three mutations in total and fewer than *M* mutations per kb on average, and (2) measure an adjusted clade span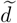 using the positions of the leftmost and rightmost mutations that support the given clade. That is, we define

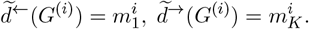

The second issue is that the clade carrying the inversion may not be supported by mutations uniformly along the inverted region, causing it to appear and disappear multiple times in quick succession in the reconstructed ARG, which can cause false negatives. We correct for this by “merging” pairs of clades that are nearby on the genome (in terms of genetic distance, to allow for varying recombination rates). For two clades *G*^(*i*)^, *G*^(*j*)^ ∈ *G* that have identical sets of sample descendants and are less than *L* cM apart, that is

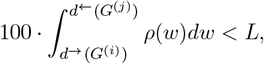

form *G*_*i,j*_ := *G*^(*i*)^ but setting 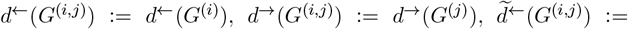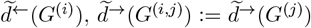, and update *G* as

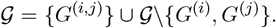

We apply this to all pairs of clades in *G* iteratively, until no more clades can be merged together. We note that this correction also helps to handle the presence of gene conversion within inverted regions, which can be commonplace (Korunes and Noor, 2019; Crown et al., 2018): a gene conversion will result in a short stretch of the genome where the clade is disrupted but then reappears, and the described adjustment will ensure that this does not affect the calculated clade span.

For each clade in the reduced list *G*^(*i*)^ ∈ 𝒢, we thus calculate an adjusted version of (S28):

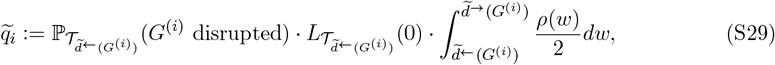

again taking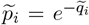. Note that with the above definitions, this can be computed even though the clades in the reduced list will now not necessarily exist (with the re-defined spans) in the ARG itself. We thus obtain adjusted *p*-values for each clade, applying a significance threshold of 0.05*/N*, where *N* is the original number of clades in the reconstructed ARG.

We apply these corrections to the ARG reconstructed using Relate for the data simulated using SLiM (as described in Section 4.6.2), setting *L* = 0.01 and *M* = 0.05. The resulting Q-Q and *p*-value plots are shown in the top row of Figure S18, showing that the correction brings the points on the Q-Q plot very close to the diagonal, and there are three significant clades (all of which overlap the inverted region, and the clade spanning the entire inverted region remains a significant outlier with the lowest *p*-value). The equivalent plots for a simulation with no inversion are shown in the bottom row of Figure S18, demonstrating that the *p*-values are approximately uniformly distributed and there are no false positives.

The choice of the parameter *L* influences power and the rate of false positives, which will both increase as *L* increases, since the merging procedure lengthens clade spans. We construct a bound on the false positive rate using the following approximation. For a given set of *S* sequences, the probability that these form a clade in a random coalescent tree of size *n* is

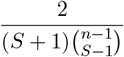

(e.g. Hein et al., 2004, p. 84, eq. 3.26). This probability is very small unless *S* is small or, by symmetry, close to *n*. Thus, for a given clade *G* of size *S*, the probability that the clade is broken up by recombination at position *d*^→^(*G*) but then appears again within *L* cM (purely due to random chance) is bounded above by

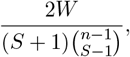

where *W* is the number of trees within *L* cM of *d*^→^(*G*) (and further requiring that the clade is supported by at least one mutation results in a smaller bound).

For the 1KGP data, there are ≈ 2m trees in total along the genome. With *n* = 100, setting *L* = ∞ (so *W* ≤ 2 000 000), gives an upper bound of 2 *·* 10^−7^ on the probability that a clade of size *S* = 10 reappears by random chance anywhere along the genome. Thus, even if 10m clades of size 10 are considered, we expect at most two false positives to arise due to the merging procedure. The rate of false positives decreases as *n* (and *S*) increase; with *n* = 1000 and *S* = 10 the upper bound falls to 1 *·* 10^−16^. In conclusion, choosing *L* to be large only slightly increases the rate of false positives, while increasing power.

### S1.13.2 Test error rates

We examined the performance of the test by applying it to ARGs simulated using SLiM with the parameters given in Section 2.4.1 (with inversions at intermediate frequency, on average 50%), varying the length of the inverted region from 0 to 200kb (100 ARGs in each case). Defining positive detection as there being at least one clade within the inverted region with a significant *p*-value, the resulting ROC curve is shown in Figure S8 (left panel). Fixing the false positive rate at 5% (corresponding roughly to one false positive per 100Mb) gives the confusion matrix shown in Table 1a, demonstrating very high sensitivity. For each inversion length, out of all the clades with significant *p*-values across the simulations, a high percentage lie within the inverted region.

**Table 1.** Confusion matrices and results summaries for inversion detection test, based on 100 simulations for each given length of the inverted region: using (a) the simulated ARGs, (b) ARGs reconstructed using Relate.

| (a) |  | Inversion length (kb) |  |  |  | (b) |  | Inversion length (kb) |  |  |  |
| --- | --- | --- | --- | --- | --- | --- | --- | --- | --- | --- | --- |
|  |  | 200 | 100 | 50 | 0 |  |  | 200 | 100 | 50 | 0 |
| Inv. detected | + | 100 | 97 | 99 | 5 | Inv. detected | + | 93 | 78 | 57 | 5 |
|  | – | 0 | 3 | 1 | 95 |  | – | 7 | 22 | 43 | 95 |
| Clades tested |  | 3.3m | 3.4m | 3.4m | 3.4m | Clades tested |  | 116k | 116k | 117k | 122k |
| Significant clades |  | 201 | 144 | 127 | 5 | Significant clades |  | 227 | 123 | 79 | 5 |
| Within inv. region |  | 96% | 92% | 97% | 0% | Within inv. region |  | 96% | 97% | 96% | 0% |

Reconstructing an ARG using Relate for each simulated dataset and applying the adjustments described in Section 2.4.2 gives the ROC curve shown in Figure S8 (right panel); the results in Table 1b show high sensitivity is maintained for inversions longer than around 100kb. These results demonstrate very good performance in detecting the presence of inversions, as well as pinpointing the candidate clade and its position along the genome.

**Figure S8:**
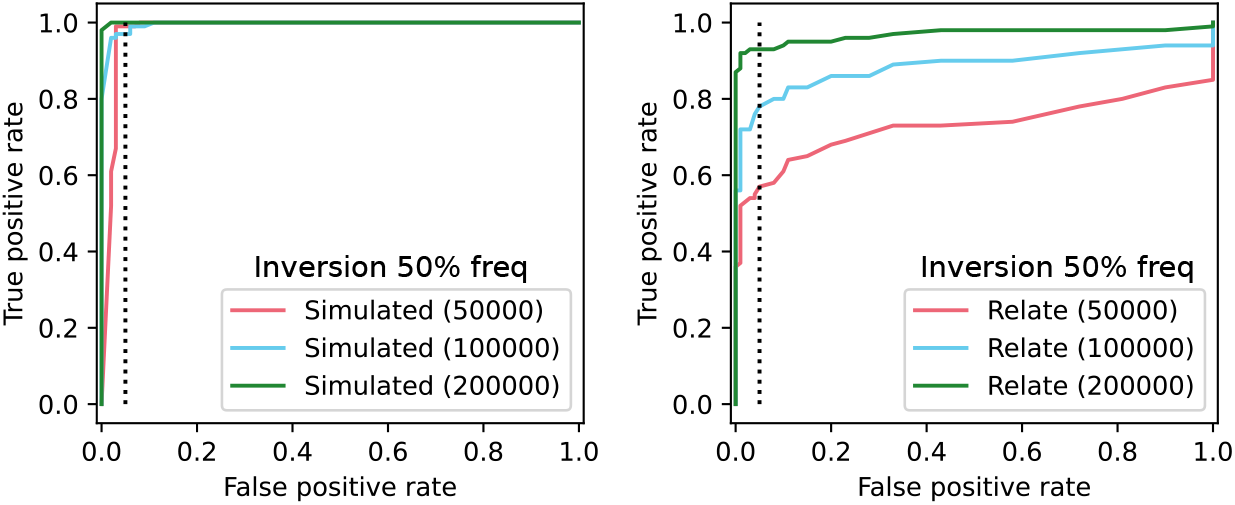
ROC curve for inversion detection test, based on 100 simulations for each given length of the inverted region. Left: using the simulated ARGs; right: ARGs reconstructed using Relate. Dotted line corresponds to a false positive rate of 5% (false positive being defined as an ARG simulated with no inversion but having at least one significant clade in the region). Colours correspond to the different inversion lengths.

To investigate how performance depends on inversion frequency for reconstructed ARGs, we further simulated the same scenario but with the inversion at 10% and 20% average frequency. The resulting ROC curves are shown in Figure S9. As expected, power decreases with decreasing inversion frequency, since smaller clades are expected to have shorter genomic spans, so it is more difficult to detect them as outliers.

**Figure S9:**
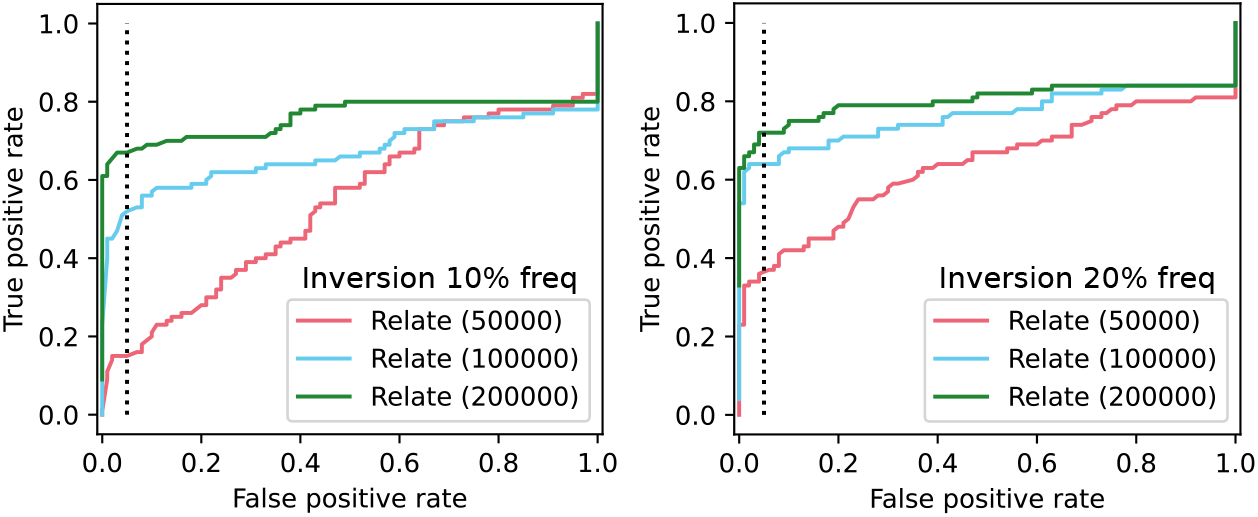
ROC curve for inversion detection test, based on 100 simulations for each given length of the inverted region. Left: inversion at 10% average frequency. Right: inversion at 20% average frequency.

### S1.13.3 Comparison to other methods

We compared the performance of our test in predicting inversion genotypes against that of invClust (Cáceres and González, 2015), a method based on clustering haplotypes using multidimensional scaling of SNPs. We ran simulations using SLiM with varying inversion sizes, as described in Section S1.13.2. Since invClust requires the candidate location of the inversion, we gave the true simulated position as this input (using a larger region containing the inversion gave the same results, and using regions not overlapping with the inversion gave very poor performance, as can be expected). We then used invClust to predict inversion genotypes (homozygous non-carrier, heterozygous, or homozygous carrier) and calculated the squared correlation with the simulated ground truth. We also predicted inversion genotypes using our method, by considering the sequences within the top significant clade. The results are presented in Figure S10 (left panel), showing that our method achieves very high prediction accuracy, consistently outperforming invClust for all simulated inversion sizes.

We also calculated an accuracy score for how well our method predicts the location of the inverted region (by calculating the proportion of overlap between the span of the top significant clade and the true simulated region). A histogram of this is shown in Figure S10 (right panel), demonstrating very good accuracy, with the predicted region overlapping more than half of the true region in 81% of simulations. In both of these comparisons (and in the ROC curves in Figures S8 and S9), it can be seen that the performance of our method improves as the size of the inversion increases.

**Figure S10:**
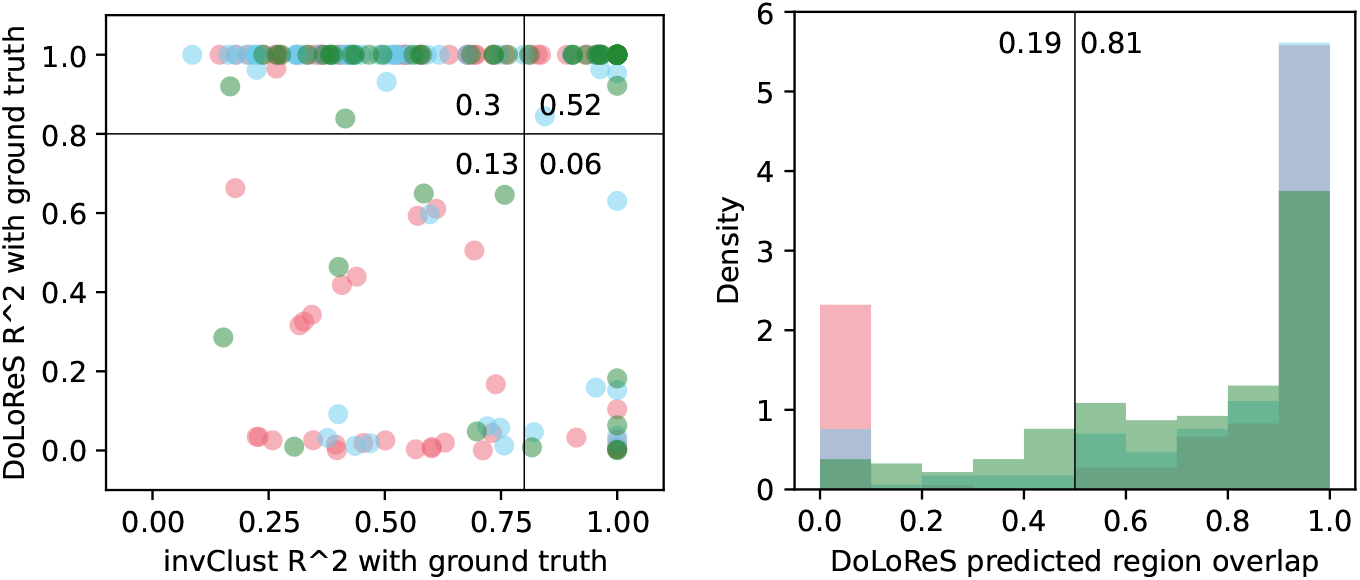
Left panel: comparison of performance against invClust, based on 100 simulations for each given length of the inverted region (colours correspond to region length as in Figure S8). Points show squared correlation between true and predicted inversion carriers. Numbers show proportion of points falling in each quadrant. Right panel: histogram of proportion of overlap between predicted and true inverted region.

We also compared the performance of our method in predicting genotypes and inversion positions against Asaph (Nowling et al., 2022), a method using PCA to detect and localise inversions, but found Asaph to perform very poorly on our simulated data. This is likely because the focus of Asaph is on scalability and the detection of very large and old inversions.

### S1.14 Detection of local recombination suppression: Test 2

Under our approximation, recombination events arrive as an inhomogeneous Poisson process along the genome with rate *λ*(*w*) given by (S26). For a particular clade *G*, call recombination events which do not change the membership of *G* “Type 1”, and other events “Type 2” (our key assumption is that Type 1 events also don’t change the local trees). We thus have Type 1 events arriving at rate *zλ*(*w*), and Type 2 events at rate (1 − *z*)*λ*(*w*), where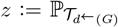(*G* disrupted). Let *D* be the number of Type 1 events before the first Type 2 event. Then it is easy to show that the marginal distribution of *D* is geometric with parameter *z*.

Thus, if *G* is first disrupted by the *R*-th recombination event, we can calculate a corresponding *p*-value as

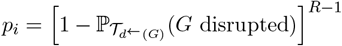

If *R* is known exactly, this is equivalent to Test 1.

## S2 Proofs

### S2.1 Proof of Proposition S1.1

Conditioning on the recombination happening on edge *β*, the density of the recombination event time is

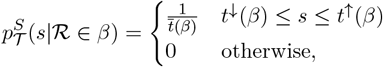

as the recombination time is chosen uniformly at random along the length of the edge. The conditional density of the coalescence time is

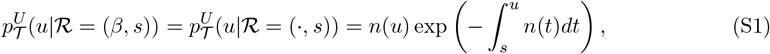

for *u > s* and 0 otherwise. Conditional on the coalescence time *u*, the probability that the coalescence point is on *β* is

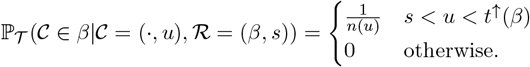

Letting *k* = *n*(*s*), so that *T*_*k*_ is the first coalescence time just above time *s*,

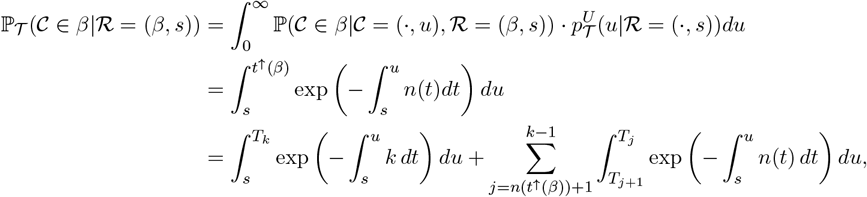

note that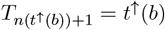. The first term is

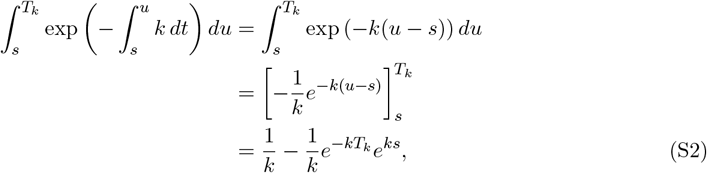

and the summands of the second term are

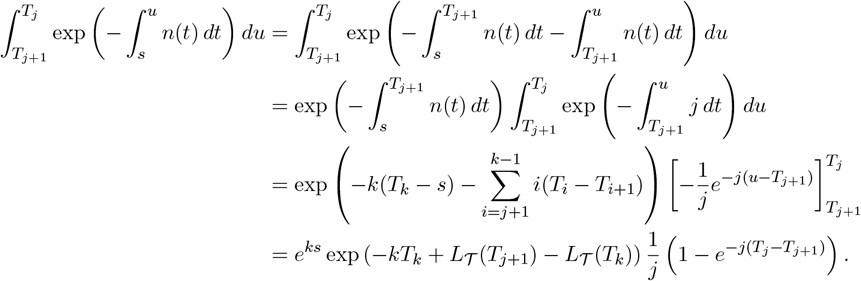

Thus,

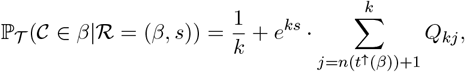

where

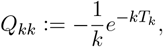

and

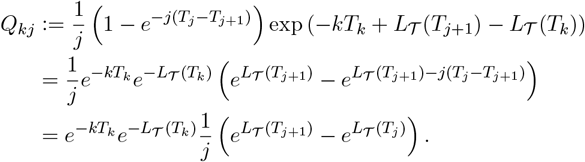

### S2.2 Proof of Proposition S1.2

Marginalising out the recombination time in (S4),

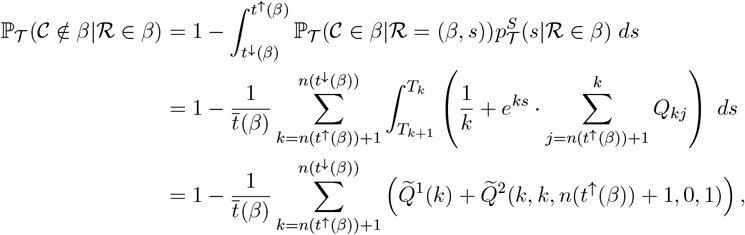

where

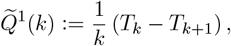

and for *x, y, A, B* ∈ Z, *x* ≥ *k*, 2 ≤ *y* ≤ *x*,

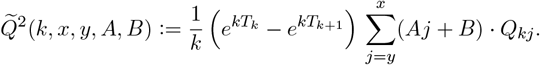

### S2.3 Proof of Proposition S1.3

We have

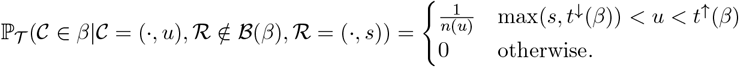

For *s < t*^↓^(*β*),

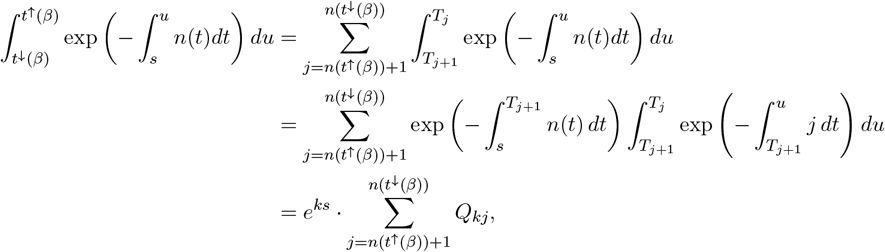

and the case *s* ≥ *t*^↓^(*β*) is given by (S2). Thus,

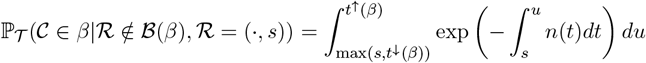

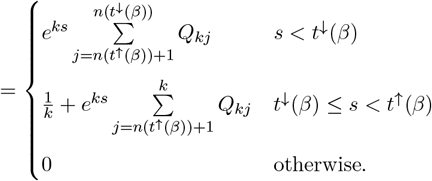

### S2.4 Proof of Proposition S1.4

Consider all of the possible orderings of the event times *t*_1_, …, *t*_4_, as illustrated in Figure S11.

**Figure S11:**
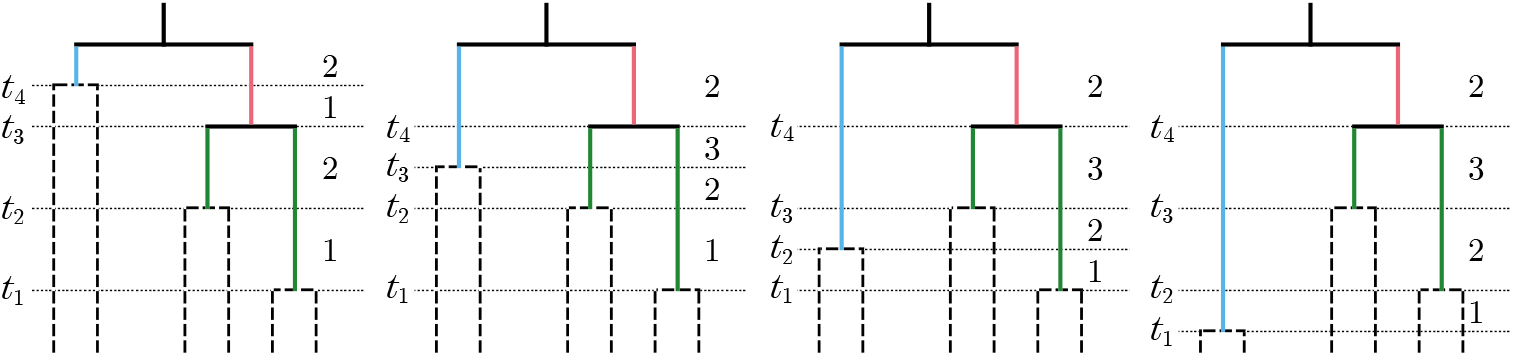
All possible orderings of *t*↓(ch1(*β*)), *t*↓(ch2(*β*)), *t*↓(sib(*β*)), *t*↓(*β*). The edge β is shown sib(*β*) in blue, ch_1_(*β*) and ch_2_(*β*) in green. Numbers to the right of each tree show the number of lineages in (*β*) in each time interval. For instance, in the leftmost tree, *t*_1_ = *t*^↓^(ch_2_(*β*)), *t*_2_ = *t*^↓^(ch_1_(*β*)), *t*_3_ = *t*^↓^(*β*) and *t*_4_ = *t*^↓^(sib(*β*)).

The number of lineages in setℬ(*β*) at time *s* can be written as

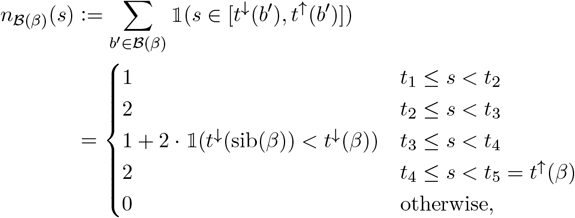

where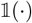 is the indicator function. Then conditional on the recombination point not being on an edge in the set ℬ(*β*), the density of the recombination event time is

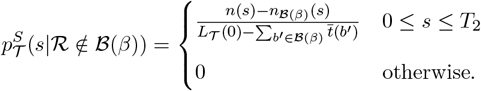

Marginalising out the recombination time in (S10),

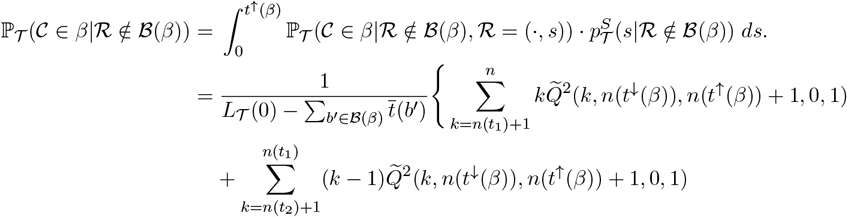

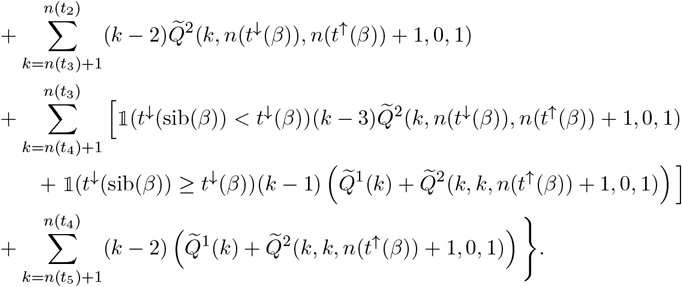

### S2.5 Proof of Theorem S1.1

If we now say that an edge is disrupted only if there is a change in topology (but not edge length), we can allow the events shown in Figure 2 in blue (marked with dots), i.e. those where the recombination point is on edge *β* ∈ *B*(*β*), and the coalescence point is on one of the edges in 𝒜(*β*). Thus,

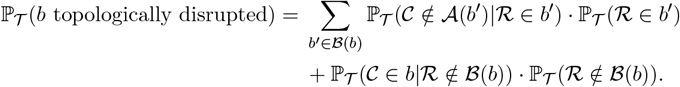

To calculate the probability ℙ_*𝒯*_ (𝒞 ∉*𝒜*(*β*)*|*ℛ∈ *β*), we follow the same approach as the proofs of Propositions S1.1 and S1.2, first conditioning on the recombination pointℛ = (*β, s*). Let *k* = *n*(*s*), so that *T*_*k*_ is the first coalescence time just above time *s*. Let

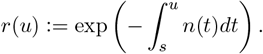

Then if *t*^↓^(sib(*b*)) *< s*,

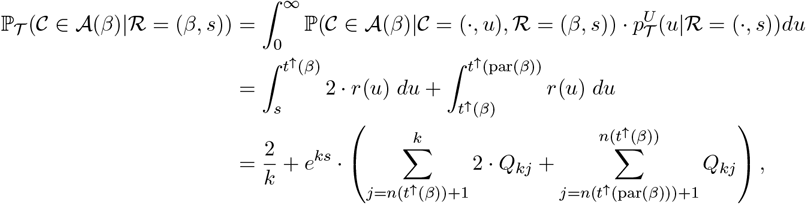

and if *t*^↓^(sib(*b*)) ≥ *s*,

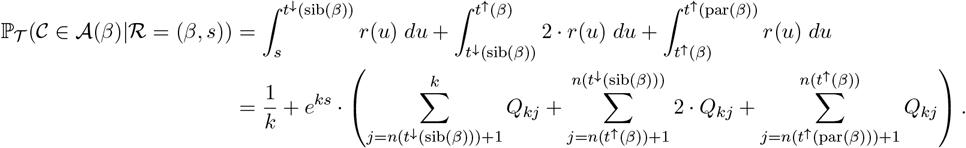

Marginalising out the recombination time,

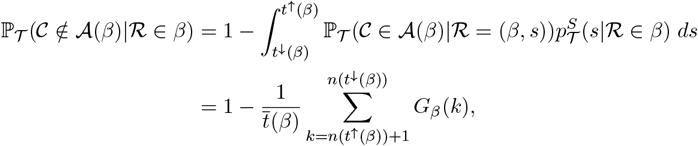

where for *k* ≤ *n*(*t*^↓^(sib(*β*))),

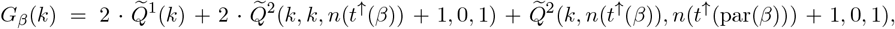

and for *k* > *n*(*t*^↓^ (sib(β)))

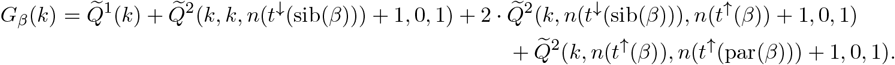

### S2.6 Proof of Proposition S1.5

#### S2.6.1 Probability of no change in total branch length

The probability that the recombination event does not result in a change in total branch length is

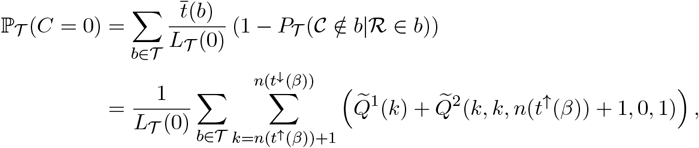

using (S7). This is equal to the probability derived by Deng et al. (2021, Theorem 1).

### S2.6.2 Probability that change in total branch length is negative

We have

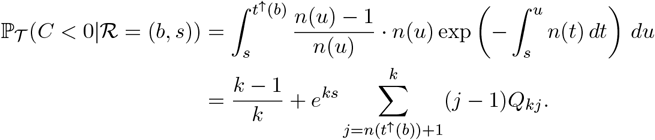

Integrating over the recombination time,

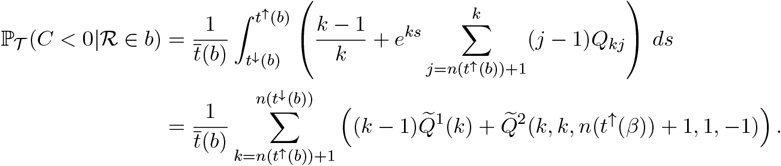

Then

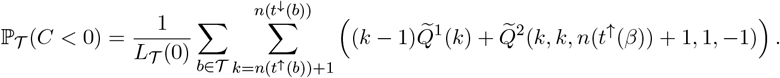

#### S2.6.3 Probability that change in total branch length is positive

Similarly, the probability that the change in total branch length is positive is given by

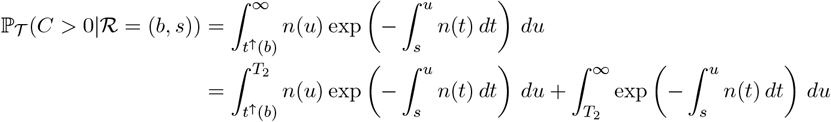

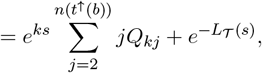

giving

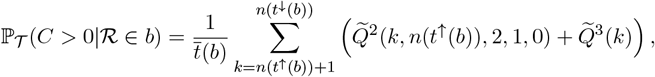

where

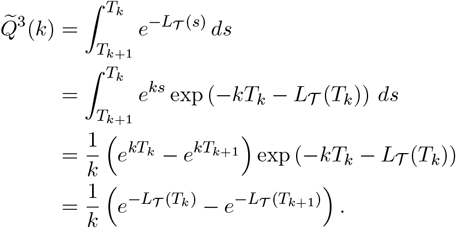

Thus,

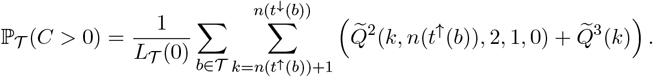

### S2.7 Proof of Proposition S1.6

We first calculate the density of change in total branch length conditional on the recombination pointℛ = (*b, s*), then marginalise out the recombination time and edge.

### S2.8 Density of change in total branch length conditional on recombination point

Suppose that the recombination point is on edge *b* at time *s*. Then given the coalescence time *u*, the change in total branch length *C > t*^↓^(*b*) + *T*_3_ − 2*T*_2_ given *C* ≠ 0 is

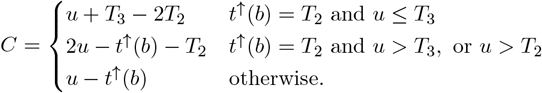

Let *l* = *n*(*u*), so *T*_*l*_ is the time of the first coalescence event just above time *u*. We need to condition on the coalescence point not being on edge *b*, and as a simplification we take

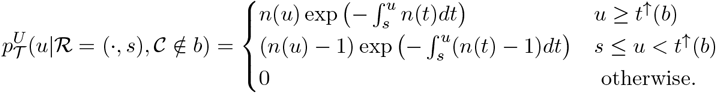

This essentially assumes SMC rather than SMC’ dynamics, since the two models differ only in that the latter allows the coalescence event to occur on the same branch as the recombination event, so this is a very close match for the conditional distribution (and simplifies our calculations). Thus, through a change of variable in (S1), for *t*^↑^(*b*)≠*T*_2_ and *s* − *t*^↑^(*b*) *< c <* 0,

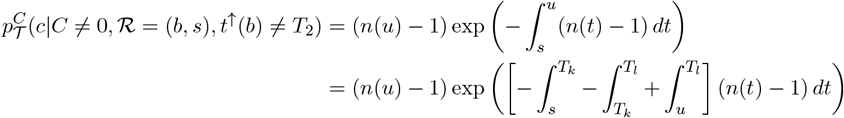

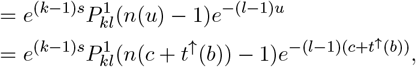

where

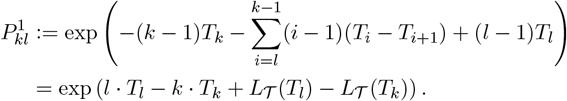

For *t*^↑^(*b*) = *T*_2_ and *t*^↓^(*b*) + *T*_3_ − 2*T*_2_ ≤ *c* ≤ 2(*T*_3_ − *T*_2_), similarly,

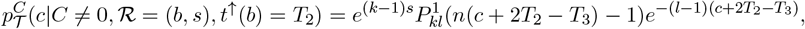

and for *t*^↑^(*b*) = *T*_2_ and 2(*T*_3_ − *T*_2_) *< c <* 0, since *l* = 2,

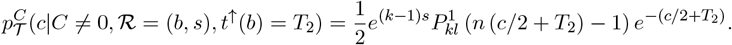

For 0 *< c* ≤ *T*_2_ − *t*^↑^(*b*),

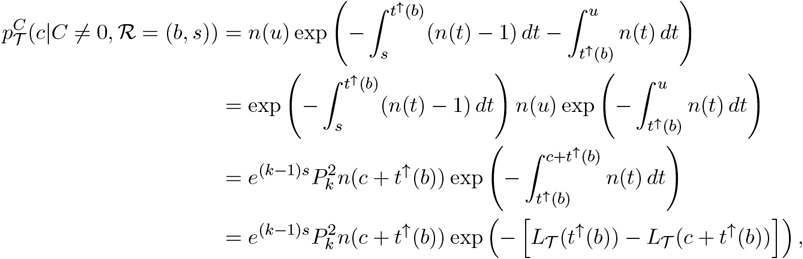

where

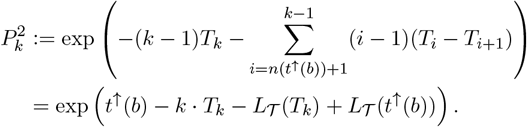

Finally, for *c > T*_2_ − *t*^↑^(*b*),

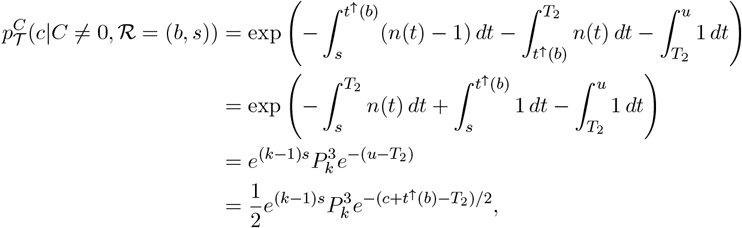

where

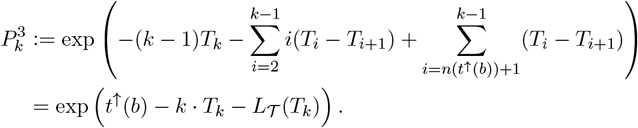

Note that

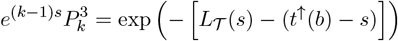

gives the probability that the coalescence event happens above *T*_2_. The conditional density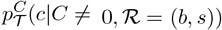 is thus

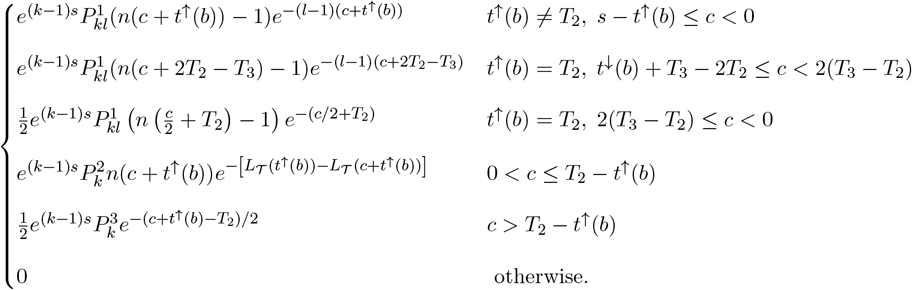

### S2.8.1 Density of change in total branch length

Marginalising out the position and time of the recombination point, we have

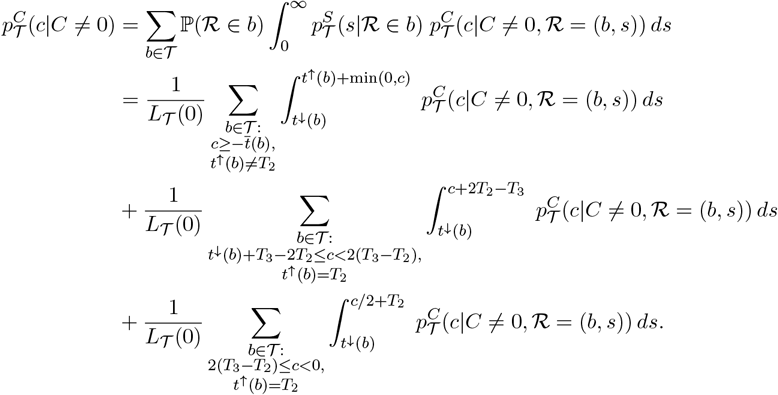

Let

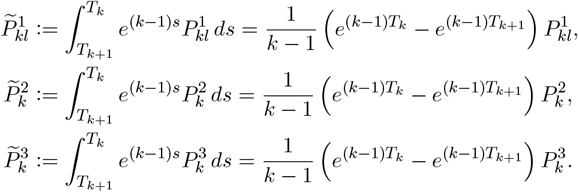

Then for *t*^↑^(*b*)≠ *T*_2_ and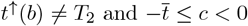,

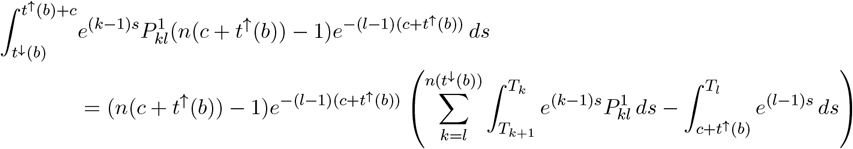

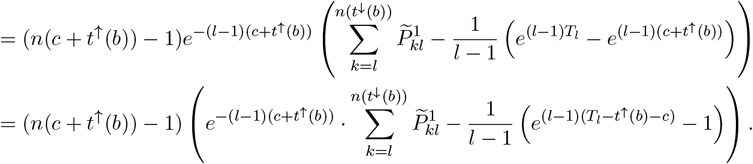

Similarly, for *t*^↑^(*b*) = *T*_2_ and *t*^↓^(*b*) + *T*_3_ − 2*T*_2_ ≤ *c <* 2(*T*_3_ − *T*_2_),

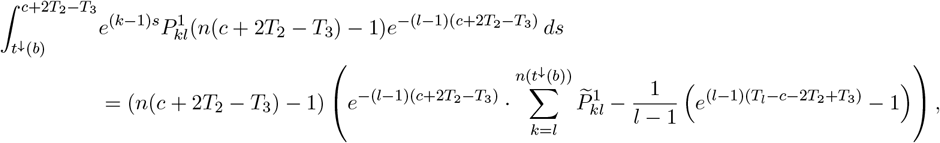

and for *t*^↑^(*b*) = *T*_2_ and 2(*T*_3_ − *T*_2_) ≤ *c <* 0,

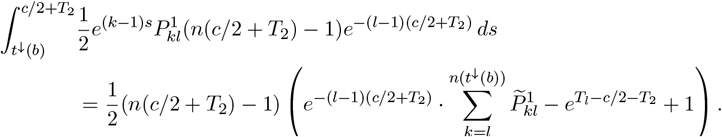

For 0 *< c* ≤ *T*_2_ − *t*^↑^(*b*),

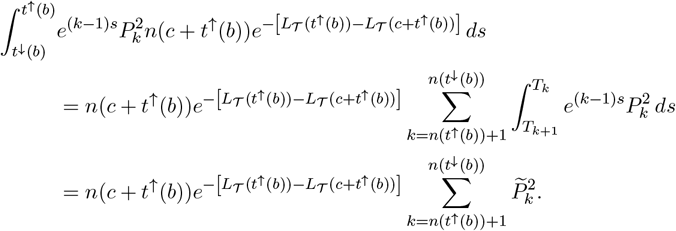

For *c* > *T*_2_ − *t*^↑^(*b*),

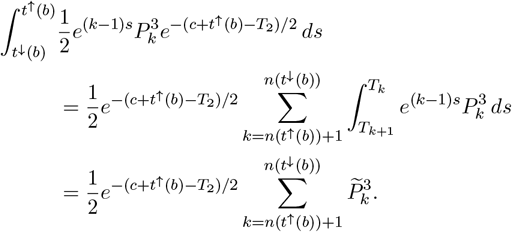

Thus,

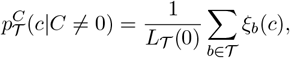

where *ξ*_*b*_(*c*) is given by

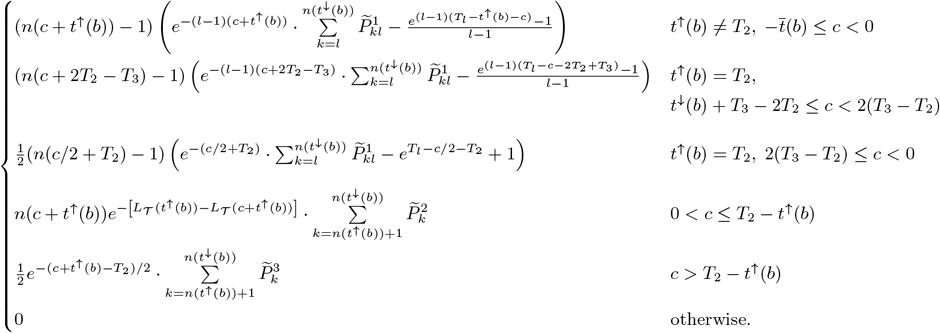

### S2.9 Proof of Proposition S1.7

The probability of a negative change in tree height, conditional on the location and time of the recombination point, is

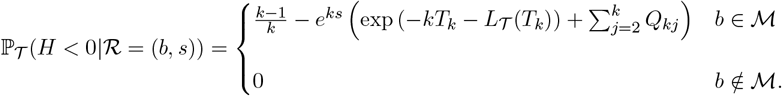

Integrating over the recombination time gives

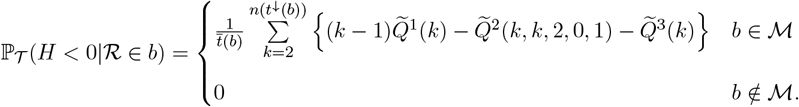

Summing over the edges and multiplying by the corresponding probability, the unconditional probability that the change in tree height is negative is thus

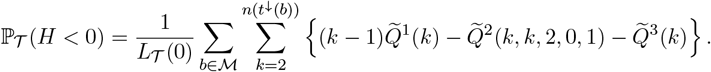

The probability of no change in tree height, conditional on the recombination point, is

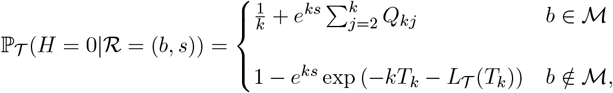

and

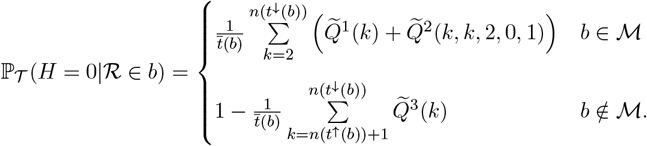

The probability of no change in tree height is thus

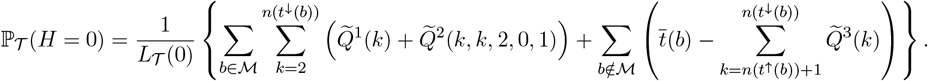

Finally, the probability of a positive change in height, conditional on the recombination point, is

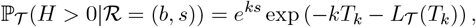

and

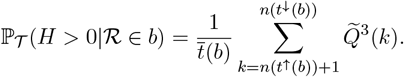

The probability that the change in tree height is positive is

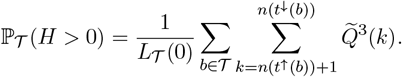

### S2.10 Proof of Theorem S1.2

Conditional on𝒯, the recombination point is chosen uniformly along the edges, so

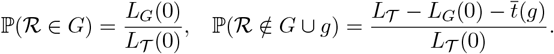

Conditional on the recombination point being in *G* and letting the clade MRCA time be *t*^↓^(*g*) = *T*_*m*_, the density of the recombination time is

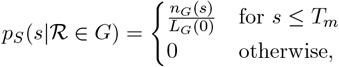

and similarly

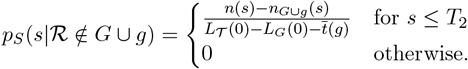

First conditioning on the recombination time,

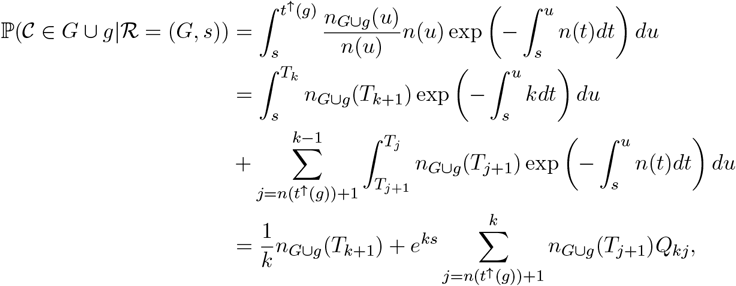

and so

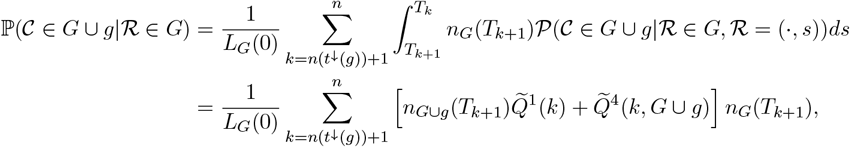

with

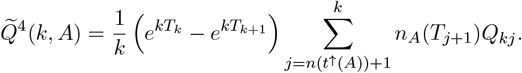

Similarly,

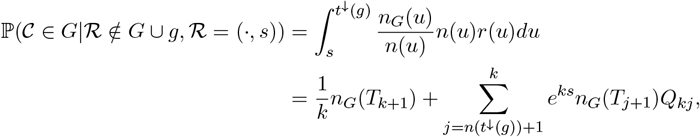

and

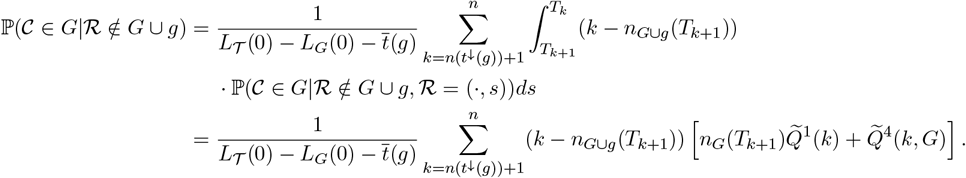

## S3 Supplementary Figures

**Figure S12:**
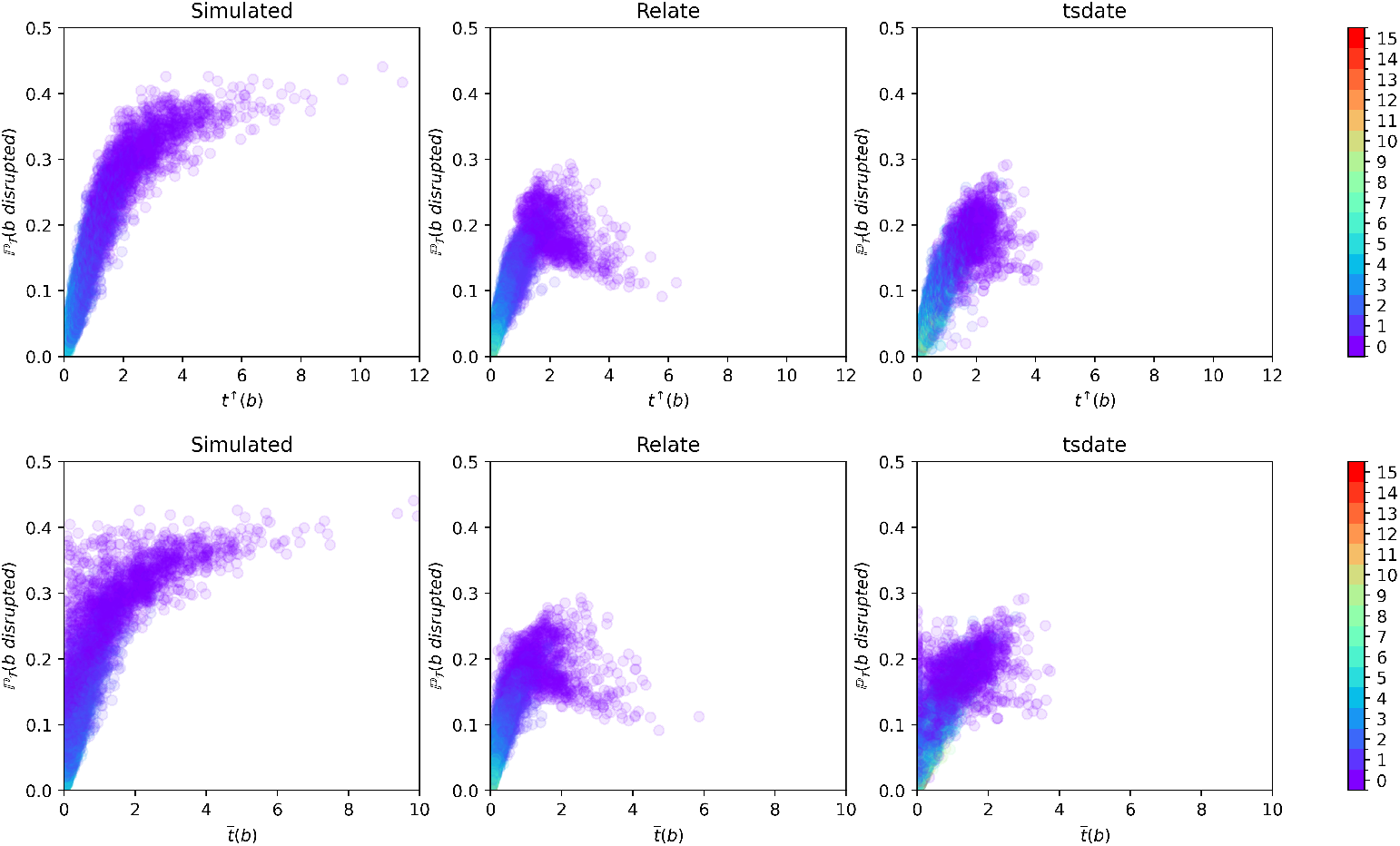
Same as Figure 2, but with *x*-axis showing the (un-normalised) time at the upper endpoint of each edge *t*^↑^(*b*) (top panel), and time-length of each edge *t*(*b*) (bottom panel).

**Figure S13:**
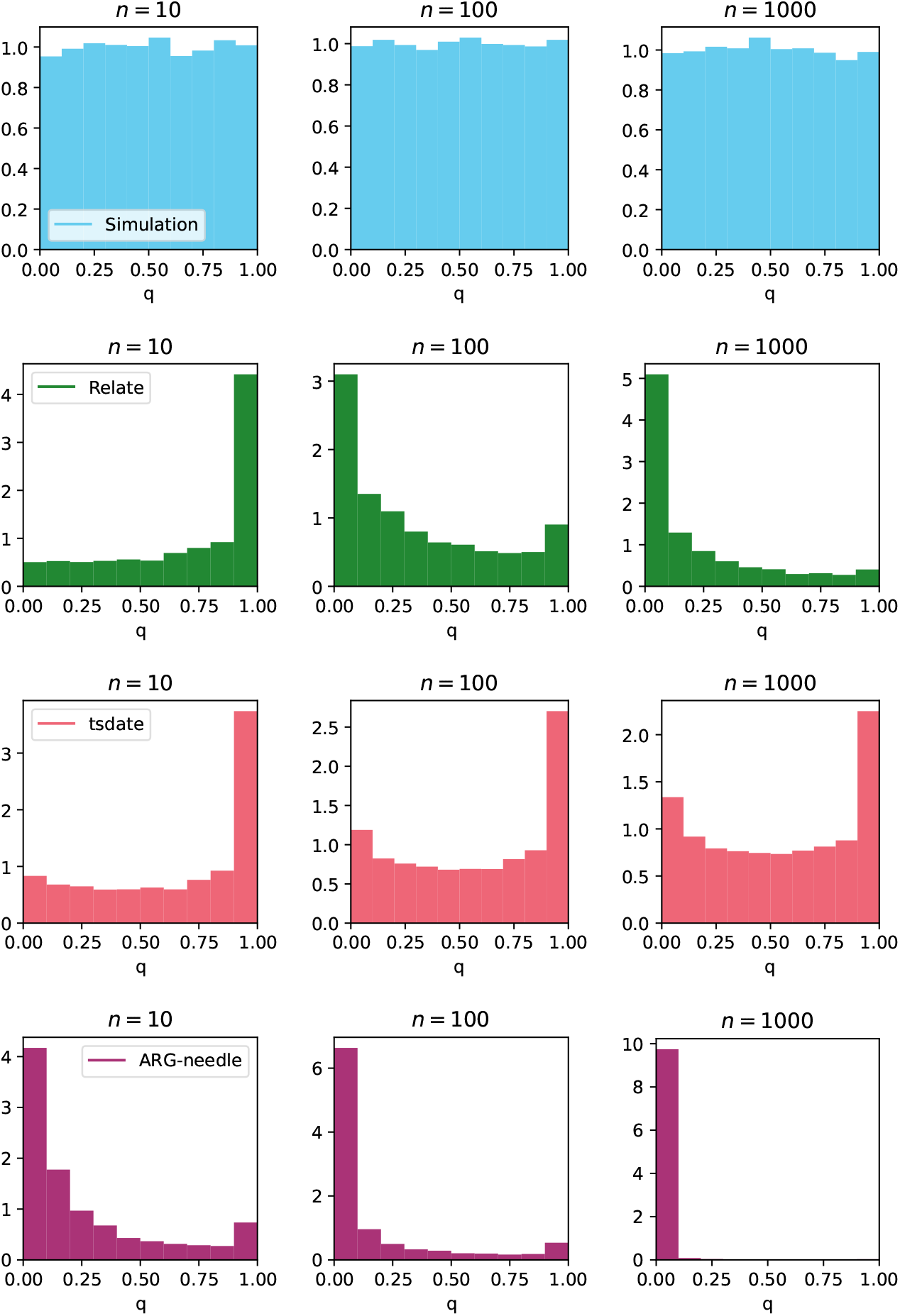
Histograms corresponding to Q-Q plots in Figure 3.

**Figure S14:**
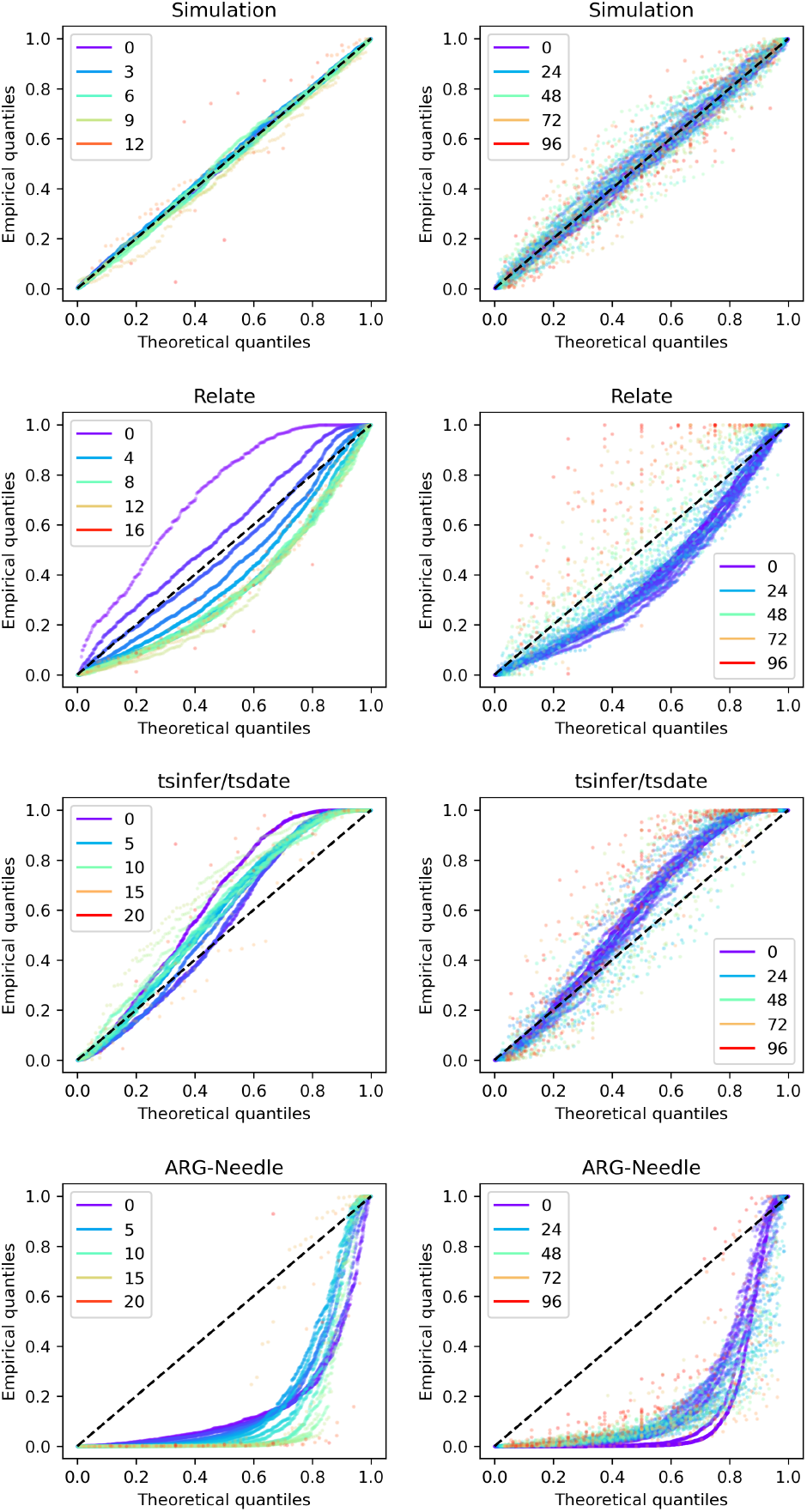
Q-Q plots split by clade size and depth. Q-Q plots using (S23) computed from an ARG simulated using dataset 1 parameters under the SMC’ (with *n* = 100). Left panels: split by depth of the edge (defined as the number of edges to the root of the tree); right panels: split by clade size (defined as the number of samples subtended by the edge). Dashed line: diagonal from (0, 0) to (1, 1). For the simulated ARGs, none of the corresponding K–S *p*-values for each group are significant using a 0.05 significance threshold.

**Figure S15:**
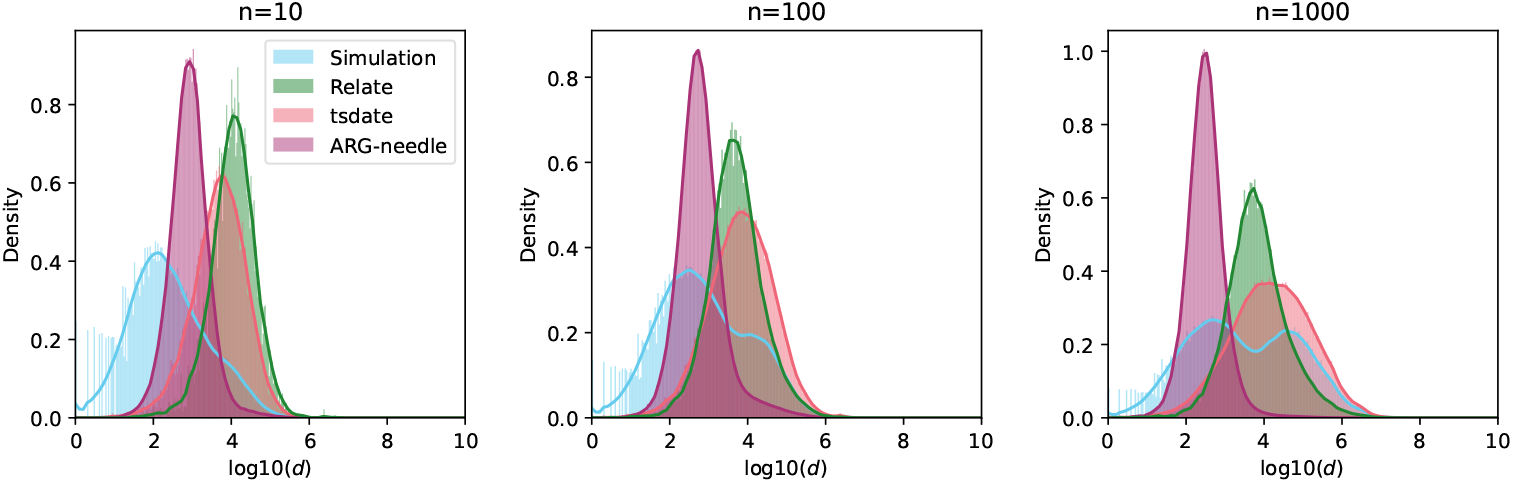
Histograms of observed edge span for simulated and reconstructed ARGs. Histograms of (observed) edge span, calculated as *d*(*b*) = *d*^→^(*b*) − *d*^←^(*b*) for each edge *b* in simulated and reconstructed ARGs (same as those in Figure 3). Note log scale on the *x*-axis.

**Figure S16:**
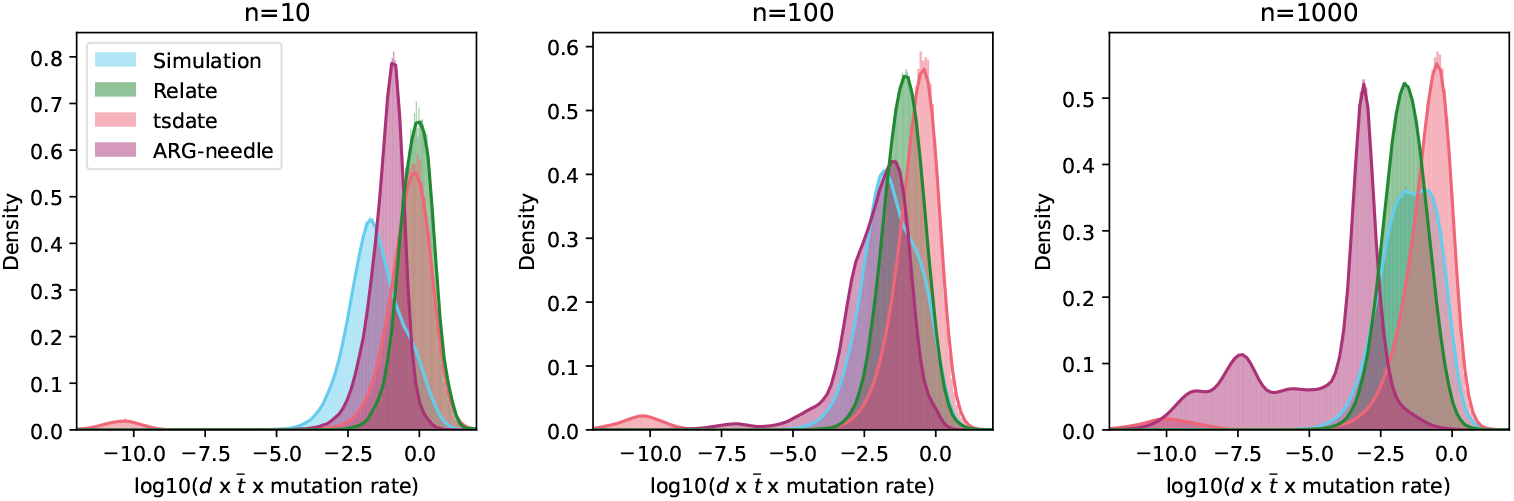
Histograms of expected number of mutations per edge for simulated and reconstructed ARGs. Histograms of (observed) expected number of mutations per edge, calculated as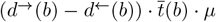 for each edge *b* in simulated and reconstructed ARGs (same as those in Figure 3). Note log scale on the *x*-axis.

**Figure S17:**
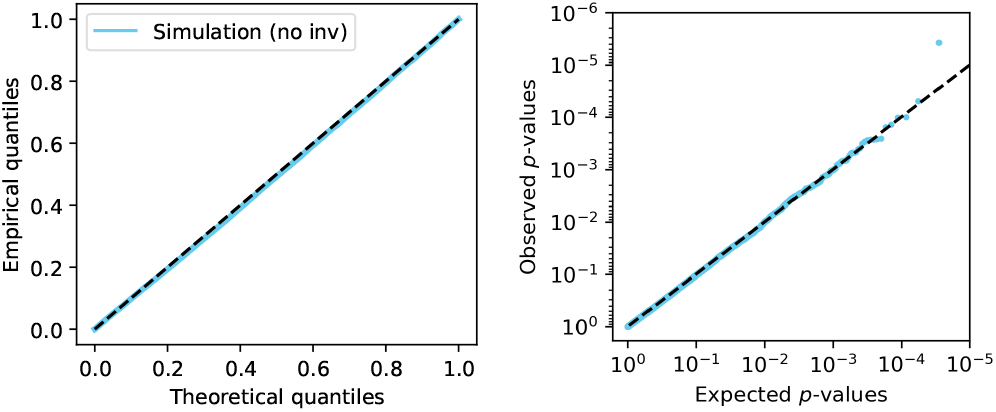
Q-Q and *p*-value plots (simulated ARG without inversion). Q-Q plot (left panel) and *p*-value plot (right panel) for ARG simulated using SLiM (without inversions and otherwise same parameters as in Section 4.6.2, main text). No clades have *p*-values below the Bonferroni-corrected significance threshold.

**Figure S18:**
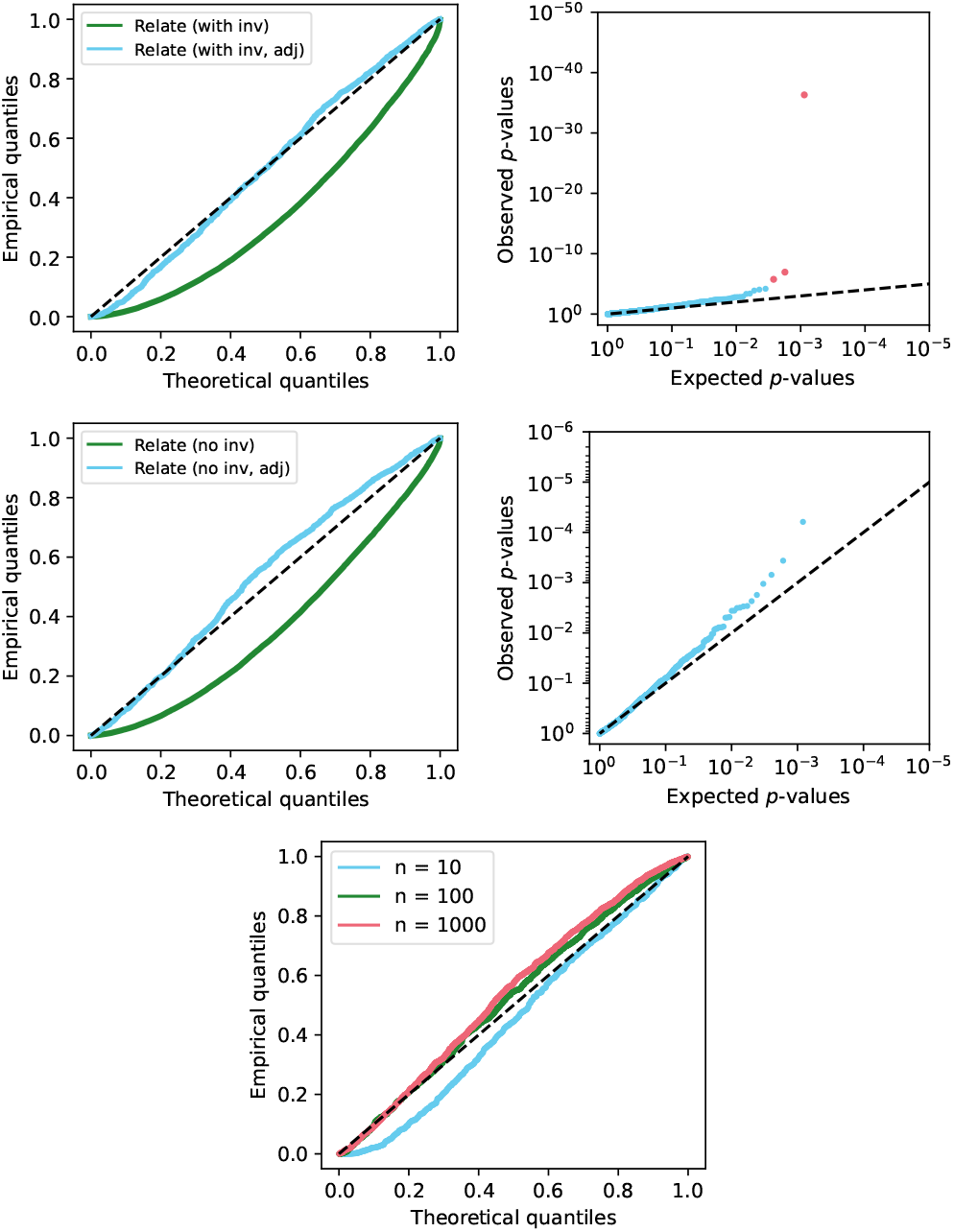
Q-Q and *p*-value plots (ARGs reconstructed using Relate). Top row: Q-Q plot (left panel) and *p*-value plot (right panel) for ARG reconstructed using Relate from data simulated using SLiM with one inversion under balancing selection (as described in Section 2.4.1). Blue (resp. green) points show values calculated after (resp. before) applying the adjustments described in Section S1.13.1; red points correspond to clades with *p*-value below the Bonferroni-corrected significance threshold. Middle and bottom rows: same using SLiM simulation without inversions (and otherwise same parameters); bottom panel shows QQ plot for Relate trees after the adjustments are applied, with varying sample sizes.

**Figure S19:**
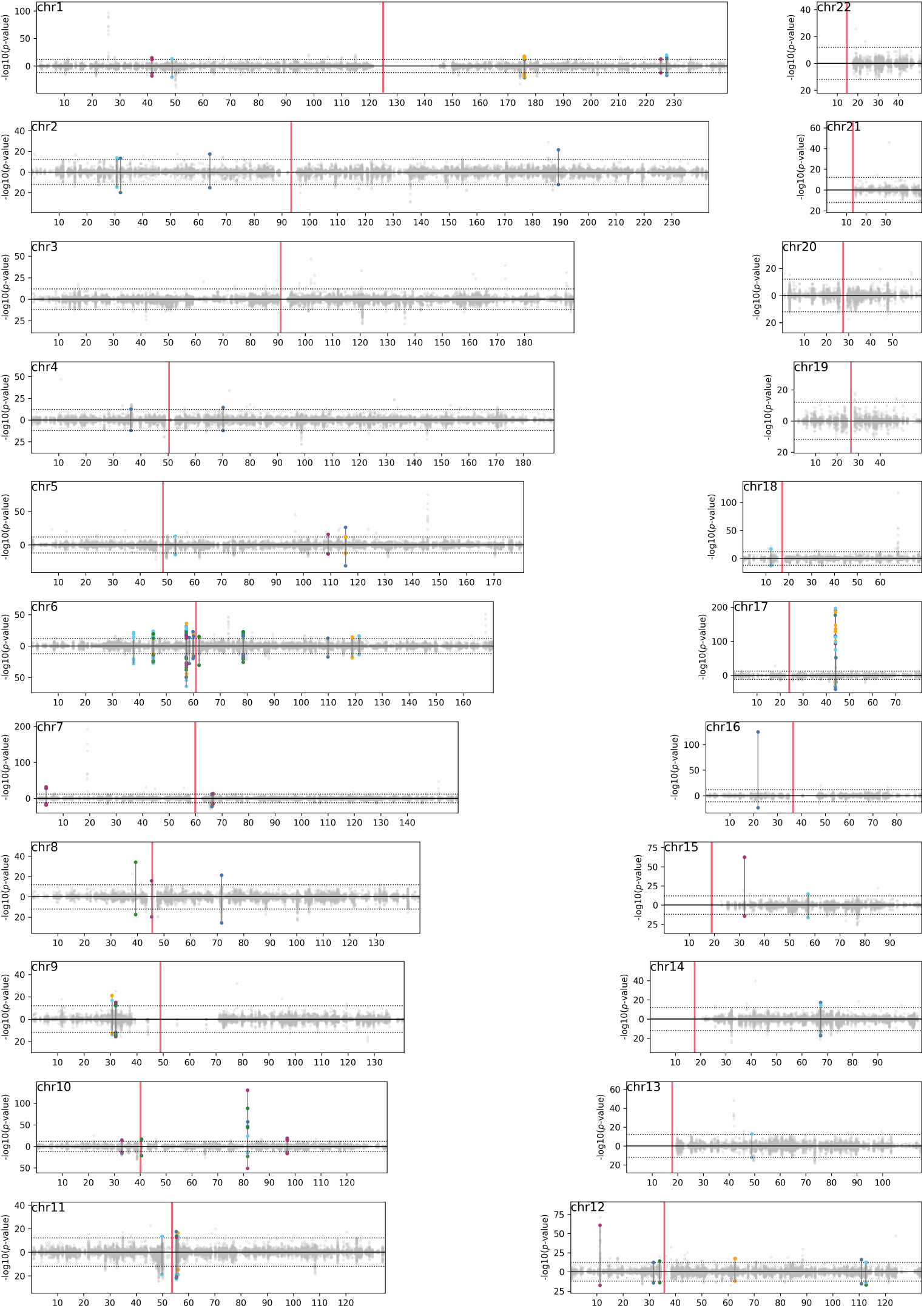
DoLoReS *p*-values for 1KGP ARG. Red vertical lines indicate positions of centromeres. See caption of Figure 6 (main text).

**Figure S20:**
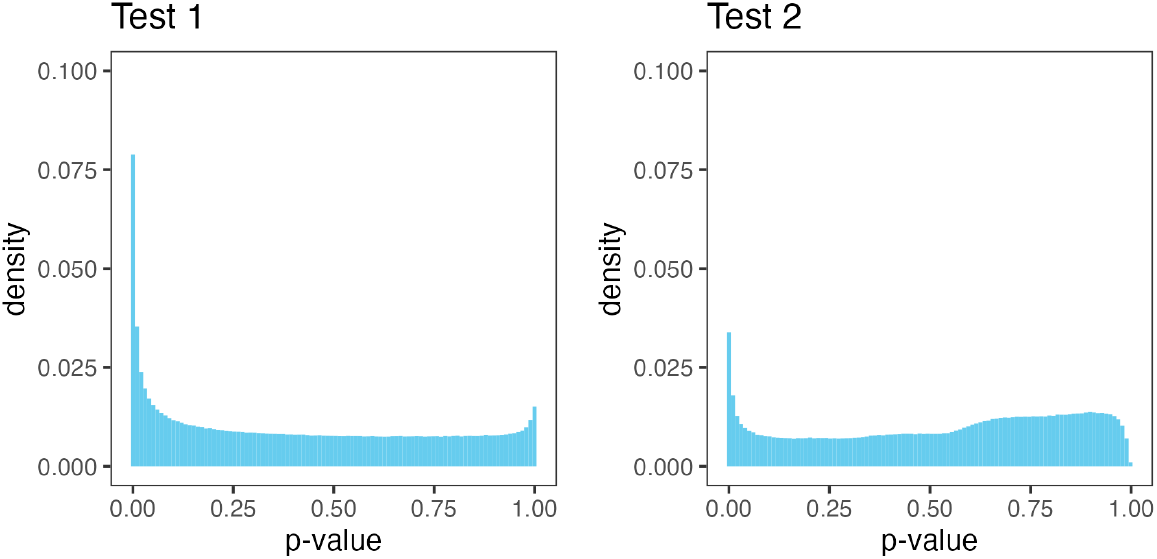
Histograms of *p*-values for Tests 1 and 2 for the 1KGP ARG (all populations combined, all clades with more than 2 mutations, at least 10 and at most *n* − 10 samples, spanning at least 2 local trees).

**Figure S21:**
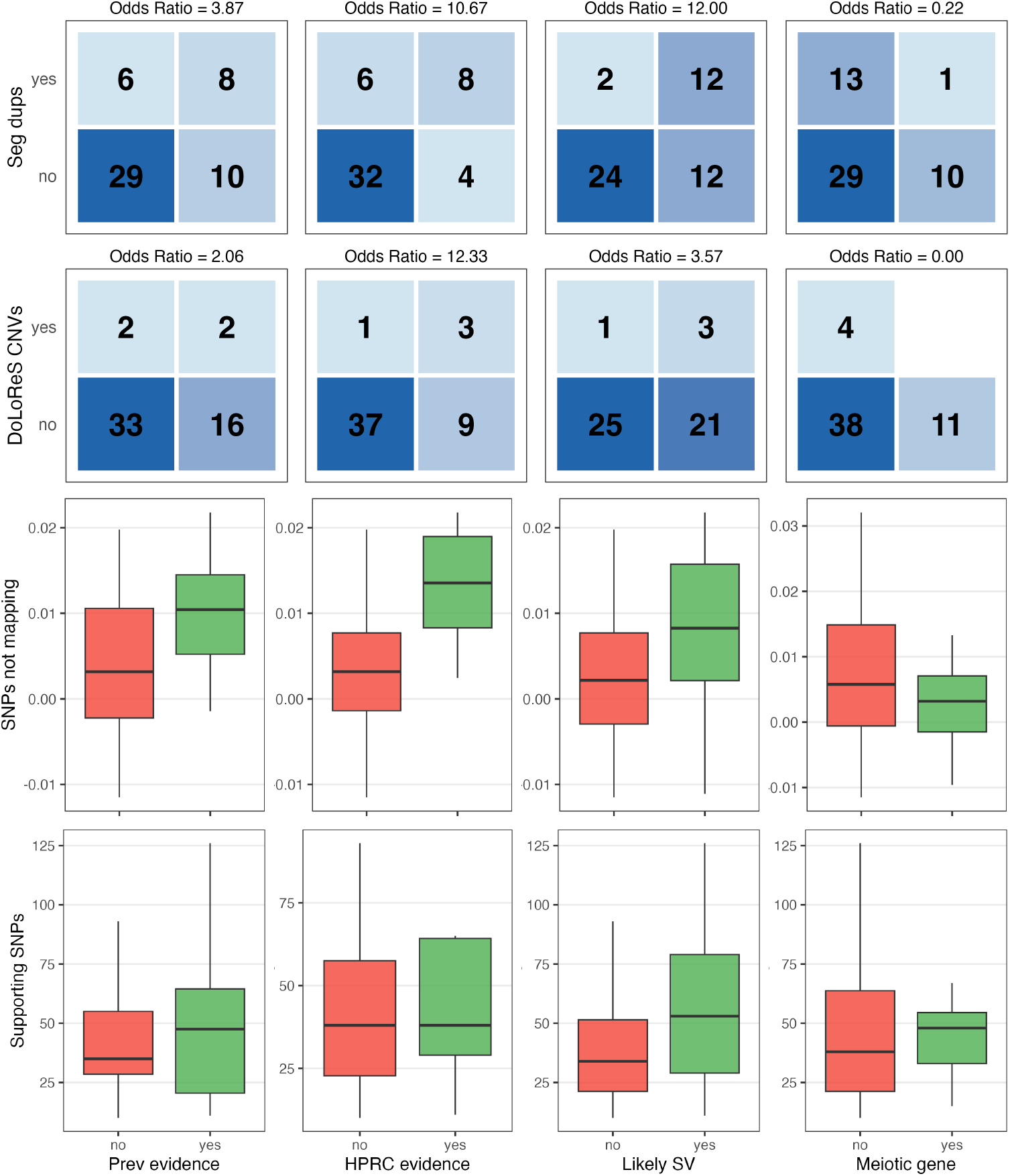
Some of the diagnostics that can be used to classify nature of recombination suppression in detected regions (given in full in Supplementary Table S1). Rows show the presence of direct or inverted segmental duplications (row 1), the detection of CNVs in the region by DoLoReS (row 2), the percentage point difference between the proportion of SNPs within the region that do not uniquely map to a branch of the local tree and the chromosome average proportion (row 3), the number of SNPs supporting the top significant clade (row 4). Columns show whether there is prior evidence from the literature for the detected region (column 1), whether our analysis of HPRC data shows evidence of SV (column 2), whether our assessment of the evidence together with other information (such as analysis of reads) points to the presence of an SV (column 3), and whether the region appears to exactly span a gene expressed in meiosis (column 4).

**Figure S22:**
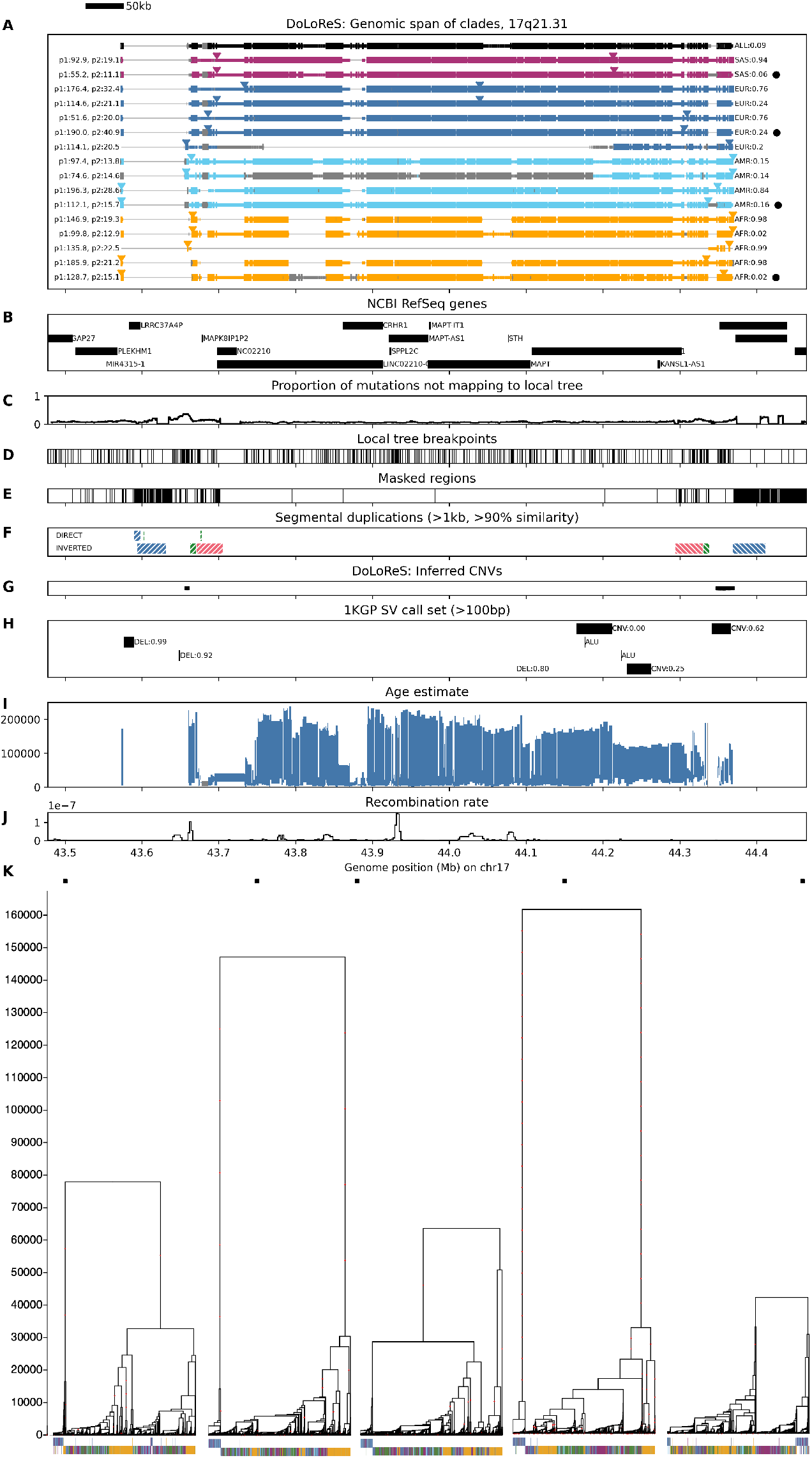
See caption of Figure 9 (main text). Age is estimated using the ARG subsetted to European populations.

**Figure S23:**
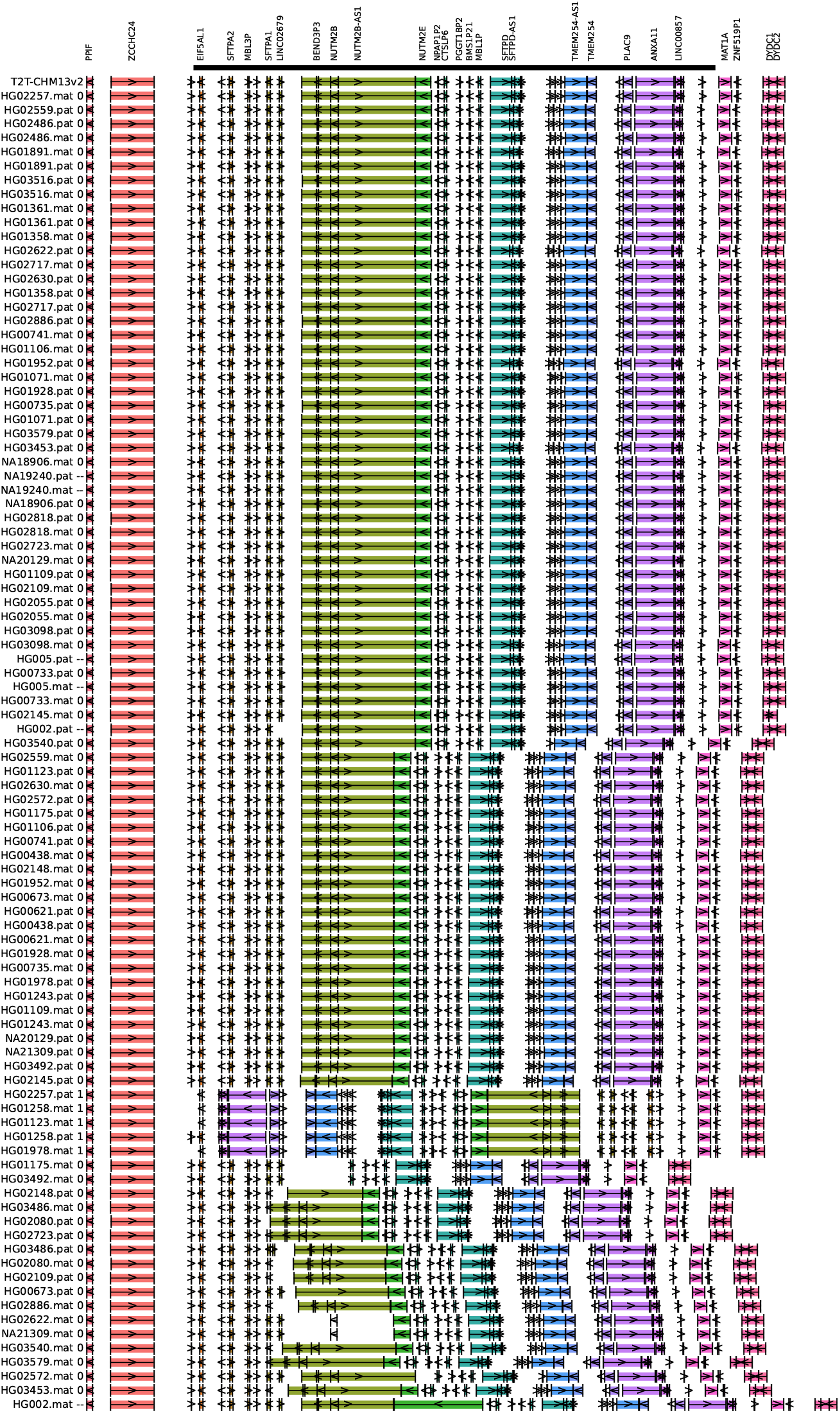
10q22.3 inversion region for HPRC data and T2T-CHM13 reference. Each row is a sequence, labelled by the individual ID, whether it corresponds to the maternal (mat) or paternal (pat) haplotype, and the predicted inversion status (0 for non-carrier, 1 for carrier). Genes are coloured uniquely, from orange to purple left-to-right for the un-inverted (ancestral) orientation. Black bar shows predicted span of inversion.

**Figure S24:**
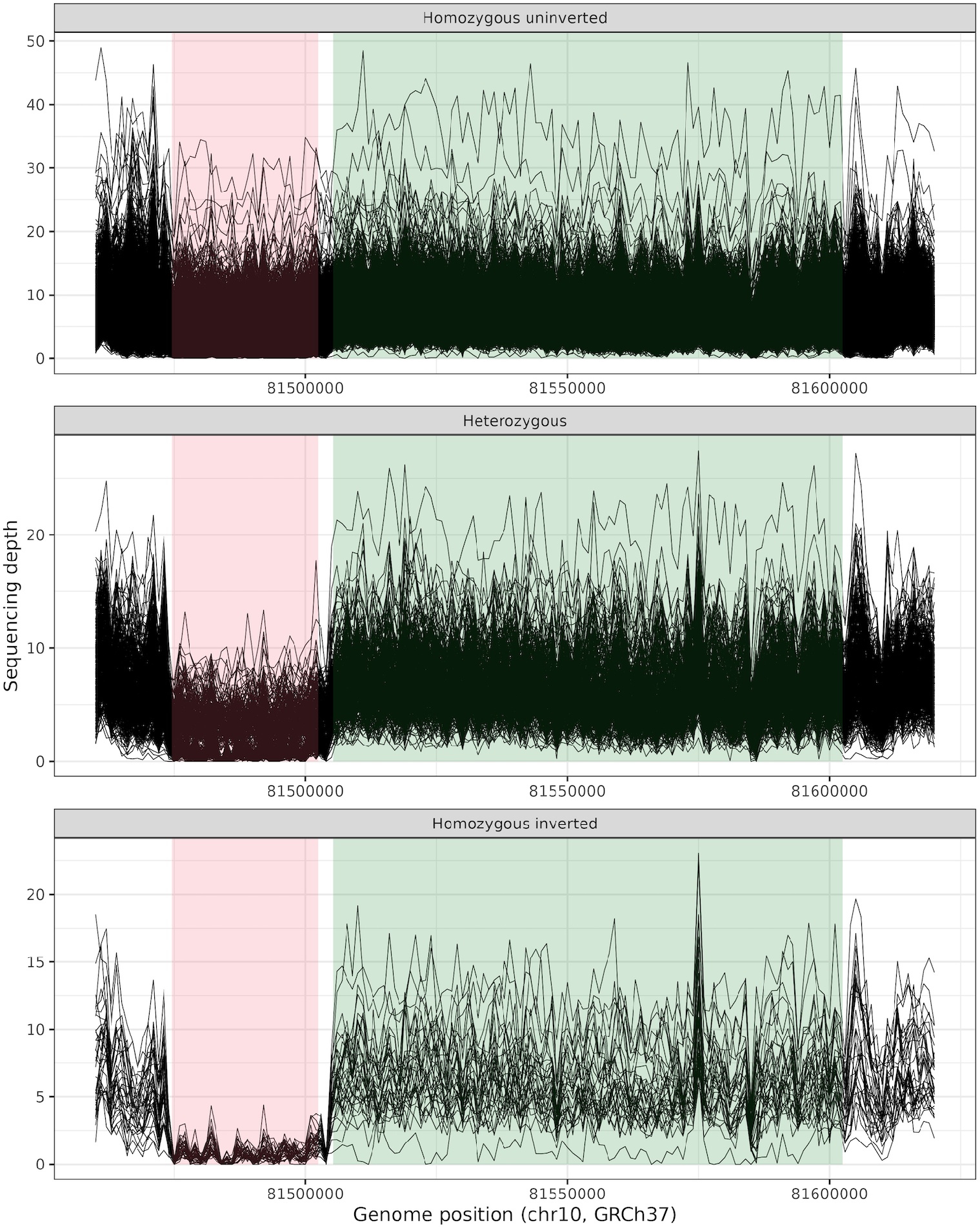
Sequencing read coverage on 10q22.3 around CNV1 (positions shown in red) and CNV2 (positions shown in green). Average coverage calculated in windows of 1000kb (each line corresponds to one individual) using 1KGP (Phase 3) low-coverage WGS GRCh37 data.

**Figure S25:**
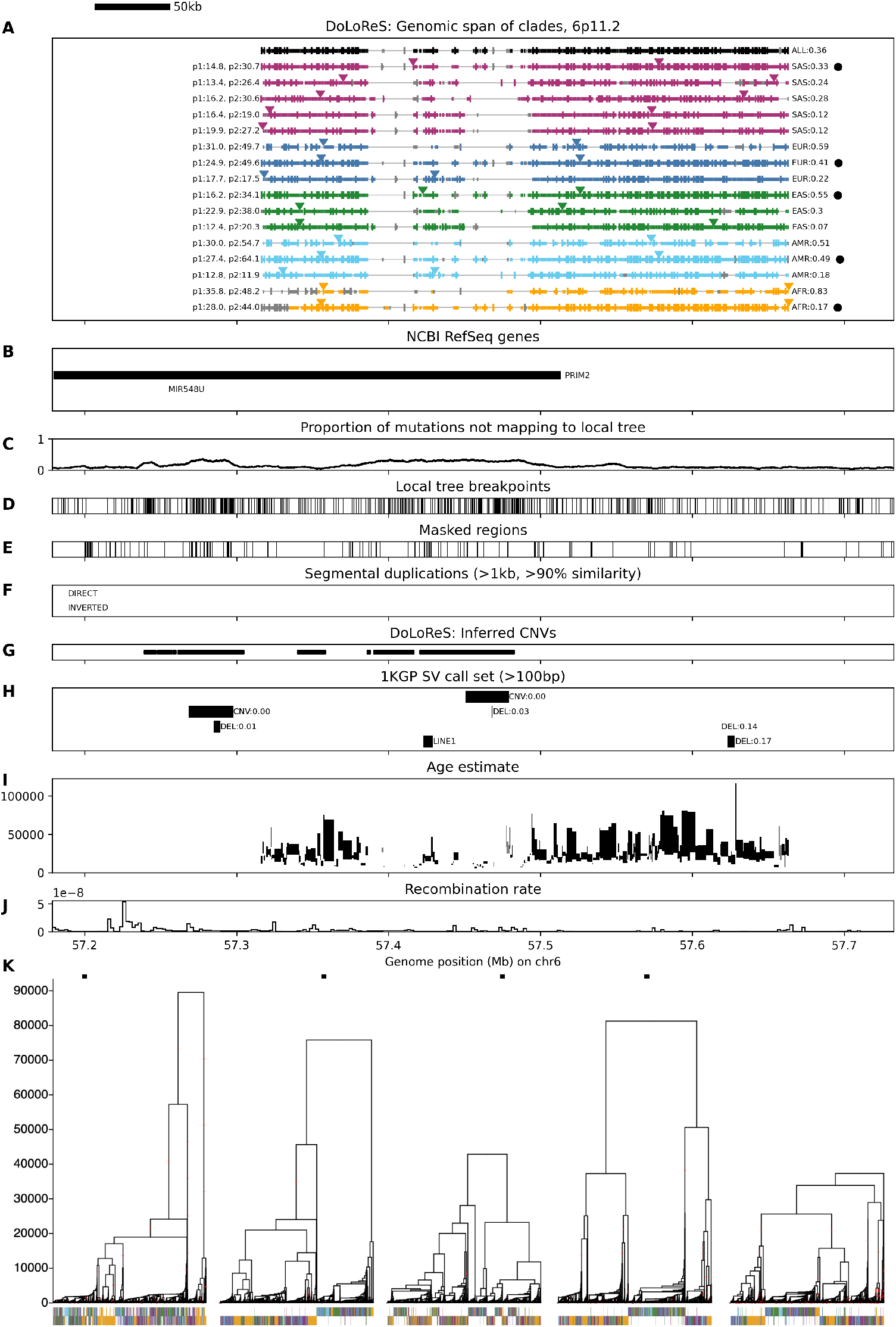
See caption of Figure 9 (main text). Age is estimated using the ARG for all populations.

**Figure S26:**
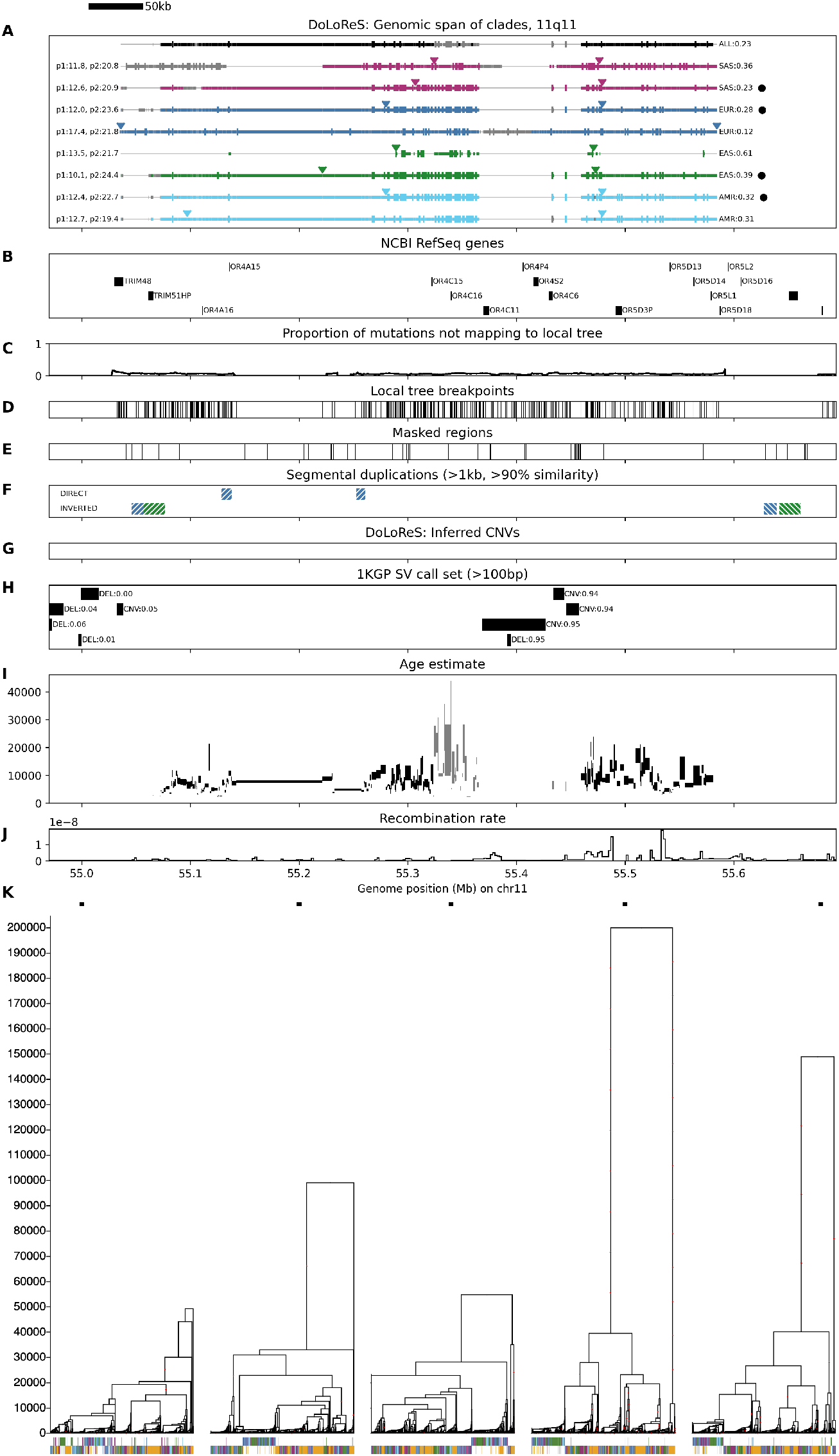
See caption of Figure 9 (main text). Age is estimated using the ARG for all populations.

**Figure S27:**
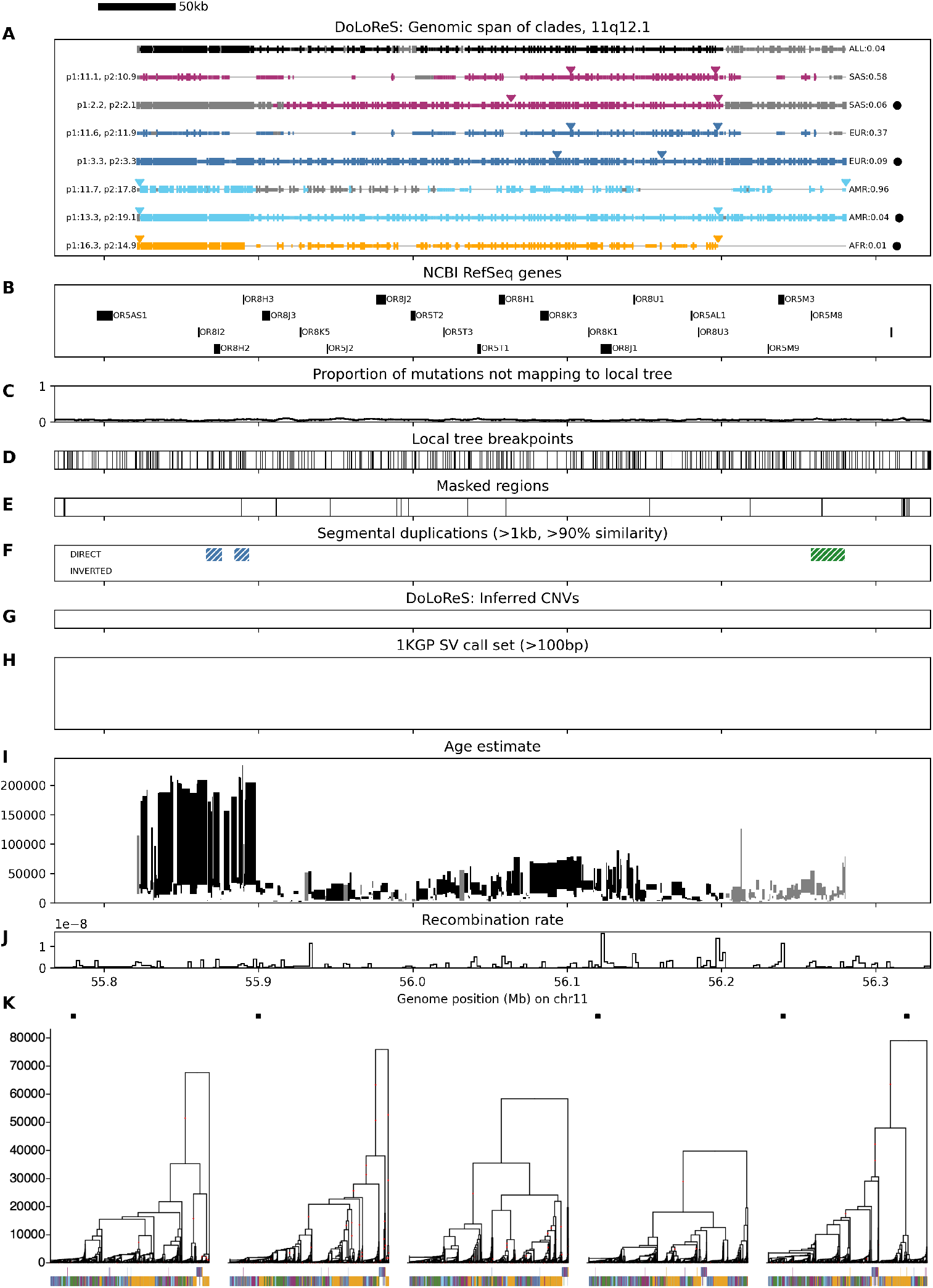
See caption of Figure 9 (main text). Age is estimated using the ARG for all populations.

**Figure S28:**
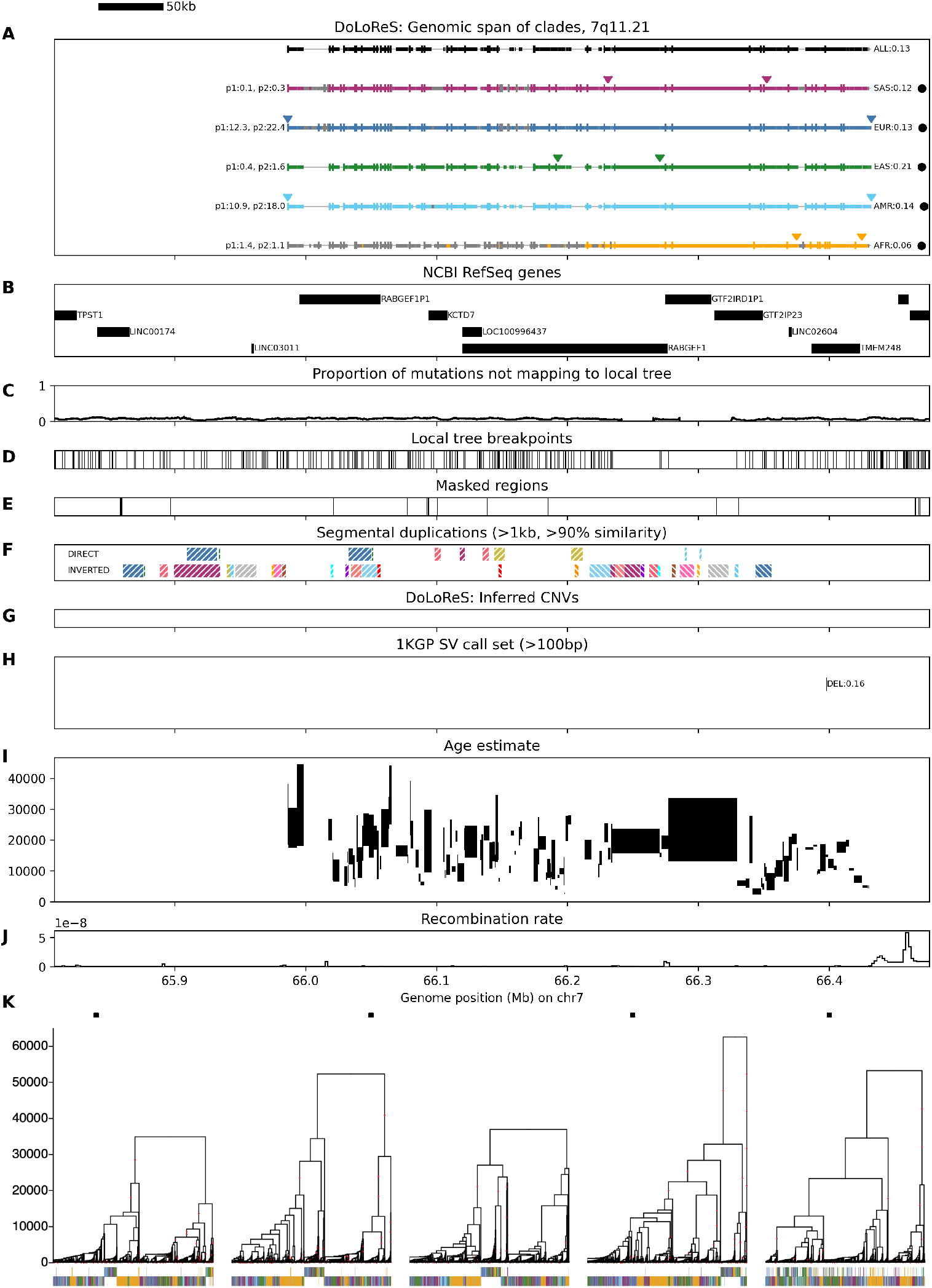
See caption of Figure 9 (main text). Age is estimated using the ARG for all populations.

